# Massively parallel assay of human splice variants reveals cis-regulatory drivers of disease-associated and cell type-specific splicing regulation

**DOI:** 10.1101/2025.10.12.681955

**Authors:** Samantha E. Koplik, Angela M Yu, Madelyn R. Shelby, Gabriel C. Fonseca, Charles M. Roco, Yue Zhang, Nicholas Bogard, Alex K. Sabo, Alexander B. Rosenberg, Johannes Linder, Georg Seelig

## Abstract

Splice-disrupting variants (SDVs) underlie many human diseases, yet systematic functional maps of their effects across cell types remain limited. We developed Cell-type Oriented Massively Parallel reporter Assay of Splicing Signatures (COMPASS) to measure splicing outcomes for 87,546 single and double variants across more than 1,700 genes in five human cell lines of diverse tissue origin. COMPASS targets disease relevant gene sets, including ACMG actionable genes and SFARI autism-associated genes, enabling systematic dissection of splicing impacts in health and disease. Our measurements reveal numerous SDVs, including ClinVar variants currently classified as variants of uncertain significance. Using prime editing, we validate variant effects in the genome for ClinVar variants in *BIN1*, a gene implicated in Alzheimer’s disease, cancer, and cardiac pathology. Benchmarking our MPRA against state-of-the-art predictive models shows that while these approaches capture broad variant effects, important gaps remain. Our data reveal putative RNA-binding protein motifs whose disruption drives splicing changes, providing mechanistic insight into variant impact. Finally, cross-cell line comparisons reveal a distinct subset of variants that drive cell type–specific splicing programs. This study delivers the largest cell type-resolved, base-resolution atlas of splicing variant effects to date, providing a resource to support variant reclassification in clinical genomics and aid in the selection of therapeutic targets.

**Graphical abstract:** 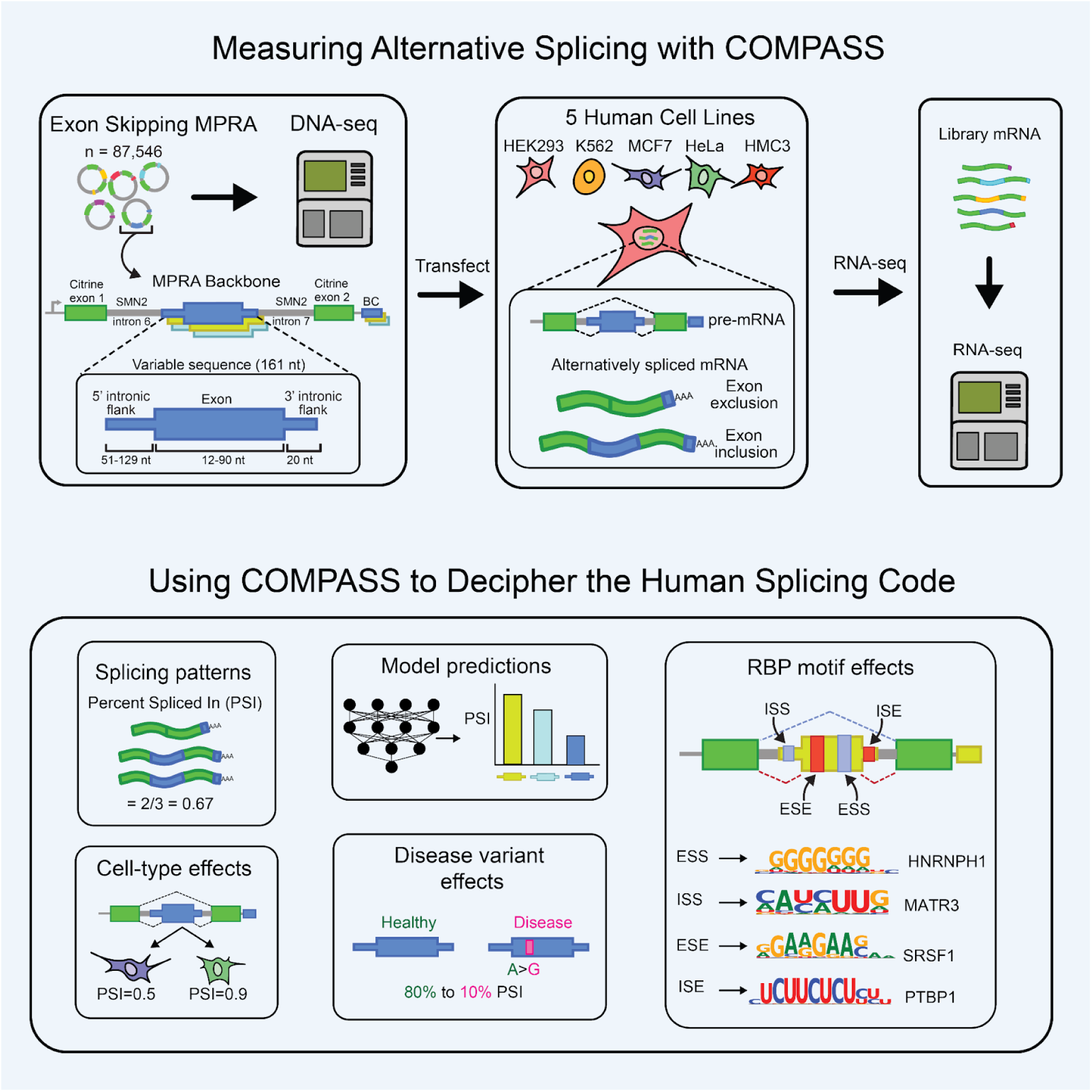

## Introduction

RNA splicing removes non-coding introns from pre-mRNA and joins protein-coding exons to enable translation of mRNA into functional proteins. An embedded *cis*-regulatory code determines when and where emerging transcripts are spliced. Splice donor (SD) and splice acceptor (SA) sequences define exon-intron boundaries and engage the core splicing machinery while intronic and exonic splice silencers and enhancers recruit RNA-binding proteins (RBPs) that modulate splice site usage. Multiple distinct splice isoforms often derive from the same gene through the usage of competing splice sites. Such alternative splicing (AS) greatly enhances the diversity of proteins in eukaryotes^1,2^. Heterogeneity in isoform usage arises from cell type differences in RBP expression, with nearly half of alternatively spliced isoforms displaying tissue-restricted patterns.^3,4,5^ However, quantifying splicing outcomes in heterogeneous tissues may be insufficient, as averaging across diverse cell types obscures cell type–specific effects.^6^

Variants that disrupt the *cis*-regulatory splicing code can change RNA and protein isoform ratios and identities, leading to a variety of human diseases.^7,8^ Splicing quantitative trait loci (sQTL) studies can link genetic variants to splicing outcomes at the population level, but they remain correlative and offer limited mechanistic insight into how individual variants alter splicing.^9–13^ The recent development of increasingly accurate splicing models has enabled considerable progress towards the identification of splice-disrupting variants (SDVs).^14–22^ Still, such models are not yet equipped to fully capture cell type-specific variant impact and, with few exceptions,^21,22^ are not designed to be easily interpretable and suggest mechanistic explanations for observed splicing phenotypes. Thus, while progress has been made toward deciphering the *cis*-regulatory splicing code, we still cannot always accurately predict and understand the impact of variants on splicing, beyond those that directly modify SD and SA sequences.

Synthetic splicing reporters are an attractive tool for studying sequence-to-function relationships, overcoming some of the limitations of traditional transcriptomics studies, enabling us to characterize less commonly observed sequences, including rare, pathogenic, or *de novo* variants. Prior work demonstrated the utility of splicing massively parallel reporter assays (MPRAs) to quantify variant impact on splicing for thousands of variants across hundreds or even thousands of exon contexts (MFASS, MapSY, Vex-seq).^23–25^ Most studies using synthetic reporters examine both references and single nucleotide variants (SNVs) in introns or exons to measure the variant’s impact on isoform abundance. Complementary research employed MPRAs with designed or fully randomized sequence elements, which can be cheaply scaled to hundreds of thousands of sequences to systematically characterize all possible *cis*-regulatory elements of a given length (e.g., hexamers) or even train predictive sequence function models.^15,21,26–33^ Alternatively, saturation mutagenesis of single disease-relevant exons has been used to assay a few hundred to more than a thousand SNVs (eg, *FAS* exon 6, *WT1* exon 5, *RON* exon 11, *CD19* exons 1–3, and *POU1F1* exon 2).^34–39^

Here, we build on this work and introduce Cell-type Oriented Massively Parallel reporter Assay of Splicing Signatures (COMPASS), scaling splicing MPRAs to nearly 90,000 variants across over 2,000 exons in five cell lines. In this context, cell lines provide a powerful system: they are homogeneous at a population level yet distinct enough from each other to resolve regulatory differences that would be masked in heterogeneous tissues, while still experimentally accessible for large-scale functional assays such as MPRAs. We test both single and double variants in the same sequence background to understand positional and combinatorial variant effects. By comparing the impact of *de novo* variants and variants of uncertain significance (VUS) with that of pathogenic variants in the same exons and introns, we can identify candidate disease-causing variants. Our dataset provides a foundation for further validation and potential improvement of computational splicing models; here, we assess leading ML models and retrain one such model on our dataset. Furthermore, we aim to explain variant effects by identifying putative RBP motifs and comparing the impact of variants that disrupt them across different sequence contexts. We identify sequences with cell type-specific splicing patterns and propose regulators for these behaviors. Together, our work provides an unprecedented resource for understanding how sequence variation shapes splicing outcomes both broadly and in a cell type-specific manner.

## Results

### MPRA design and variant selection

We developed COMPASS, an exon skipping MPRA of short human exons and their flanking introns (**Figure 1A**). We selected exons from the human genome 90 nt or shorter; we further filtered their nucleotide sequence length to be a multiple of three to avoid destabilizing reporter transcripts through nonsense mediated decay. Exons matching these requirements were included even if they are naturally constitutively spliced, in order to broadly learn the relationship between nucleotide sequence and exon inclusion. For each exon, we also included 20 nt of the corresponding downstream intron and a variable stretch of the upstream intron such that the total length of the assayed region is 161 bp. The variable regions are flanked by constant sequences derived from SMN2 introns 6 and 7 to ensure the exon can be recognized by the splicing machinery.^15,40,41^ Random 3’UTR barcodes were associated with the variable cassette exon and introns through DNA sequencing. The barcode enables mapping of each transcript to a reporter gene of origin, even when the entire variable intronic and exonic region is excised.

**Figure 1.**
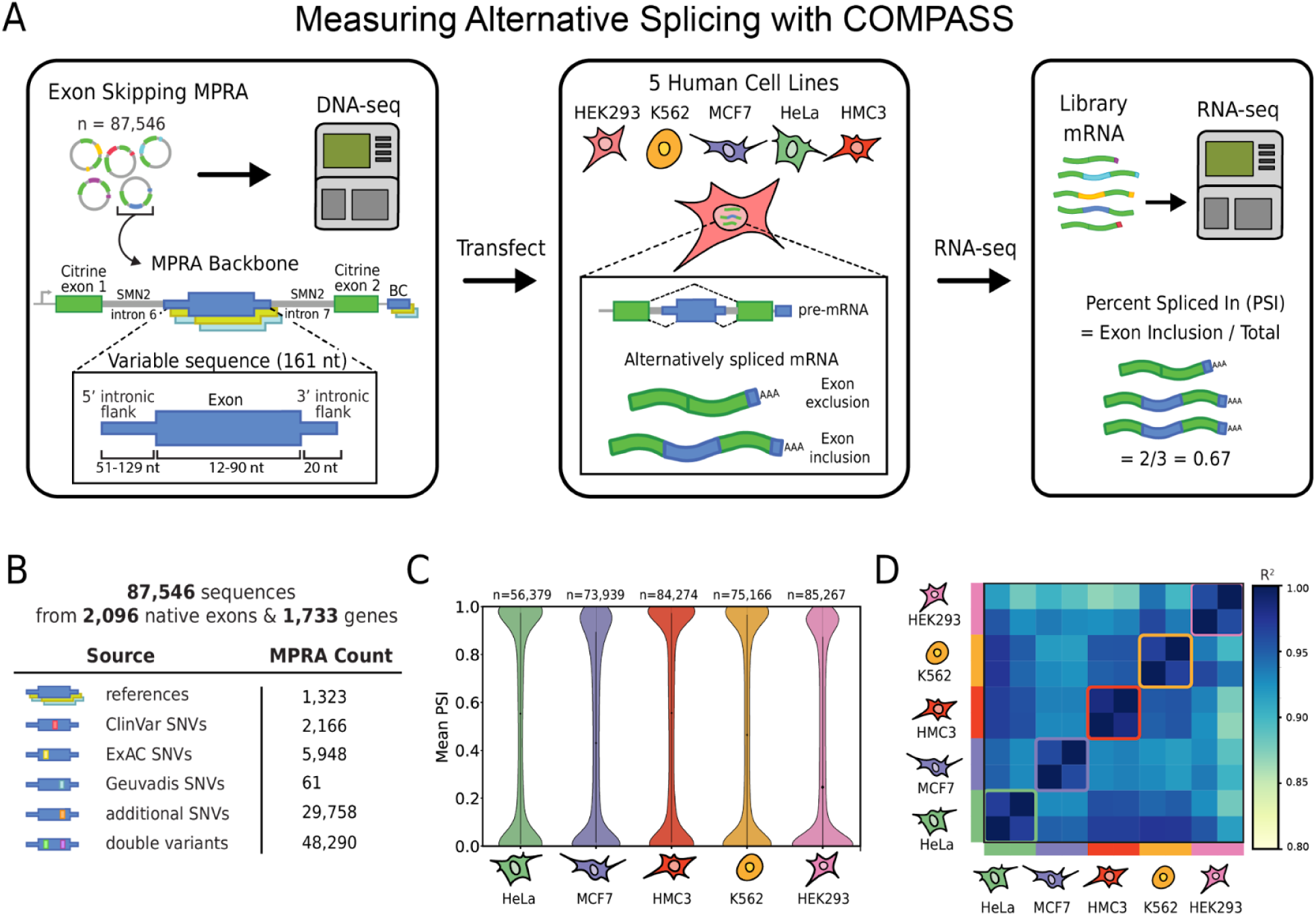
An exon skipping MPRA was performed on over 87,000 sequences in 5 human cell lines. (A) A workflow diagram of COMPASS showing the structure of the splicing reporter containing variable and constant regions, which are transfected into 5 human cell lines (HEK293, HeLa, K562, MCF7, and HMC3), with resulting mRNA sequences via RNA-seq. (B) The breakdown of the sources from which the COMPASS sequences containing short exons and intronic flanks were derived. (C) Measured PSI distribution across library sequences in 5 cell lines. (D) Pearson R^2^ correlation of the MPRA from 5 cell lines (HEK293, HeLa, K562, MCF7, and HMC3).

We mined publicly available databases to identify disease-associated and common human variants that fall within these introns and exons (**Figure 1B**). COMPASS contains 1,323 references, 61 variants from Geuvadis; 5,948 from the Exome Aggregation Consortium (ExAC); 2,166 from ClinVar; and an additional 29,758 SNVs and 48,290 double variants randomly targeted to introns and exons that contain at least one ClinVar variant.^42–44^ In total, the library contains 87,546 sequences spanning 2,096 native human exons from 1,733 genes.

### Mapping of splice junctions and quantification of isoform abundance

We delivered the pooled plasmid library to five different human cell lines: the ENCODE reference cell lines HEK293 (embryonic kidney), K562 (leukemia), HeLa (Adenocarcinoma, uterus), MCF-7 (Adenocarcinoma, breast) and, additionally, HMC3 (microglia) to capture microglia-specific splicing^45,46^ We collected and sequenced the library-specific mRNA, and bioinformatically determined the splicing profiles of each sequence as outlined in **Methods**, with the aid of the barcode sequence in the 3′ UTR. Next, we calculate Percent Spliced In (PSI) of expected splice junctions, defined as the number of RNA molecules containing the expected exon divided by the total number of RNA molecules with the corresponding barcode (**Figure 1A**). PSI is reported as a fraction on the 0–1 scale rather than an actual percentage, consistent with prior literature.^1,17,18,47^ A majority of reporters exhibit either dominant exon inclusion or exclusion, with relatively fewer reporters exhibiting moderate inclusion levels consistent with alternative splicing (**Figure 1C**). Splicing outcomes between cell lines are highly correlated, though less than replicates from the same cell line **(Figure 1D; Figure S1A**). Literature values (**Figure S2A**) and gel-based validation experiments (**Figure S2B**) showed good agreement with sequencing-based measurements for a set of control reporters. We also observed strong concordance with MFASS for a set of overlapping variants (**Figure S2C**) and our own independent low-throughput reporter assay (**Figure S2D**), supporting the reliability of our measurements.

### Quantifying and predicting variant impact on splicing

Next, we turned to characterizing the impact of single nucleotide variants (**Figure 2A**). We found 87,547 sequences with measurements in at least one cell type. Of these, 2,559 lacked a matched reference, precluding calculation of variant-associated changes. In total, 83,852 variants remained with an associated reference, allowing calculation of ΔPSI, defined as the difference between the variant and reference PSI. Replicates were highly correlated not only at the PSI level but also at the ΔPSI level (**Figure S1A**). A variant that increases exon inclusion is associated with a positive ΔPSI. While easy to interpret, ΔPSI is highly sensitive to the reference PSI value (**Figure 2B**). Measured ΔPSI values tend to be small if the reference exon is fully spliced in or out, as in these cases, even a large perturbation may not be enough to dislodge the exon from its preferred state. Conversely, for alternatively spliced exons where the competing SDs and SAs are relatively balanced, comparably weaker perturbations can result in large PSI changes. This dependence on the starting inclusion level means that a large ΔPSI in the reporter context may not correspond to a similarly large ΔPSI in the native gene context. Although the splice regulatory signals in the central exon and near intronic context are unchanged, the competing distal splice signals, i.e., those used when the exon of interest is spliced out, can vary between reporter and native context.

**Figure 2.**
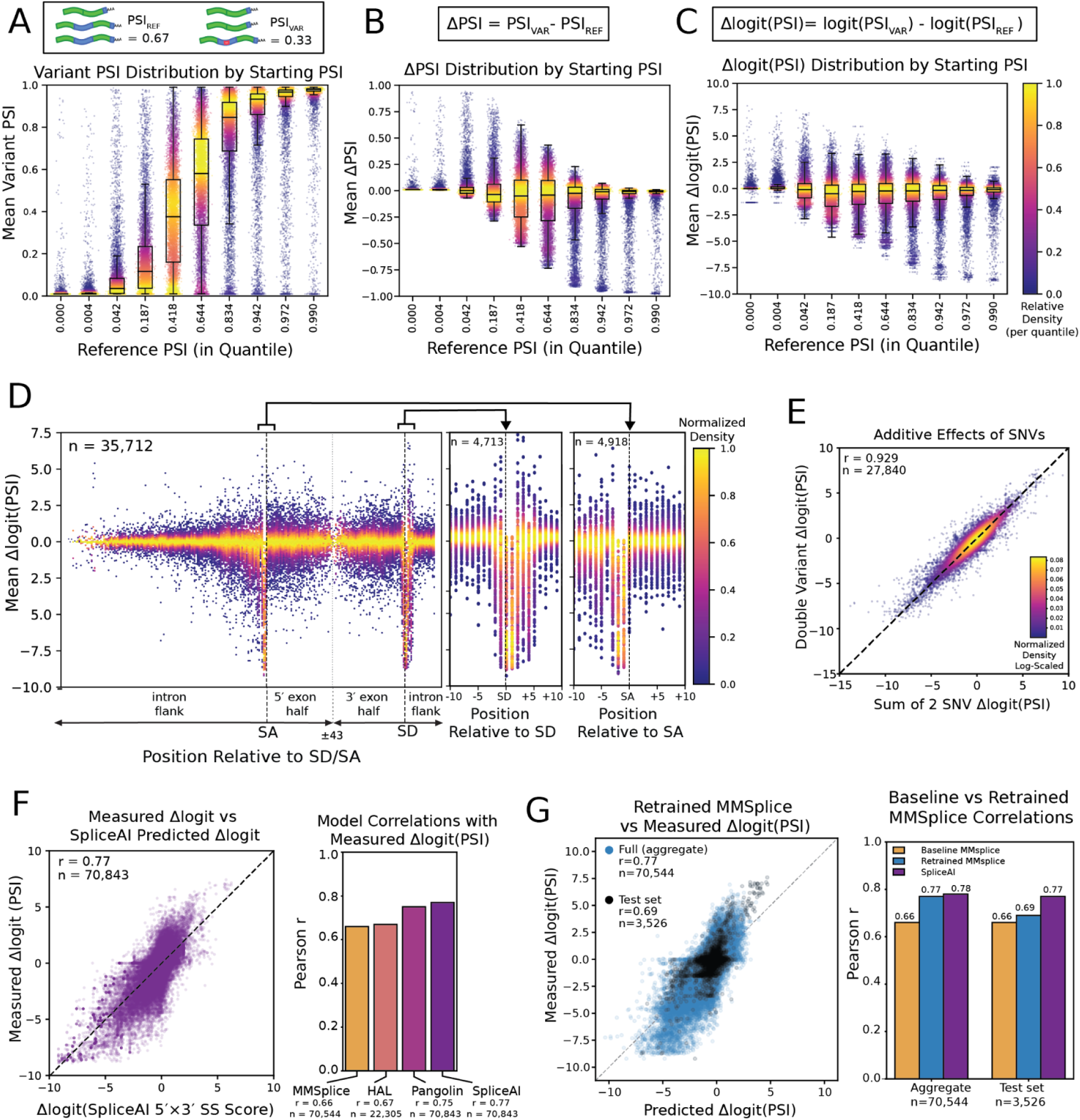
Global splicing effects of single and double variants. (A) Variant PSI values grouped into ten quantiles based on the reference (starting) PSI, showing the distribution of single and double variants within each quantile. (B) ΔPSI values grouped into the same reference PSI quantiles, highlighting how variant-induced splicing changes scale with the baseline inclusion level. (C) Δlogit(PSI) values grouped by reference PSI quantiles, showing that the logit transformation attenuates the dependence of variant effects on the starting PSI. In all panels, densities are normalized within quantiles to facilitate comparison of relative effects. (D) Distribution of Δlogit(PSI) as a function of SNV position relative to the SA or SD (n = 35,712). Densities are normalized per position and show an enrichment of large-magnitude effects in proximity to the SD and SA. Insets to the right highlight the regions spanning 10 nucleotides upstream and downstream of each splice site. (E) Additive effects of SNVs, comparing the summed Δlogit(PSI) values of two individual variants with the measured Δlogit(PSI) of the corresponding double variant carrying both changes (n = 27,840, Pearson r = 0.929). (F) (left) Predictive performance of computational models, comparing experimental Δlogit(PSI) with SpliceAI predictions. SpliceAI achieved the strongest performance (n = 70,843, Pearson r = 0.77), capturing the greatest number of variants and showing the best agreement with measured effects. (right) Pangolin performed to a similar level (n = 70,843 Pearson r = 0.75) and captured the same number of variants as SpliceAI. For comparison, MMSplice, which is limited to variants within 100 nucleotides of the variable exon, achieved Pearson r = 0.66 (n = 70,544), and HAL, which is limited to exonic variants, achieved Pearson r = 0.67 (n = 22,305). (G) Retraining MMSplice on our dataset improved performance to Pearson r = 0.69 on a held-out test set of 3,526 variants, demonstrating increased concordance with experimental measurements relative to the baseline MMsplice model. When evaluated across the full aggregate dataset (training, validation, and test combined), the retrained model reached Pearson r = 0.77, comparable to SpliceAI on the same sequences (r = 0.78) and markedly higher than the baseline MMSplice model (r = 0.66). All other correlation plots are shown in **Figure S2.**

For most of our analysis, we use the delta log odds ratio (Δlogit) as a more robust and generalizable metric for variant impact (**Figure 2C**).^15,17,18,24,48–50^ Variants that increase exon inclusion have a positive Δlogit(PSI). These Δlogit(PSI) values are less sensitive to starting PSI and are more evenly distributed (**Figure 2C**). To define splice-disrupting variants (SDVs), we considered any variant with |Δlogit(PSI)| ≥ 1 to have a significant effect on splicing. Prior MPRA studies defined similar thresholds on the log_2_ scale; here we use the equivalent cutoff on the natural log scale to define SDVs, which corresponds to roughly a 0.25 change in PSI at mid-range inclusion. While the Δlogit(PSI) is thus a useful measure of variant impact, practical applications ultimately require knowledge of the variant ΔPSI in the native gene context. These values can be estimated with a biophysical model given knowledge of the reference PSI obtained from RNA sequencing data and the corresponding variant logit measured in the reporter construct.^48^ Still, we note that Δlogit(PSI) is also an imperfect metric, as it is highly sensitive to sequencing depth, as discussed in the **Methods**. To mitigate this and avoid artifacts introduced by logit transformation, PSI values were clipped prior to transformation, as is standard in Δlogit(PSI) analyses.^17,18,49,51,52^ A clipping interval of [0.01, 0.99] was selected empirically as outlined in the **Methods** and is consistent with FRASER.^49^ Without clipping, PSI estimates become strongly dependent on read depth, particularly when the reference and variant used in the calculation are not sequenced to comparable coverage.

To examine how positional context influences variant effects, we plotted the Δlogit(PSI) of each variant at its position relative to the nearest splice site. This visualization highlights that variants disrupting SDs and SAs tend to have the most pronounced and most negative effect (**Figure 2D**). However, variants with large negative or positive effect sizes can be found anywhere in the assayed introns and exons, consistent with a *cis*-regulatory code that extends beyond core elements. Varying exon length means that the number of variants per position decreases with increasing distance from the splice sites (**Figure 2D**). Because we sequenced across the variable exon and both flanking junctions, we could also detect unexpected splice isoforms, often arising from variants that create novel donor or acceptor sites (**Figure S2E**).

We next turned to measurements of double variants and asked whether double variant effects could be explained by those of the two variants individually. We find that indeed, variant effects are almost perfectly additive in logit space and therefore independent (**Figure 2E**). This result is consistent with an earlier observation that effect sizes associated with short k-mer motifs are additive in log space.^15^

Next, we asked whether our measurements could be predicted with existing splicing models. For this analysis, we excluded variants where the associated reference has exactly 0 or 1 PSI. Variants that disrupt SDs or SAs in these cases have no measurable impact if the reference sequence has a PSI of 0, and therefore, the measured Δlogit is exactly zero. Conversely, models recognize that the corresponding variants indeed weaken the splice signals and can predict a non-zero effect. Likewise, any variants that increase exon inclusion would not be experimentally observable if the reference PSI is 1. Experimentally, such effects might be observable at deeper sequencing depth (i.e., because then rare spliced-in reference transcripts might be detected) or in a reporter context with weaker competing splice signals.

After making these adjustments, we found that splicing predictors HAL, MMSplice, Pangolin, and SpliceAI could accurately predict variant impact (**Figure 2F**).^15–17,19^ SpliceAI in particular performed well on both intronic and exonic single and double variants with a Pearson r of 0.77 (**Figure 2F, left**). For SpliceAI, the predicted PSI was calculated as the product of the donor probability at the expected SD and the acceptor probability at the expected SA on the variable exon, since the model does not directly predict PSI.^53^ We observed performance with Pangolin comparable to SpliceAI, as expected given their similar model architecture.^16^ Older and simpler models, such as MMSplice and HAL, also achieved good accuracy, albeit HAL can only predict exonic variants **(Figure 2F, right)**.^15,17^ The full correlation plots for Pangolin, HAL, and MMSplice are provided (**Figure S2 F-H)**. These results suggest that current splice predictors have learned a fairly accurate representation at least of those aspects of the splicing code that are similar across cell types. To further test the utility of our dataset, we retrained MMSplice on COMPASS, which improved the model’s performance on a held-out test set (r = 0.66 to 0.69). Across the full dataset, the retrained model reached r = 0.77, comparable to SpliceAI (r = 0.78) and markedly higher than the baseline MMSplice (r = 0.66), demonstrating how this resource can be used to fine-tune existing predictive models (**Figure 2G)**.

### COMPASS identifies splice-disrupting variants and suggests potential modes of pathogenicity for variants of uncertain significance

We next turn to the analysis of measured variants with annotations in ClinVar (**Figure 3A**). We observe that variants classified as benign or likely benign (B/LB) generally have a limited impact on splicing, with most Δlogit(PSI) values being close to zero. In contrast, several pathogenic or likely pathogenic (P/LP) variants have a large negative Δlogit(PSI), suggesting that they might act by disrupting splicing; the two highlighted variants in the pathogenic category both disrupt core splice signals. The majority of variants of conflicting or uncertain significance (CL/VUS) have small effect sizes, indicating that they are likely benign or at least do not act by disrupting splicing. However, there are also several CL or VUSs variants with large negative or positive effects. Because of their strong molecular phenotypes, these variants are prime candidates for further investigation as they suggest a plausible mode of pathogenicity through altered splicing. The highlighted, large effect size variants occur in disease-associated genes and are either intronic or disrupt core splicing signals (**Figure 3B**).

**Figure 3.**
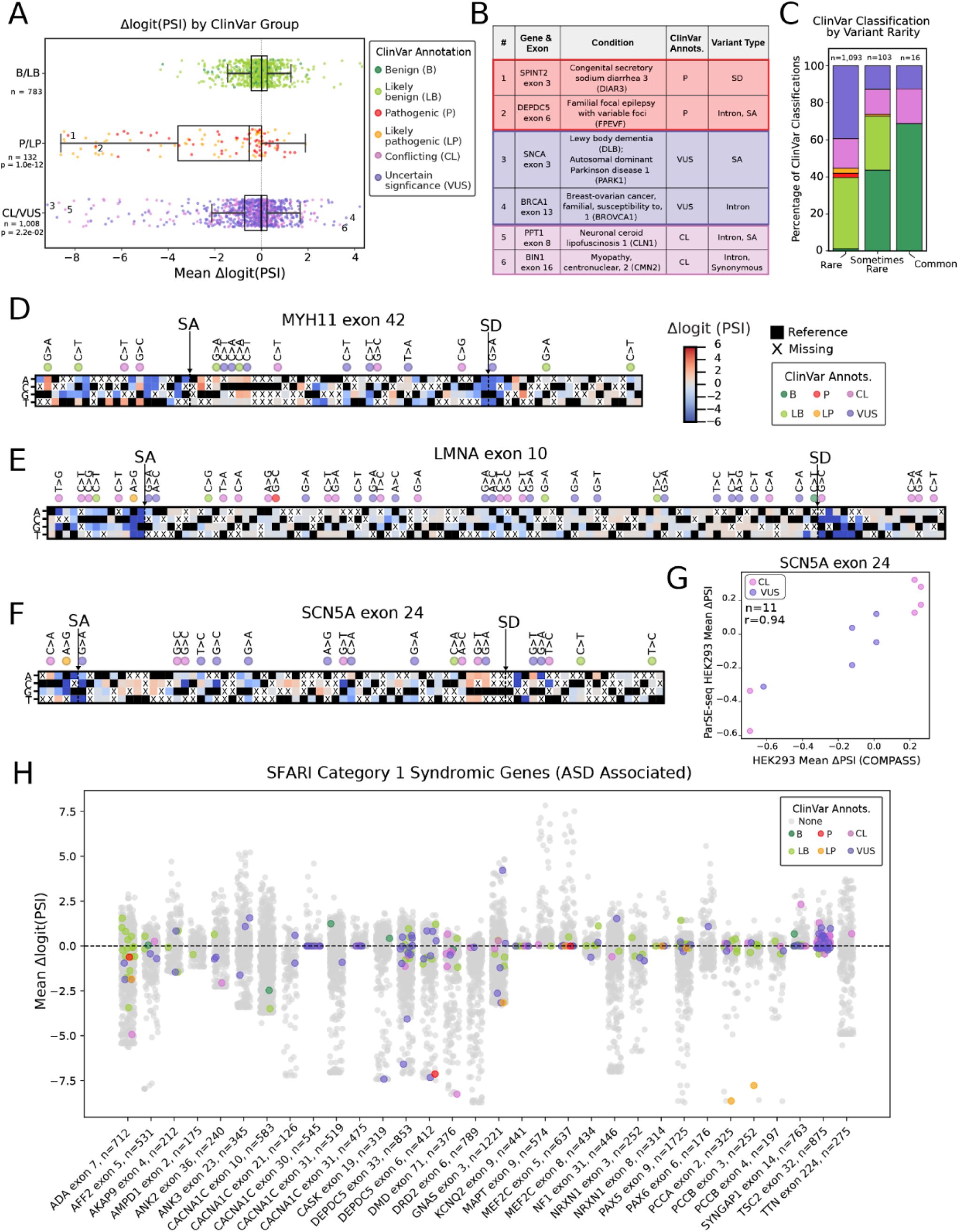
Splicing-associated effects of ClinVar-annotated population variants. (A) Distribution of average Δlogit(PSI) for variants present in all lines. Variants were grouped by ClinVar classification: benign/likely benign (B/LB, n = 783), pathogenic/likely pathogenic (P/LP, n = 132), and conflicting/uncertain significance (CL/VUS, n = 1,008). B/LB variants exhibit consistently lower Δlogit(PSI) magnitudes, whereas P/LP and CL/VUS variants span broader ranges, with many high-magnitude outliers. Fisher’s exact test confirmed significant differences between B/LB and P/LP (*p-*value = 1.0 × 10^−12^) and between B/LB and CL/VUS (*p-*value = 2.2 × 10^−2^), consistent with splicing disruption contributing to pathogenic and uncertain classifications. These results confirm that P/LP variants are frequently associated with aberrant splicing and suggest that a subset of CL/VUS variants may exert pathogenic effects through similar mechanisms. (B) Six representative variants from 3A are highlighted: two pathogenic, two VUS, and two CL. Each is associated with a splicing disruption, including SD/SA disruption or intronic variants annotated in ClinVar. (C) ClinVar classifications stratified by allele frequency categories from ClinVar dbSNP155: rare (n = 1,093), sometimes rare (n = 103), and common (n = 16). Rare variants in dbSNP155 are defined by having a MAF of less than 1%. Common variants are those with a MAF of at least 1%. Sometimes rare variants have a MAF of less than 1% in some, but not all, reporting projects. (D, E, F) Near-site saturation mutagenesis (NSSM) plots span the assayed sequence, with the three possible substitutions at each position colored by their measured Δlogit(PSI), alongside the reference allele (black). Positions marked with an “X” indicate variants absent from the dataset. Shown are sections of NSSMs for (D) MYH11 exon 42, (E) SCN5A exon 24, and (F) LMNA exon 10, all of which are ACMG-listed genes. ClinVar annotations displayed above the heatmaps for variants of known clinical significance. All maps span 161 nucleotides, with truncation applied in this figure; the full length NSSMs are shown in **Figure S3.** (G) PSI values from our assay were compared with ParSE-seq, which measures splicing of SCN5A exons in HEK293 cells using a rat insulin minigene reporter. For ClinVar CL and VUS variants in SCN5A exon 24 (n = 11), the two assays were highly concordant (Pearson r = 0.94). (H) Variants from SFARI category 1 syndromic genes (1S). Of the 240 1S genes in the SFARI Gene database, 32 exons from 24 of these genes were represented in our dataset. For these exons, we show the distribution of cell line averaged variant effects on splicing, with ClinVar-annotated SNVs (n = 309) highlighted. This overlap emphasizes the contribution of SDVs to pathogenic mechanisms in autism-associated genes.

Finally, **Figure 3C** shows a subset of measured variants that have annotations in ClinVar and that also occur in dbSNP and for which we thus have minor allele frequency (MAF) information. As expected, most P and LP variants are rare while most common variants are benign. Very few rare variants are labeled as benign, though more are classified as LB than strictly B. CL and VUS variants were enriched among rare alleles, whereas B and LB classifications made up the majority of common variants, consistent with pathogenic and uncertain annotations being concentrated in rare variants. COMPASS variants derived from ExAC further connect these trends, with the rarest variants showing the strongest splicing effects (**Figure S2K**), linking the observation that pathogenic variants are both rare in the population (**Figure 3C**) and more likely to disrupt splicing (**Figure 3A**).

### Near-site saturation mutagenesis maps of disease-relevant exons

For exons and flanking introns from disease-relevant genes, we generated high-throughput nearsite saturation mutagenesis maps (NSSMs). For this purpose, we define an exon family as a reference sequence (an exon with its flanking introns) together with all associated variants. Each NSSM is represented by at least 150 SNVs in at least one cell line. In total, 107 exon families met this threshold, with variant coverage for each sequence spanning nearly one-third of all possible substitutions and likely perturbing most or all *cis*-regulatory elements in the exon family. Position-based heatmaps were used to visualize the effects of SNVs on exon inclusion across all measured cell lines. Heatmaps display the average Δlogit(PSI) values across measured cell lines for each SNV, with axes indicating the SNV position and nucleotide change. We highlight exons from the ACMG list of clinically actionable genes, including exon 42 of *MYH11* (familial thoracic aortic aneurysm 4), exon 10 of *LMNA* (dilated cardiomyopathy 1A), and exon 24 of *SCN5A* (cardiac arrhythmia syndromes) (**Figure 3D–F**). All three exons are naturally alternatively spliced and are thus particularly susceptible to SDVs.

The *MYH11* gene encodes smooth muscle myosin heavy chain (SM-MHC) and gives rise to two principal isoforms, SM1 and SM2, through alternative splicing at exon 42^54,55^ Exon 42 is included in the SM2 protein isoform but not in SM1, resulting in a shorter, alternative protein C terminus as the exon encodes for only 9 amino acids followed by a stop codon in the *MYH11* reading frame (albeit not in that of the reporter)^56^. A number of pathogenic splice variants have been reported across multiple exons in this gene^54,55,57,58^. In COMPASS, we observe multiple variants with conflicting and uncertain significance in *MYH11* exon 42 with variable effects on splicing. Mutations with conflicting annotations that disrupt a polyC stretch just downstream of the SA coincide with a putative binding motif for the poly(C)-binding proteins PCBP1 and PCBP2, raising the possibility that their strong splicing effects may reflect disruption of RBP binding^59^ (**Figure 3D**). However, these variants are interspersed with benign mutations of similar effect size, suggesting that they are likely benign themselves in spite of their potential to modulate exon inclusion, possibly because isoforms SM1 and SM2 have similar functions. ^59^

Nuclear envelope proteins lamin A and C are both derived from the *LMNA* gene through alternative splicing. A third common splice isoform, lamin AΔ10 results from skipping of exon 10 but is otherwise identical to lamin A and has no defined function^60^ *LMNA* exon 10 contains a large number of VUSs, but our measurements reveal that most have relatively small effect sizes, suggesting that they are unlikely to result in exon skipping **(Figure 3E**). Still, it cannot be excluded that some of the (non-synonymous) variants are pathogenic but directly modify protein stability or folding rather than splicing, similar to the G>C mutation of the *LMNA* exon 10 NSSM heatmap. Moreover, we observe a LP variant overlapping the SA and a CL variant near the SD that both have a strong negative impact, as do multiple other SDVs in the adjacent intronic region.

*SCN5A* encodes the cardiac sodium channel NaV1.5, and its loss-of-function is the leading monogenic cause of the arrhythmia disorder Brugada syndrome (BrS) as well as other cardiac arrhythmias.^61–63^ Prior studies in the *SCN5A* gene have identified intronic and exonic SDVs, which have been associated with BrS.^61,64–66^ COMPASS revealed a spectrum of SDVs, including both variants overlapping ClinVar annotations, such as a pathogenic variant in the SA, and additional SDVs not previously annotated in ClinVar (**Figure 3F**). These measurements also showed strong concordance with the same measurements from the ParSE-seq MPRA in HEK293 cells (**Figure 3G**).

A larger set of exon families with NSSMs includes ACMG exons such as *MUTYH* exon 6, *TNNT2* exons 4 and 5, *RYR1* exon 94, *TSC2* exon 32, and *MYH11* exon 6, as well as exons with a large number of P/LP pathogenic variants or VUSs (**Figure S3**). NSSMs densely sample the mutational landscape, but by definition, they do not capture every possible variant. We thus examined whether predictors that perform well on our measured data, such as SpliceAI, can interpolate the missing positions to generate complete saturation mutagenesis maps. We find that such interpolation often works well (**Figure S4A**), but, as illustrated in other cases (**Figure S4B**), models sometimes predict spurious effects or miss true splicing effects, underscoring that they are not yet a substitute for direct experimental measurement. In this regard, MPRAs provide a powerful tool for systematically validating and refining computational predictions. We next extended our analysis beyond ACMG exons to disease-associated exon families, including those linked to autism spectrum disorder and cancer.

### Genes associated with autism spectrum disorder and cancer

Finally, we turn to a set of genes strongly associated with Autism Spectrum Disorder (ASD) and are categorized as syndromic category 1 genes in the SFARI gene database.^67^ We find that 32 exon families from 24 genes fall into this category, with a minimum of 126 variants and a maximum of 1,221 variants per exon family. Across the gene panel, we observe a large number of SDVs that have not yet been annotated in ClinVar but that are promising targets for further investigation because of their strong splicing phenotypes (**Figure 3H**). Of those variants with existing ClinVar annotations, a pathogenic variant in *DEPDC5* exon 6 has a strong negative impact on splicing. Notably, a VUS in the same gene has a similar impact, suggesting a possible reclassification. Likely pathogenic SDVs are found in *PCCA* exon 2 and *PCCB* exon 3. Conversely, pathogenic variants in ADA exon 7 and *MEF2C* exon 5 have minimal impact on splicing and likely act through different mechanisms. Notably, *MEF2C* exon 5 has a reference PSI near zero in COMPASS (**Supplementary Table 1)**, meaning that only positive effects that increase exon inclusion can be practically detected. Similar circumstances occur in other exons, and we document these cases in **Supplementary Table 1**, which lists near-saturation exon families alongside their starting PSI and the range of observed Δlogit values. The large scale of COMPASS enables similar visualizations for many additional gene categories beyond the SFARI ASD set. Genes from the ACMG SF v3.2 list, COSMIC Tier 1 cancer genes, transcription factors, and cytokines/cytokine receptors are shown in **Data S1**.^68–71^ Additionally, we identify exon families that are highly saturated with more than 300 single and double mutants, as well as “hotspot” exons that are enriched for and particularly susceptible to SDVs (**Data S1**).^72^

### Validation of ClinVar BIN1 exon 16 variants by prime editing

To validate COMPASS-derived splicing effects, we used prime editing (PE) in HEK293 cells to create variants in their native genomic contexts (**Figure 4A**). *BIN1* exon 16 (referred to as exon 17 in some transcripts) undergoes alternative splicing and has been shown to play a role in the onset of cancer, Alzheimer’s disease, and cardiac pathology.^73–76^ To further validate functional splicing consequences at this locus, we selected two ClinVar variants in *BIN1* exon 16 family with conflicting pathogenicity annotations, based on their strong but opposite splicing effects in the MPRA (**Figure 4B**). The chr2:127,051,220:C>G variant in *BIN1* has conflicting ClinVar classifications of LB and VUS, the latter reflecting insufficient evidence to either confirm or rule out pathogenicity. This variant (variant 1) disrupts a poly-G tract and increases exon inclusion in our MPRA.The chr2:127,051,153:C>T variant in *BIN1*, affecting the donor splice site of intron 16, has conflicting ClinVar classifications ranging from VUS to Likely Pathogenic, with the latter submission noting splice site disruption. This variant (variant 2) reduces exon inclusion in COMPASS. PE introduced each variant at the endogenous locus, and genomic sequencing confirmed high editing efficiency for variant 1 (>90%) and lower efficiency for variant 2 (∼28%) (**Figure 4C–D**).

**Figure 4.**
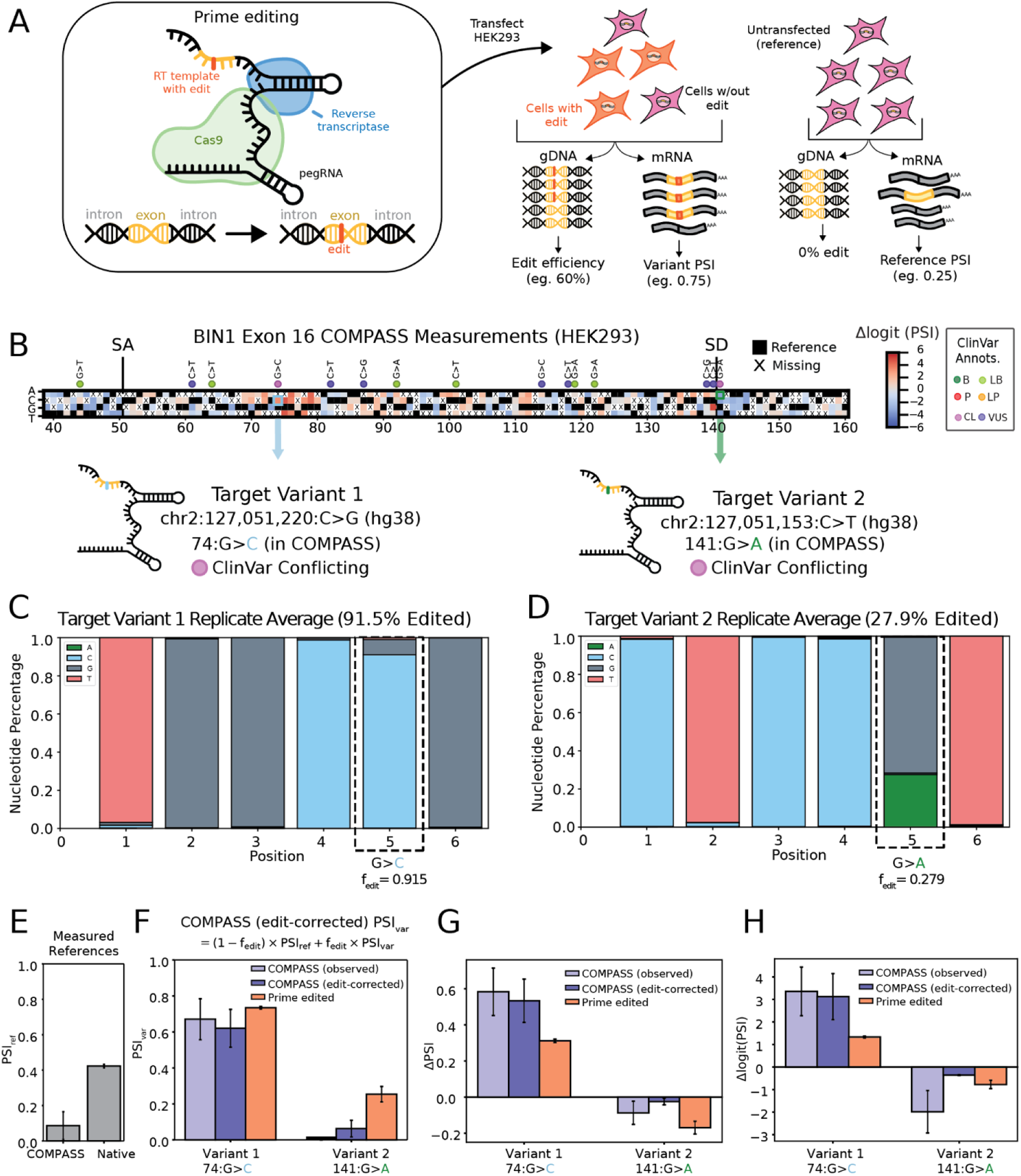
Prime editing validation of *BIN1* exon 16 ClinVar variants. **(A)** Schematic of the prime editing (PE) system and experimental workflow. PE enables introduction of precise base edits in an exon or intron of interest. HEK293 cells were transfected with pegRNAs targeting *BIN1* exon 16, where the reverse transcriptase template specifies the desired substitution (e.g., G>C). Editing efficiency was measured from genomic DNA (gDNA) sequencing, and splicing outcomes were quantified from mRNA sequencing. An unedited population of HEK293 cells served as the control reference. **(B)** NSSM results for *BIN1* exon 16 in HEK293 cells, highlighting the two ClinVar variants chosen for validation. target variant 1 is chr2:127,051,220:C>G (hg38), corresponding to position 74:G>C in the COMPASS construct. Target variant 2 is chr2:127,051,153:C>T (hg38), corresponding to position 141:G>A in the MPRA construct. **(C)** Editing efficiencies for variant 1 measured from gDNA sequencing in two biological replicates of PE-edited HEK293 cells (fraction edited in repl. 1 = 0.919, fraction edited in repl. 2 = 0.910 ; Average fraction edited = 0.915). **(D)** Editing efficiencies for variant 2 measured from gDNA sequencing (fraction edited in repl. 1 = 0.281 and fraction edited in repl. 2 = 0.277; Average fraction edited = 0.279). **(E)** Reference PSI values (unedited controls) measured in MPRA context and in endogenous context both in HEK293. **(F)** Variant PSI values measured in COMPASS, the edit-corrected expectation derived from COMPASS, and the observed PE PSI in HEK293 cells. Edit-corrected values were calculated from COMPASS reference and variant PSI values, weighted by the average editing fraction f_edit_ across both PE replicates for each variant. **(G)** ΔPSI values in MPRA and PE contexts for both variants, comparing variant PSI to the respective reference PSI. **(H)** Δlogit(PSI) values calculated as demonstrating concordance of variant effects across MPRA and PE contexts after accounting for baseline PSI differences.

We measured the reference PSI in unedited HEK293 cells and compared it to the corresponding COMPASS reference (**Figure 4E**). Variant PSIs were then measured in PE-edited cells and compared to MPRA values, with an edit efficiency-corrected MPRA PSI calculated to account for the mixture of edited and unedited alleles (**Figure 4F**). ΔPSI values in both contexts showed concordant effects (**Figure 4G**), and Δlogit(PSI) further strengthened this agreement by correcting for baseline differences in reference PSI (**Figure 4H**). Variant 1 consistently increased exon inclusion in both MPRA and PE assays, while variant 2 consistently reduced inclusion. Even in the native locus, variant 2 did not fully ablate exon inclusion despite changing the canonical U1 SD recognition sequence, suggesting U12 may partially rescue this SS.^77^ Overall, these experiments demonstrate that MPRA-measured effects closely match those in the endogenous context. Both *BIN1* variants, despite conflicting ClinVar annotations, showed reproducible splicing changes in the same direction across MPRA and PE.

An obvious question raised by this variant analysis is whether we can identify mechanisms that might explain impacts on splicing. One hypothesis is that SDVs have a large effect because they disrupt a cis-regulatory element important for splicing. For example, the strong effect of disrupting the polyG in *BIN1* exon 16 is consistent with these sequences acting as splicing silencers and could indicate the potential involvement of RNA-binding proteins that recognize this element. However, outside of SDs and SAs, recognizing functional RBP binding sites and their cognate RBPs can be challenging.

### Effect sizes can be associated with RBP binding motifs

To identify putative RBP binding sites, we assembled a database of RBP-associated motifs from ATtRACT, CISBP-RNA, and oRNAment, which integrates data from *in vitro* binding assays (SELEX, RNAcompete) with select curated *in vivo* CLIP-derived motifs.^78–80^ The combined database contains 1,007 motifs represented as position weight matrices (PWM). Using FIMO, we scanned all library sequences with the PWM collection to identify statistically significant matches corresponding to putative RBP binding sites (**Figure 5A**).^81^

**Figure 5.**
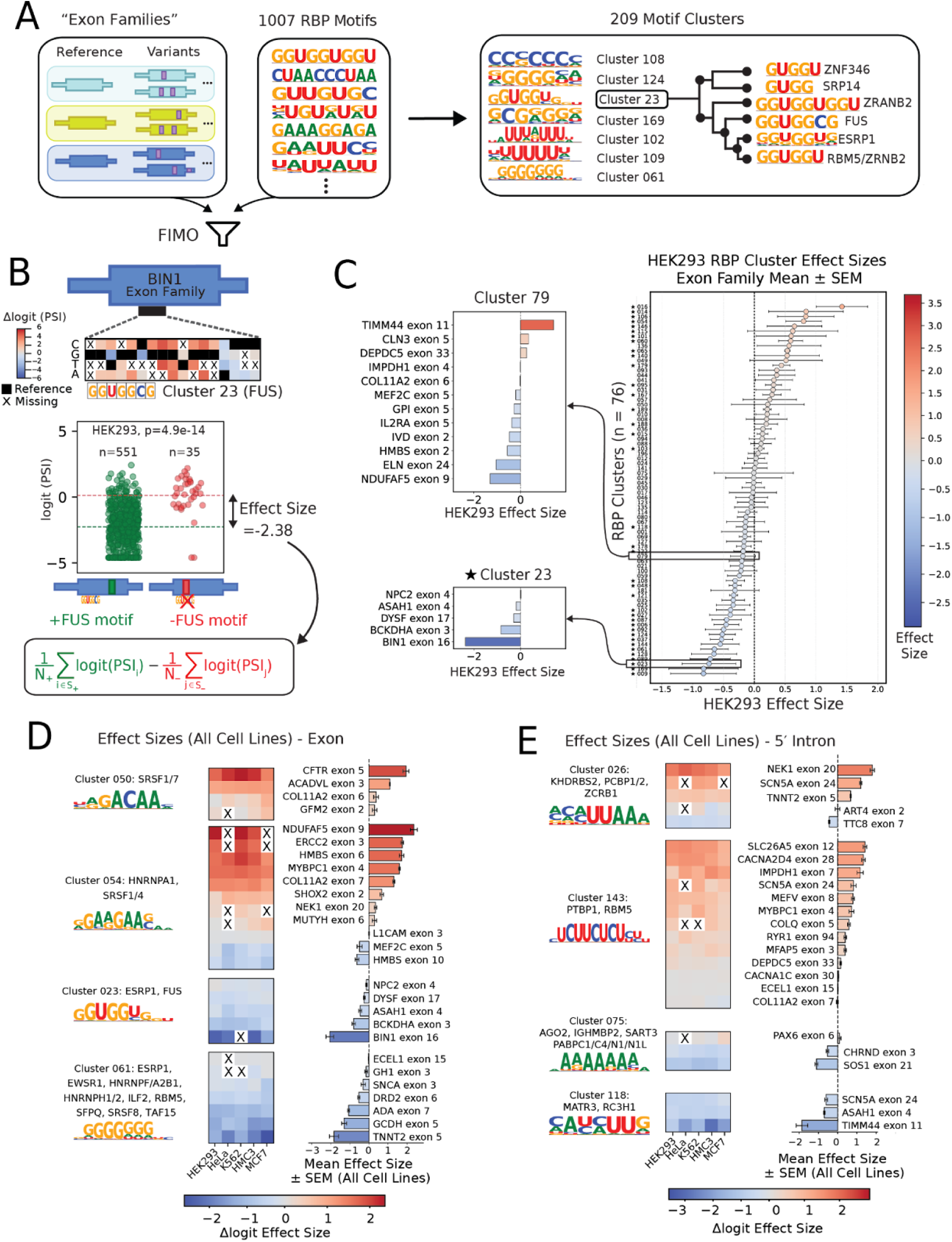
Cis-regulatory effects of RBP motifs across exon and intron regions. (A) Workflow of data processing prior to motif effect size analysis. Exon families were defined as groups of sequences containing a reference sequence and its corresponding variants. We compiled a set of 1,007 RNA-binding protein (RBP) motifs from the ATtRACT, CISBP-RNA (Ray et al. 2013), and oRNAment databases. These motifs were collapsed into 209 representative clusters to reduce redundancy, and the clusters were used for downstream effect size analyses. All sequences were scanned for motif matches using FIMO, and the resulting clusters were used for downstream effect size analyses. (B) Example of motif effect size analysis. Effect sizes were calculated within exon families and defined regions. For each motif identified in a sequence by FIMO, we compared the set of sequences with a significant hit (q ≤ 0.01) to those without a detectable hit. In the example of *BIN1* exon 16 measured in HEK293, variants (single or double) were grouped into motif-present and motif-absent sets. The equation defines the effect size as the difference in mean logit(PSI) between the motif-present and motif-absent groups, yielding a Δlogit(PSI) motif effect size. This analysis was performed within exon families and within specific regions (exonic and intronic) and repeated across all cell types. Further details are provided in the Methods. (C) Concordance filtering of motif effect sizes in HEK293. Motif clusters with effect size calculations in at least three exon families are shown. Each point represents the mean Δlogit(PSI) across exon families in HEK293, with error bars denoting the SEM. Clusters marked with a star pass the concordance filter (≥75% agreement in effect direction across exon families). Cluster 23 exemplifies consistent effects across contexts (including *BIN1* as in panel B), whereas cluster 79 does not pass concordance, reflecting differential effects of this motif across exon families. (D) Concordant RBP motifs in exonic regions. Examples of motif clusters with consistent effects across exon families are shown. SR protein motifs act as exonic splicing enhancers (ESEs) that promote exon inclusion, whereas G-rich motifs act as exonic splicing silencers (ESSs), in line with their previously reported repressive activity in MPRAs.^15^ (E) Concordant RBP motifs in intronic regions. Shown are examples of motif clusters with consistent effects across multiple exon families. PTBP1/RBM5 motifs function as intronic splicing enhancers (ISEs) that promote exon inclusion, consistent with PTBP1 binding to extended CU-rich elements with and has been hypothesized to regulate splicing in a location dependent manner^121–124^ Poly(A)-associated motifs, recognized by PABPs, are shown as intronic splicing silencers (ISSs) which are often known to regulate splicing on terminal introns^125–127^ Other motifs are shown as ISSs or ISEs for comparison.

Next, we asked whether we could infer the importance of an RBP binding site from the impact of variants that disrupt it. To reduce redundancy of similar motifs across and within databases, and to streamline subsequent analysis, we clustered individual motifs into 209 consensus motif clusters, a common approach used for transcription factor motifs that has also been successfully applied to RBP motifs.^82–84^ For each pair of motif cluster and exon family, we then divide the family into members that contain a motif from that cluster and those that do not. A sequence matching a motif in the reference exon may be sufficiently altered by an overlapping variant such that it no longer constitutes a significant match to the motif PWM in FIMO. Conversely, a variant might introduce a motif match even if the reference does not match the motif. We then calculate an average logit(PSI) for the two groups within each cell line and define the difference of these averages as the motif effect size (**Figure 5B**). The example shows exon 16 of *BIN1* which contains a good match for the FUS (fused in sarcoma) binding motif from oRNAment. This motif overlaps variant 1 which we characterized using PE above (**Figure 4B**). The family members that do contain the motif on average, have a more negative logit than those that do not contain it, resulting in an overall negative effect size for the motif. This negative effect size is consistent with the cognate RBP, likely FUS, being an exonic splice silencer (ESS), consistent with literature.^85^ The result is also consistent with the native variant impact measured in the PE experiment.

Splice site variants were excluded from this analysis because their effects are typically large, direct, and mediated through canonical splicing machinery, rather than through auxiliary RBPs. Double variants, however, were retained to increase the number of sequence perturbations considered, with the caveat that PSI values for a given motif family member may deviate from the reference due to a second variant distal to the motif of interest; this bias is minimized because both motif-containing and motif-lacking groups include such variants.

We generalized motif effect size calculations to all motif–exon family pairs in each cell line, provided that within that cell line, there were at least ten family members containing the motif and at least ten lacking it. **Figure 5C** shows data for HEK293 as a representative example, with all cell line data provided in **Figure S5A,B**. We calculated effect sizes separately for motifs in the leading intron, central exon, and trailing intron, since RBPs are known to exert different regulatory effects depending on where they bind.^86–88^

Motifs belonging to Cluster 23, which includes motifs associated with ESRP1 in addition to FUS, have putative binding partners in five exon families (**Figure 5A)**. All observed instances are associated with a negative effect size, with the most negative effect size observed for *BIN1*, exon 16 (**Figure 5B, C**). Conversely, Motif Cluster 79, which includes motifs associated with ERI1, PCBP2, PCBP1, and HNRNPK, has a mixture of negative and positive effect sizes when comparing different exon families (**Figure 5C**).

There may be several reasons for the effect size variation observed for the same motif cluster across different exon families (**Figure 5C**). First, each motif cluster aggregates multiple similar motifs, each of which may be recognized by a different RBP (**Figure 5A**), and therefore different RBPs may be responsible for the observed splicing effects in different exon contexts. Still, we chose to calculate effect sizes at the cluster level because accurately assigning a particular instance of the motif to a particular RBP is challenging. Second, even for identical motifs recognized by the same RBP, local sequence context, including distance to splice sites, RNA secondary structure, and the presence of neighboring RBP motifs, can influence the regulatory outcome.^89^ Finally, technical factors such as uneven sequencing depth across exon families may introduce additional noise, which can make RBP-associated effects present differently across families.

For further analysis, we focus on motifs for which the effect sizes have predominantly the same sign across all exon families (**Figure S5C**). Examples of motifs with concordant effect sizes are shown in **Figure 5D,E**. **Figure 5D** shows examples of exonic splice regulators. These include a motif associated with SRSF1/7 that is found in four exon families and has a positive effect size across all cell lines, consistent with the known role of SR proteins as exonic splice enhancers.^83–86^ Conversely, polyG motif clusters associated with multiple RBPs, including several HNRNP family members, act as ESSs as previously described.^85,94–99^ **Figure 5E** shows intronic splice enhancers and silencers. A polypyrimidine motif found to recruit PTBP1 and RBM5 broadly acts as an intronic splicing enhancer (ISE), consistent with prior reports.^93,94^ Additional examples of motifs further show that variation of effect sizes between cell types tends to be more subtle than those between different exon contexts (**Figure 5D, E, Figure S5D, E**).

### Cell type-specific exon inclusion

As noted above, splicing patterns are highly correlated between cell types (**Figure 1D**) and, consequently, RBP effect sizes are also broadly similar (**Figure S5D,E**). Still, there are many individual sequences that exhibit cell type-specific splicing. To identify sequences that are most differentially spliced in one cell type relative to all others, we calculated a mingap score defined as the absolute difference between the PSI in the target cell type and the next closest PSI in any of the other cell types^45^ and then ordered mingap scores by size (**Figure 6A**). For four of the five cell lines (HeLa, K562, MCF7, HMC3), we were able to identify at least one sequence with strong cell-type specificity, here defined by a mingap ≥0.25 between the most and second most included cell line. For **Figure 6A**, we retained only the top 20 sequences per cell type based on their mingap ranking, but in total 21 sequences for HeLa, 8 for K562, 88 for MCF7, and 99 for HMC3 met the criterion for strong specificity (**Figure S6B).**

**Figure 6.**
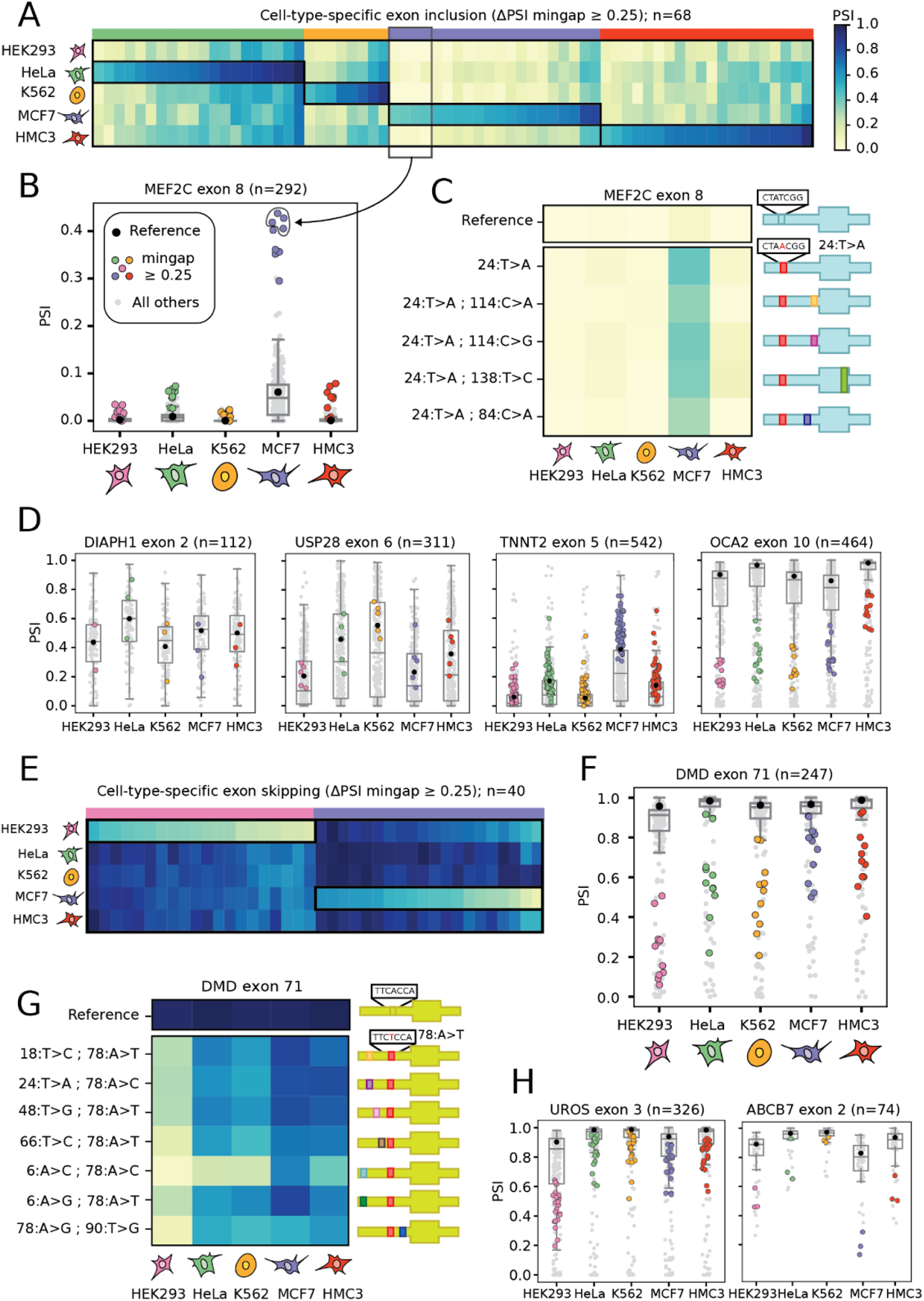
Cell type–specific splicing regulation revealed through ΔPSI mingap analysis. **(A)** Heatmap of sequences with cell-type-specific exon inclusion, identified using the ΔPSI mingap ≥ 0.25 threshold, defined as the absolute difference between the highest PSI value and its next closest value. Up to the top 20 events per cell line were retained (HeLa, 20; MCF7, 20; HMC3, 20; K562, 8). **(B)** Swarm plot of MEF2C exon 8 sequences across cell lines. 9 variants pass the mingap ≥ 0.25 threshold in MCF7 and are colored for MCF7, with the reference in black. The same 9 variants are colored in the other four cell lines for comparison. .**(C)** Example of a recurrent event: a single 24T>A variant in MEF2C exon 8 repeatedly observed to drive MCF7-specific inclusion. Statistical testing confirms reproducible differences across sequences. **(D)** Representative exon families where many variants drive cell-type-specific inclusion, including DIAPH1 exon 2 (HeLa; n = 3), USP28 exon 6 (K562; n = 5), TNNT2 exon 5 (MCF7; n = 52), and OCA2 exon 10 (HMC3; n = 13). Variants passing the ΔPSI mingap ≥ 0.25 threshold in the indicated cell line are shown in color, with the reference PSI in black. These variants are also colored in the other four cell lines for comparison. The total number of sequences in each exon family is shown. **(E)** Heatmap of sequences with cell-type-specific exon skipping, identified using the ΔPSI mingap ≥ 0.25 threshold, defined as the absolute difference between the lowest PSI value and its next closest value. Up to the top 20 events per cell line were retained (HEK293, 20; MCF7, 20). **(F)** Swarm plot of DMD exon 71 sequences across cell lines, a representative example of cell-type-specific skipping (n = 11 variants with ΔPSI mingap ≥ 0.25). Variants meeting the threshold are shown in color, with the reference PSI in black. The same 11 variants are colored in the other four cell lines for comparison.**(G)** Recurrent variants in DMD exon 71 passing the ΔPSI mingap ≥ 0.25 threshold exhibit HEK293-specific exon skipping and MCF7-specific inclusion. **(H)** Representative exon families where many variants drive cell-type-specific skipping, including UROS exon 3 (HEK293; n = 23) and ABCB7 exon 2 (MCF7; n = 3). Variants passing the ΔPSI mingap ≥ 0.25 threshold in the indicated cell line are shown in color, with the reference PSI in black. The same variants are colored in the other four cell lines for comparison.

To evaluate the robustness of observed cell type-specific splicing effects, we next asked whether the same shift recurs across multiple independent sequences within the same exon family. Variants associated with exon 8 of myocyte enhancer factor 2C (*MEF2C)* provide an intriguing example. This exon is referred to as exon β in the literature and is included in transcripts in the brain and striated muscle but excluded elsewhere. MEF2C encodes a transcription factor, and exon β inclusion has been shown to enhance its transcriptional activation capacity.^102–104^ In our reporter, most members of the *MEF2C* exon 8 family are excluded across all cell lines. However, 9/292 family members are differentially included (mingap ≥0.25) in MCF7 cells.

Intriguingly, we identified a recurrent mutation at the 24th position (T>A) that promotes exon inclusion in MCF7 cells (**Figure 6E, F**). This effect was observed across five independent sequences, including one single-nucleotide change and four double mutants that also carried additional substitutions elsewhere in the sequence. The reproducibility of the 24T>A change across sequences provides a high-confidence example of cell type-specific variant activity. The variant position overlaps a motif from Cluster 54 that encompasses several SR proteins and hnRNPs (HNRNPA1, SRSF1, SRSF4, SRSF5, SRSF6, and TRA2B), implicating altered RBP binding as a potential driver of the observed specificity. *MEF2C* is a known oncogene and is expressed in epithelial tumor cells of breast cancers^105,106^; however, its precise role in this context remains unclear.

Additional representative examples of cell type-specific exon inclusion are *OCA2* exon 10 (high in HMC3), *USP28* exon 6 (high in K562), *DIAPH1* exon 2 (high in HeLa), and *TNNT2* exon 5 (high in MCF7) (**Figure 6B**). Many of these exons are alternatively and cell type-specifically spliced.^107–109^ In *DIAPH1* exon 2, two of the three cell type–specific variants occur at base 30 and overlap cluster 44 (HNRNPK, PCBP1). The RBPs listed in the parentheses are the best matches from all motifs in the cluster for this particular sequence. In *TNNT2* exon 5, 52 variants exceed the mingap threshold, including 5 at position 12 that overlap cluster 026 (ZCRB1, PCBP1/2) and cluster 138 (KHDRBS3), both of which function as ISEs (**Data S3**). For *USP28* exon 6 exons, no clusters provided sufficient evidence to explain the observed specificity, suggesting contributions from additional or uncharacterized regulatory motifs.

### Cell type-specific exon skipping

Applying the same approach to exon skipping **(Figure 6E**), we identified strong evidence of preferential exon exclusion, defined by mingap ≥ 0.25, in two of the five cell lines (HEK293 and MCF7). In total, 614 sequences in HEK293 and 37 in MCF7 were selectively skipped in the target cell line while being broadly included in the others (**Figure 6E, S6A)**. These results hint at global cell line-specific biases in exon inclusion. A relevant example is *DMD* exon 71, where recurrent variants at position 78 reproducibly yielded HEK293-specific skipping while also slightly increasing MCF7-specific inclusion (**Figure 6G**). This variant site overlaps clusters 092 (PCBP2/4) and 063 (HNRNPK), and effect sizes for Cluster 063 match the MCF7 inclusion phenotype, consistent with disruption of an intronic splicing enhancer. Notably, DMD exon 71 is very short and shares characteristics with neuronal microexons, which are highly conserved short exons strongly regulated by RBPs.^110–113^ Additional representative examples of cell type-specific skipping are *UROS* exon 3 (low in HEK293) and *ABCB7* exon 2 (low in MCF7) (**Figure 6H**). In *UROS* exon 3, 23 variants surpass the mingap threshold; 5 map to position 108 and overlap cluster 144 (TIA1L1), which functions as a strong ESS with slightly stronger effects in HEK293 as observed in the effect size analysis (**Data S2**). Additional variants overlap cluster 201 (SRSF1) and cluster 048 (ELAVL2). We were not able to calculate effect sizes for these clusters because they were not observed sufficiently frequently. For *ABCB7* exon 2, which is known to be alternatively spliced,^114^ no clusters provided sufficient evidence to explain the observed specificity, suggesting contributions from additional or uncharacterized regulatory motifs. In both cell type-specific inclusion and skipping, most of the exon families that support highly cell type-specific variants also exhibit cell type-selectivity across the entire family. Global exon family differences are accentuated by specific variants, yielding pronounced cell-type-specific splicing outcomes.

### Other forms of cell type-specific splicing

So far, we have focused on the two most dramatic types of cell type-specific splicing, where an exon is included (or excluded) in one cell type versus all others in our panel. However, there is a more general class of splice events where differential splicing occurs between two cell lines with intermediate PSI values for the other three. A representative example of that group is *COLQ* exon 5, where a large fraction of family members exhibit markedly higher exon inclusion in HeLa compared to HEK293 (**Figure S6C**). For further characterization of this differential splicing, we selected one member of the *COLQ* exon 5 family (**Figure S6D**). We first confirmed that HeLa PSI of the original variant from the MPRA is approximately twice that of HEK293 in the context of a gel assay (**Figure S6F**). *COLQ* exon 5 contains motif hits for motif clusters 61 (HNRNPH1/2/3/F), 80 (SRSF3), and 166 (SRSF9,) and we next asked whether disrupting any of these motifs would change the observed cell type bias. We created three additional splicing reporters with two or three mutations each, disrupting one motif at a time (**Figure 6E**). Motif disruption experiments revealed distinct contributions of motifs from clusters 80 (SRSF3), 166 (SRSF9), and 61 (HNRNPH1/2/3/F2). The putative SRSF3 binding motif acted as a strong enhancer: disruption of the motif reduced PSI to zero in both cell lines as measured on a gel. Disruption of the SRSF9-binding motif had a very slight positive effect in HEK293 but a fairly strong negative impact in HeLa, where it reduced PSI by about half. Finally, HNRNPH1/2/3/F functioned as a HeLa-specific silencer as deletion of the corresponding motif resulted in increased PSI in HeLa but not HEK293 (**Figure S6F).** This example demonstrates how local motif architecture and differential RBP activity can impact cell type-specific splicing regulation. Expression data from the Human Protein Atlas (proteinatlas.org) (**Figure S6G**) showed that HNRNPH2 is more highly expressed in HeLa, consistent with it acting as a stronger silencer in this cell line, whereas the effects of SRSF9 were not fully consistent with their expression patterns.^115^

## DISCUSSION

Deciphering how sequence variation and tissue or cell type context shape pre-mRNA splicing remains a central challenge in understanding RNA regulation and disease. Here, we report COMPASS, the largest splicing variant MPRA to date, screening over 87,000 single and double variants across 2,096 exons to map their effects on splicing in both disease-relevant and cell type–specific contexts. Much of our data is focused on disease-relevant genes, such as ACMG genes or genes containing pathogenic ClinVar variants. To reclassify variants of unknown or uncertain significance, we propose a simple heuristic: variants in exons already harboring known P/LP variants should be considered higher priority if they cause similar or more severe splicing disruption, whereas variants with effects similar to B/LB variants should be deprioritized. However, even in the absence of known pathogenic reference variants, the Δlogit(PSI) score can serve as a prioritization metric by comparing a variant’s impact to other variants within the same exon, particularly when splicing alterations in other exons of the same gene are implicated in disease. We highlight examples from disease relevant genes such as *MYH11* (familial thoracic aortic aneurysm 4), of *LMNA* (dilated cardiomyopathy 1A), and SCN5A (cardiac arrhythmia syndromes) as well as broader gene categories such as 32 exons from SFARI Category 1 Syndromic Genes associated with ASD, 13 exons from ACMG genes, 14 exons from COSMIC Tier 1 genes, and 12 “hotspot” exons, among others.

Care has to be taken in how to interpret the results of any splicing MPRA. Physiological variant effects are proportional to ΔPSI and the resulting changes in downstream protein isoform levels. However, ΔPSI is highly sensitive to the starting PSI, which may be different in the reporter assay and the endogenous context. Consequently, the properties of the reference exon in the cellular context of interest are a useful filter for identifying candidate pathogenic SDVs. For example, we highlighted several naturally alternatively spliced exons (**Figure 3**) because such exons are typically more sensitive to SDVs. Recent work also introduced the concept of “hotspot” exons, arguing that exons flanked by relatively weak core splice signals, containing many RBP binding motifs, and for which there is evidence for existing splice variants, are more susceptible to SDVs.^72^ We also reiterate that there are situations where a splicing MPRA cannot accurately capture variant impact. For example, if a reference exon is fully spliced out in the MPRA (i.e. zero reads for the spliced in isoform), variants that further disrupt splicing signal will show no measurable change.

Still, we emphasize that in general, MPRA results are good predictors of the variant effects observed in the genomic context, especially when differences in starting PSI are taken into account. We use PE to directly validate this correspondence for two variants with conflicting ClinVar annotations in exon 16 of *BIN1*, a gene implicated in cancer, Alzheimer’s disease, and cardiac pathology.^73,75,76^ Both variants are found to be splice-disrupting with large effect sizes in COMPASS and the genome editing experiments.

Measured Δlogit(PSI) scores are generally well predicted by splicing models and our experiments provide further support for the application of these models for identifying SDVs. One near-term application is the use of splice predictors to fill in gaps in experimental data (e.g. NSSMs) at least in those cases where the available data for the exon in question is well predicted. However, we also expect that such models can generalize to situations that will remain outside the reach of experiments for the foreseeable future such as making saturation mutagenesis predictions at genome scale.^116^ We also expect that the dataset provided here can be used to further finetune existing splice predictors as we demonstrated in the context of retraining MMSplice. Finally, predictions of Δlogit(PSI) can highlight potential variant effects irrespective of experimental context.

As expected, many SDVs modify core splice sites but we also identify many that instead modulate other aspects of the splicing code such as modifying putative RBP motifs or activating cryptic splice sites (**Figure S2E**). While many RBPs and their corresponding *cis*-regulatory binding motifs have been identified and compiled in databases such as ATtRACT, CISBP-RNA, oRNAment^78–80^, we have yet to fully characterize the quantitative relationships between *cis*-regulatory motif variation, cellular context and splicing outcomes. Here, we took a step toward addressing this challenge by using motif perturbations to quantify the impact of RBP binding motifs on exon inclusion. Earlier work, such as MaPSy, established important precedent by comparing reference to SNVs in specific contexts.^24^ However, that approach was limited to pairwise comparisons. By assaying thousands of SNVs across many motif occurrences, COMPASS enables a large-scale and robust quantification of motif effects on splicing regulation. Our double-variant data suggest that variant effects are largely additive in log space, supporting a relatively simple model where (non-overlapping) motifs act largely independently to modulate splicing (**Figure 2E**). We note that further work is needed to establish causal links between motifs and the RBPs that recognize them. Mapping from RBP to motif is usually not unique and many splice regulatory proteins are associated with multiple, often divergent motifs determined using different measurement methods or biological contexts. Conversely, a single putative binding site is often compatible with motifs for multiple different RBPs. A further confounder is that RBP levels may differ from those of the corresponding RNAs because of differences in stability or post-translational modifications required for protein activity.

We systematically compared splicing of tens of thousands of reporter sequences across five cell lines of diverse tissue origins. We found that splicing patterns are highly correlated across different cell types but also identified subsets of sequences with cell type-specific splicing patterns. Strongly differentially spliced exons in COMPASS (mingap of ≥0.25) are rare with fewer than 1% of sequences identified as cell type-specific. At the exon family level, fewer than 6% of families contained even a single sequence classified as cell type-specific by this criterion. These findings agree with earlier splicing MPRAs comparing smaller numbers of cell types and sequences.^23–25^ A recent MPRA measuring mRNA stability across six different cell lines similarly found that differential regulation is the exception rather than the rule.^117^ This relative homogeneity of splicing patterns across cell lines may be explained by the fact that the features underlying splicing decisions are largely shared across cell types, reflecting the widespread activity of RBPs that are broadly expressed and functional across cellular contexts.^118^ Moreover, the use of plasmid-derived splicing reporters may result in some saturation effects that could dampen the observed cell type-specificity and it is also possible that certain splicing factors are not sufficiently present in the cell lines used. Thus, additional validation of these exons in endogenous settings is suggested and we have shown two examples with COMPASS concordance. Nonetheless, these results are consistent with findings from primary neurons which identified hundreds of differentially spliced exons for a given cell type of interest when assaying exon inclusion at transcriptome scale.^119,120^ Models that treat splicing in a cell type-agnostic manner capture broad patterns of variant impact, but they may not fully account for events that depend on cell-type context. Incorporating cell type-specific regulatory information could aid in the interpretation of the subset of variants with cell type-specific splicing outcomes, particularly in disease contexts where pathogenicity derives from certain cell types.

## Funding

This work was supported by NIH awards R01GM149631 and R01CA297834 to GS and by NSF GRFP under Grant No. DGE-2140004 to S.K. AMY was a Washington Research Foundation Postdoctoral Fellow.

## Competing interests

GS, CMR and ABR are a co-founders and shareholders of Parse Biosciences, a single cell RNA sequencing company.

## Code availability

All code is available under an MIT License on GitHub at https://github.com/skoplik/ESL_MPRA_2025

## Data availability

Raw and processed data generated in this study are available in the Gene Expression Omnibus (GEO) under accession number GSE307247. Raw data include DNA-seq and RNA-seq reads from the MPRA library across all replicates. Processed data include pre-calculated splicing metrics such as PSI, ΔPSI, and Δlogit values, together with the complete list of MPRA library sequences and their associated metadata, including genomic coordinates and variant annotations. All supplemental tables are provided with this manuscript, including the full list of primers used in this study.

## Materials availability

The backbone reporter for COMPASS is available at Addgene (#247173).

## METHODS

### Experimental Overview

The exon skipping plasmid MPRA was amplified for DNA-seq in order to construct a map of SNVs and their associated unique barcode sequences. Following transfection of this MPRA in HEK293 cells, RNA-seq was performed to quantify the PSI associated with each unique barcoded reporter.

### DNA-seq Library Preparation

Because each variant contains the same backbone region, there is low nucleotide diversity in each sequence. To minimize the amount of PhiX needed for Illumina sequencing, “offset” nucleotides between the Illumina sequencing adapter region and a plasmid specific primer were used in order to create diversity of bases in generated amplicons for increased percent of clusters passed filtering. To prepare the plasmid library for DNA-sequencing, 1ng of plasmid was initially amplified for 5 cycles with primers that bind the plasmid, including those “offset” sequences to generate library diversity. Following this PCR, qPCR with P5/P7 Illumina primers was used to selectively amplify the products containing sequencing adaptors and to prevent over-amplification. Amplification was conducted and monitored with qPCR and stopped early to minimize PCR biases. Samples were size selected to keep fragments ≥ 100nt using KAPA Pure Beads (Roche). Sample size distributions were analyzed using Tapestation High Sensitivity D1000 (Agilent).

### DNA-seq

Library concentrations were quantified using qPCR quantification with the NEBNext Library Quant Kit for Illumina (NEB) as well as Qubit 1x dsDNA HS (Thermo Fisher). Concentrations between the two methods were averaged to determine optimal library loading concentrations for sequencing. Sequencing of DNA libraries, with 2% spiked-in PhiX, was iteratively performed on 300 cycle MiSeq Reagent V2 kits (Illumina) until the desired sequencing coverage and depth was obtained. Custom primers were used with 251 cycles on read 1, 45 cycles on read 2, and 12 cycles on Index 1. Reads from multiple DNA-seq interations were concatenated and processed together to build a comprehensive map of DNA barcodes.

### Cell Culture and Transfection

HEK-293 cells (ATCC, CRL-1573), MCF7 cells (ATCC, HTB22), and HeLa cells (ATCC, CCL-2.2) were cultured in DMEM (Gibco). K-562 cells (ATCC, CCL-243) were cultured in RPMI (Gibco). HCM3 cells (ATCC, CRL-3304) were cultured in EMEM (ATCC). All media was supplemented with 10% fetal bovine serum (Cytiva) and 1% penicillin/streptomycin (Gibco) and cells were cultured at 37°C and 5% CO2. For transfection, cells were seeded, to a density of 200,000 cells/mL in a 10 cm plate, and transfected with 15 μg of plasmid library using Lipofectamine 3000 (Thermo Fisher) with at least 2 biological replicates for each library. Cells were harvested for RNA extraction 36–48 h after transfection. Less than 5% of the cells were used for flow cytometry to confirm transfection efficiency and the remaining cells were used for RNA extraction.

### RNA-seq Library Preparation

RNA extraction was performed with the Monarch® Total RNA Miniprep Kit (NEB). mRNA was isolated using the NEBNext Poly(A) mRNA Magnetic Isolation Module (NEB) with 5μg of total RNA input per sample. The resulting mRNAs were reverse transcribed with SuperScript IV Reverse Transcriptase (Thermo Fisher) to generate cDNA using an RT primer specific to the 3′ end of the transcripts (downstream of the barcode sequence). Library cDNA was then amplified with KAPA HiFi HotStart ReadyMix (Roche) for 5 cycles with a forward primer that binds upstream of the Citrine exon 1 splice junction and a reverse primer that binds downstream of the barcode region. As with the DNA library preparation, additional “offset” sequences to generate library diversity were added to the 5′ end of the amplicon. A second 5 cycle PCR step was done using a forward primer that binds PE1 and P5 and a reverse primer that binds P7. Lastly, qPCR with P5/P7 Illumina primers were used to selectively amplify the products containing sequencing adaptors and to prevent over-amplification. Amplification was conducted and monitored with qPCR and stopped early to minimize PCR biases. Samples were size selected to keep fragments ≥ 100nt using KAPA Pure Beads (Roche). Sample size distributions were analyzed using Tapestation High Sensitivity D1000 (Agilent).

### RNA-seq

Library concentrations were quantified using qPCR quantification with the NEBNext Library Quant Kit for Illumina (NEB) as well as Qubit 1x dsDNA HS (Thermo Fisher). Concentrations between the two methods were averaged to determine optimal library loading concentrations for sequencing. Libraries were sequenced on a NextSeq 500/550 Mid Output (300 cycle) kit using custom primers, with 212 cycles on read 1 to span the library sequence into both splice junctions, 86 cycles on read 2 for barcode sequences, and 12 cycles on Index 1 for UMI and indices. Libraries were pooled with 10% PhiX prior to sequencing.

### Bioinformatic Workflow

DNA-seq reads were used to extract 20-nt barcodes from R2, anchored by a required 20-bp forward shared region (allowing up to 5 mismatches). Barcodes were clustered with starcode (distance = 1), and consensus barcode plus insert sequences supported by ≥5 reads were assembled into a FASTA reference for STAR indexing. Reference FASTA entries were assembled from each unique 20-nt barcode cluster. For each barcode cluster and library member combination, the reference sequence consisted of the last 25 nt of Citrine exon 1, SMN2 intron 6, the variable exon and intronic flanks, SMN2 intron 7, Citrine exon 2, and the 20-nt barcode. The matched barcode plus insert references were used to build STAR genome indexes for RNA-seq mapping. Resulting RNA-seq reads are passed through a bioinformatic pipeline to count exon skipping or inclusion events. Here, we take advantage of STAR’s capability to detect and count expected and novel exon-exon spanning reads to determine splice isoform abundance. Accurately counting PSI for single reporter constructs will be pivotal for the future development of machine learning models to predict isoform abundance based on sequence. Thus, a mini MPRA library spanning 13 sequences sampled from the full DNA library was used to optimize a computational pipeline for producing accurate PSI calculations (**Figure S2D**). The optimized pipeline 1) clusters DNA barcodes using starcode, 2) collapses unique molecular identifiers (UMIs) and initial STAR alignments to remove duplicate RNA reads caused by PCR amplification, and 3) uses STAR for mapping and subsequent PSI calculation (17-19). Once this pipeline was optimized, the mini MPRA library resulted in PSI values that correlated well with PSIs for the same sequences found in the full MPRA (**Figure S2D**). Following bioinformatic processing, PSIs were determined from the pooled included and excluded reads across two biological replicates, requiring a minimum of 10 reads per replicate spanning the expected splice junctions and at least two replicates per cell line. We note that replicate-averaged PSIs are highly correlated with replicate pooled PSIs (**Figure S1B**). For global analyses, we averaged the replicate pooled Δlogit(PSI) values for each variant across cell lines.

### Validation of MPRA plasmid backbone performance

To confirm that the COMPASS plasmid backbone did not introduce artificial splicing effects independent of the inserted variable sequence, we performed control experiments with reference exons that have well-characterized exon skipping behavior. Four exons were selected based on their association with human disease and extensive prior study: SMN2 exon 7 (54 nt), MAPT exon 10 (93 nt), CFTR exon 12 (87 nt), and DMD exon 29 (150 nt)^128–131^ Each exon was cloned with intronic flanks into the same plasmid backbone used for construction of the COMPASS library.

Splicing outcomes were assayed by transfecting plasmids into cells and measuring exon inclusion by RT-PCR with primers in Citrine flanking the splice junctions. Isoform ratios were quantified from gel band intensities in Image Lab, which correlated with published values (Pearson r = 0.83). PCR products were also sequenced by RNA-seq to provide a higher-resolution benchmark, which yielded similar results (Pearson r = 0.85 with literature values). PSI values determined by gel and RNA-seq were highly correlated with each other (Pearson r = 1.0) (**Figure S2A**). These results validate that the plasmid backbone itself does not bias exon skipping and is suitable for high-throughput MPRA assays.

We further benchmarked COMPASS against previously published alternative splicing MPRAs. Beyond the ParSE-seq comparison (**Figure 3G**), we evaluated overlap with the MFASS dataset, another large-scale minigene assay.^23^ PSI values were compared for more than 350 overlapping variants between COMPASS and the MFASS dataset, revealing a strong correlation (Pearson r = 0.74) (**Figure S2C**). Importantly, these results were obtained despite the use of different plasmid backbones in the two designs (SMN1 intronic sequence for MFASS versus SMN2 intronic sequence for COMPASS), demonstrating that splicing MPRAs are generalizable and robust across sequence contexts.

### Logit and ΔLogit PSI Transformations

PSI values of 0 and 1 are undefined on the logit scale. Therefore, PSI values were clipped to the interval [0.01,0.99] prior to logit transformation. This clipping threshold is consistent with that employed by FRASER, a splicing-focused method that similarly addresses numerical instability on the logit scale at the PSI extreme values.^49^ We selected 0.01 and 0.99 as empirically justified clipping thresholds to match the resolution imposed by the average read depth of our library, with mean per-sequence coverage across pooled replicates ranging from 112 (MCF7) to 242 (HEK293). This approach also ensures compatibility on the logit scale between variants and references, which differ in average read depth. To maximize sequencing depth per measurement, we pooled replicates to compute a single Δlogit(PSI) value per variant sequence. We confirmed that this pooling strategy yields results highly consistent with averaging Δlogit(PSI) values calculated from individual replicates, with Pearson r values ranging from 0.96 to 1.0 across comparisons (**Figure S1B**).

### Model predictions with SpliceAI

SpliceAI predictions were generated using the original ensemble of five trained models.^19^ For prediction, we used the 161nt sequence in our reporter construct (Citrine exon 1, SMN2 intron 6, variable exon, SMN2 intron 7, Citrine exon 2), resulting in a 1,563 nt sequence. To satisfy SpliceAI input requirements, sequences were further padded with 10,000 Ns (5,000 upstream and 5,000 downstream). Predictions were run on both single and double variants.

SpliceAI outputs donor and acceptor probabilities at each base in the input sequence. For our sequences, the relevant positions are the annotated SD and SA of the variable exon. To approximate exon inclusion, we defined the predicted PSI as the product of the donor probability at the expected SD and the acceptor probability at the expected SA, reflecting the requirement that both sites be recognized for splicing to occur^53^ These PSI values were then clipped to the interval [0.01, 0.99] and transformed to logit space. Δlogit PSI was then calculated as the difference between variant and reference logits for direct comparison with experimental measurements and predictions from other models. Sequences with reference PSI measurements of exactly 0 or 1 were excluded when assessing model performance, as such cases are unlikely to show measurable variant effects but can still be scored by models.

### Model predictions with Pangolin

Pangolin predictions were obtained using the probability output model, which predicts the probability that a site is used as a splice site in a given tissue. Sequences were prepared identically to those used for SpliceAI: each 161 nt variable sequence in the reporter context (Citrine exon 1, SMN2 intron 6, the variable exon, SMN2 intron 7, Citrine exon 2) and padded with 5,000 nucleotides of Ns on both sides. Predictions were run with each of the four tissue-specific models (heart, liver, brain, and testis). For every sequence, probabilities were recorded at the expected SA and SD) for each tissue. Probabilities were clipped to the range [0.01, 0.99] and transformed to logit space. For each tissue, we calculated Δlogit(PSI of SD) and Δlogit(PSI of SA) as the difference between alternate and reference logits at the SD and SA, respectively. To obtain a single PSI value and summarize across tissues, we used the absolute maximum of Δlogit(PSI of SA) and the absolute maximum of Δlogit(PSI of SD) values observed. We then averaged the Δlogit(PSI of SA) and Δlogit(PSI of SD) maxima to use as a representative Δlogit(PSI) value. PSI values were defined following the Pangolin methods as the maximum difference in splice site probabilities across tissues, using the strongest SD and SA values (which could come from different tissues) and averaging these maxima to yield a single PSI estimate per sequence. In our approach, this maximum-difference criterion was applied in logit space rather than probability space.^16^ As with SpliceAI, sequences with reference PSI measurements of exactly 0 or 1 were excluded when assessing model performance.

### Model predictions with MMSplice

Variant effects were predicted using MMSplice with the VCFDataloader.^17^ To enable this, we generated custom FASTA, GTF, and VCF files for our synthetic sequences. For predictions, we used the 161 nt sequence in our reporter construct (Citrine exon 1, SMN2 intron 6, variable exon, SMN2 intron 7, Citrine exon 2), resulting in a 1,563 nt sequence. The FASTA was composed of these full-length sequences (1,563 nt) for the references only. The GTF file defined the reference transcript length as 1,563 nt and annotated all exon boundaries, including those of the variable exon and both Citrine exons in the reporter. The VCF contained all variant positions and base substitutions. For double mutants, the VCF sequence was defined over the interval spanning both substitutions, such that the reference and alternate alleles encompassed the entire region. Predictions were limited to variants within 100 nt of the variable exon due to the default overhangs implemented in the MMSplice model; thus, intronic variants located further than 100 nt from the exon were not scored. As done with all other model predictions, sequences with reference PSI values of exactly 0 or 1 were excluded when assessing model performance.

### Model predictions with HAL

HAL predictions were obtained using the HAL web server (http://splicing.cs.washington.edu/SE).^15^ Reference PSI values were averaged across replicates within each cell line, then provided to HAL on the percent scale after clipping to 0.01–99.99 as required by the tool. HAL predicts ΔPSI for substitutions within the exon or the first 6 nucleotides of the downstream intron; single- or double-mutant sequences with variants outside this window were not used for predictions. For comparison across models and with experimental data, predicted reference and variant PSIs were converted from percent to fraction, clipped to [0.01, 0.99], transformed to logit space, and used to compute Δlogit(PSI). As done with all other model predictions, sequences with reference PSI values of exactly 0 or 1 were excluded when assessing model performance.

### MMSplice retraining and evaluation

In order to retrain MMSplice, PSI values from the five cell lines were clipped between 0.01and 0.99. The logit values derived from variant PSIs were averaged across the five cell lines. Delta logit PSI values were obtained by subtracting the reference logit(PSI) value of each family from the logit(PSI) value of each variant in that family.

Coverage for each family was computed by taking the average read count from each cell line and summing across all samples of that family. The average coverage (for purposes of sorting) was computed by dividing the total coverage of the exon family by the number of sequences in that family, which includes the reference and variants.

To maximize coverage in the test and validation splits, while keeping the train coverage moderately high, families were first sorted by average coverage, then each third family was separated. This produced three pools of families where each pool contained at least the third highest average coverage family. To obtain the final 5% test, 5% validation and 90% training data splits, a knapsack algorithm was used to maximize the sum of family coverage for the split with the size constraint of 5% of the total number of samples. This was necessary since families contained a variable number of samples. To keep PSI values balanced, these pools were further split into two sub-pools where the first sub-pool contained the top 50% PSI values, and the second sub-pool contained the bottom 50%. Knapsack was run on each sub-pool to obtain a 2.5% split each, which were then joined together to obtain the final 5% split. The algorithm was repeated for the validation pool, which included samples not chosen during the processing of the test pool. All the remaining samples were put into the training split.

MMSplice model retraining was performed by maintaining the existing MMSplice architecture and executing hyperparameter tuning using the hyperband tuner from the keras-tuner package. L1 and L2 regularization parameters were tuned in the range [0.01, 0.001] (step size = 0.001) for each convolutional layer. The dropout rate was tuned in the range [0.1, 0.4] (step size = 0.05) for each dropout layer, and the learning rate was tuned in the range [1.0 × 10 ^−7^, 1.0 × 10 ^−5^] (step size = 9. 9 × 10^−7^). The model was trained for 50 epochs after finding the hyperparameter combination that minimized the validation loss. Early stopping returned a model with the best validation loss during this training step. The model was trained with the Adam optimizer using TensorFlow 2.14.

Evaluation of the retrained MMSplice model was compared to predictions from the previously published MMSplice and SpliceAI models on the same held-out test set (n = 3,526) as well as the entire dataset for which variants occurred within MMSplice’s input window (n = 70,544).

### Motif Effect Size Analysis

We assembled a library of RNA-binding protein (RBP) motifs by aggregating position weight matrices (PWMs) from the ATtRACT, oRNAment, CISBP-RNA, and Ray et al. 2013 databases^4–6^, retaining only those annotated for *Homo sapiens*. Redundant PWMs across databases were merged, and the resulting set was clustered with the RSAT matrix-clustering tool^75^, which we modified to disable reverse-complement comparisons. Clustering was performed using normalized correlation (Ncor) as the similarity metric, complete linkage, a correlation threshold of 0.50, Ncor threshold of 0.40, width threshold of 3, and reverse-complement comparisons disabled. This procedure produced a non-redundant set of motif clusters representing distinct RBP binding preferences.

All sequences from the COMPASS library (including both reference and variant constructs) were scanned against the unclustered PWM set using FIMO from the MEME suite v5.5.0, with reverse complement scanning disabled. Only significant motif matches were retained based on FIMO q-value^110^. Because the MPRA library includes systematic single- and double-nucleotide mutations of each reference sequence, we grouped each reference and its associated variants into an “exon family.” Within each family, we identified which sequences contained at least one significant hit to a motif in each cluster and which did not.

We computed Δlogit(PSI) values for each exon family in each cell line by comparing the logit-transformed PSI values of the hit-containing vs. non-hit sequences. PSI values were calculated from pooled replicate read counts and clipped to the interval [0.01, 0.99] before logit transformation. To avoid artifacts from splice site disruption, variants overlapping the core SD or SA (defined as ±4 nt from either junction) were excluded from the analysis. Effect size analysis was performed separately for each cell line. RBP motif effect sizes were calculated for exon families that contained at least 10 variants in both groups (≥10 hit and ≥10 non-hit). These calculations were considered further only if this criterion was satisfied in at least three exon families.

For each RBP cluster, we then calculated a concordance score to assess the consistency of effects across families. Concordance was defined as the weighted sign of Δlogit(PSI), computed as the sum of Δlogit(PSI) values divided by the sum of their absolute values. This score ranges from –1 (consistently negative) to +1 (consistently positive), with values near 0 indicating inconsistent directionality. We retained clusters with absolute concordance ≥0.75 in at least three exon families and observed in at least two cell lines (**Figure S5C**). For these clusters, Δlogit(PSI) values were averaged across cell lines to estimate overall regulatory impact (**Figure S5D,E**).

### Mingap cell-type-specific analysis

To quantify cell-type-specific splicing, we defined the ΔPSI mingap metric as the separation between the most extreme PSI value for a sequence and its next closest value across cell lines. Specifically, ΔPSI mingap high was calculated as the absolute difference between the highest PSI and the next highest PSI, reflecting cell type-specific exon inclusion. Conversely, ΔPSI mingap low was calculated as the absolute difference between the lowest PSI and the next lowest PSI, reflecting cell type-specific exon skipping. The larger of the two was taken as the overall ΔPSI mingap value for a sequence with measurements present in all 5 cell lines. Variants with ΔPSI mingap ≥ 0.25 were classified as cell type-specific.

### Prime Editing

Prime editing experiments were performed following the workflow described in Doman et al^132^ PEmax (cas9-RT fusion protein, Addgene ID: 174820), pU6-tevopreq1-GG-acceptor (cloning vector, Addgene ID: 174038), and hMLH1dn (mismatch repair suppressor, Addgene ID: 178114) were ordered from Addgene. Guide epegRNA were designed using Predict and cloned into the pU5 backbone using Golden Gate assembly^133^. After being sequence verified, individual colonies of all plasmids were grown in LB with carbenicillin, and plasmids were extracted using Qiagen Midi prep kits. HEK293 cells were seeded in 24 well-plates with a density of 100,000 cells/well in HyClone DMEM with High Glucose at 36 C and 5% CO2. Cells were transfected with 1.94 ug of DNA in ratio 3:1 Prime editing vector: epegRNA vector using Lipofectamine 3000 (Thermo Fisher) with at least two replicates of each condition. Cells were harvested for genome extraction 48 hours after transfection or reseeded for subsequent transfections. Up to 5 subsequent transfections were performed for maximum editing efficiency. Cells were detached using TrypLE (Gibco) and resuspended in PBS with Proteinase K Between 1 and 5% of cells were used for flow cytometry to confirm transfection. gDNA and total RNA were extracted from the remaining cells using the NEB Monarch Total RNA Kit and mRNA extraction was performed using the NEBNext Poly(A) mRNA Magnetic Isolation Module. The resulting mRNAs were reverse transcribed with SuperScript IV Reverse Transcriptase (Thermo Fisher) to generate cDNA using an anchored polyT primer. Target PCR using KAPA HiFi HotStart ReadyMix (Roche) is used to amplify the region surrounding the edit (∼300 bp). Genomic PCR amplification was performed with a single set of primers while cDNA amplicons were generated using nested PCRs with a total of 25 cycles for both conditions. Genomic and cDNA fragments were sequenced using Plasmidsaurus Premium PCR (performed with Oxford Nanopore MinION R10.4.1). From the genomic DNA samples, reads were filtered for those containing exact matches for both the expected primer sequence as well as the gRNA sequence. Editing efficiency was calculated as the ratio of reads containing the expected edit at the expected location over the total number of reads correctly assigned to the reference sequence. For RNA read alignment, reference sequences were generated for the exon-included and exon-skipped isoforms. UCSC Genome Browser sequences for BIN1 exon 16 and its surrounding exons were used to define these references. Each read was compared against these two reference sequences with Bio.pairwise2 (up to 3 mismatches allowed) and assigned to the matching isoform. Reads were classified as included or excluded and PSI was calculated of the ratio of included reads over the total number of reads.

## Supporting information

sequences and primers used in experiments

## Notes

https://github.com/skoplik/ESL_MPRA_2025

