## Supplementary Information for "Massively parallel assay of human splice variants reveals cis-regulatory drivers of disease-associated and cell type-specific splicing regulation"

### Supplementary Figures

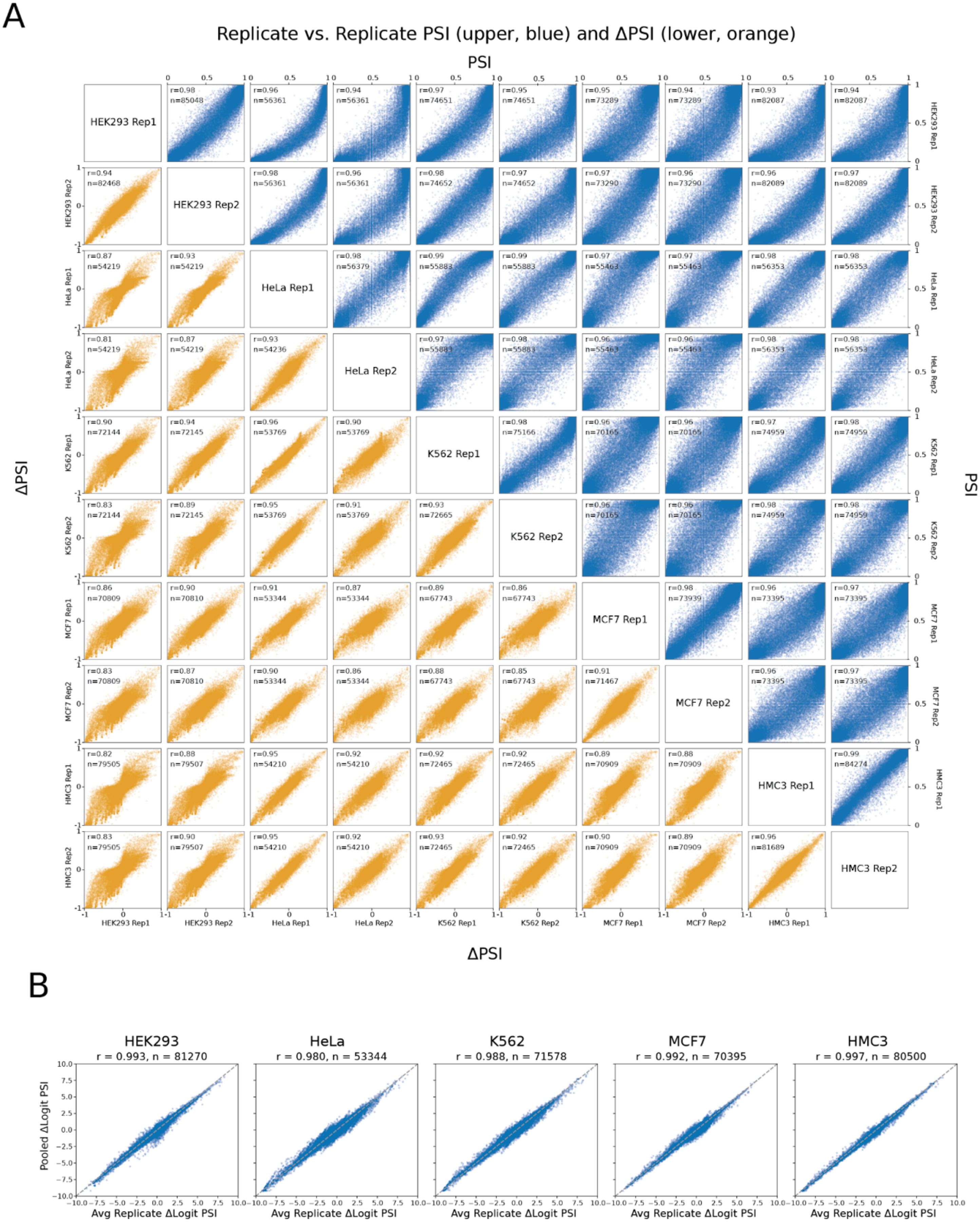
A
Replicate vs. Replicate PSI (upper, blue) and ΔPSI (lower, orange)
PSI
ΔPSI
PSI
ΔPSI
HEK293 Rep1
HEK293 Rep2
HeLa Rep1
HeLa Rep2
K562 Rep1
K562 Rep2
MCF7 Rep1
MCF7 Rep2
HMC3 Rep1
HMC3 Rep2
B
HEK293
r = 0.993, n = 81270
HeLa
r = 0.980, n = 53344
K562
r = 0.988, n = 71578
MCF7
r = 0.992, n = 70395
HMC3
r = 0.997, n = 80500
Pooled ΔLogit PSI
Avg Replicate ΔLogit PSI

**Figure S1 | Correlations of cell line replicates (PSI and ΔPSI) and replicate-averaged versus pooled Δlogit(PSI)**

**(A)** PSIs are shown in the upper triangle in blue, covering two replicates each for HEK293, HeLa, K562, MCF7, and HMC3. ΔPSIs are shown in the lower triangle in orange, covering two replicates each for HEK293, HeLa, K562, MCF7, and HMC3. Each panel displays the number of sequences included and the Pearson r. **(B)** To maximize sequencing depth per measurement, replicates were pooled to compute a single Δlogit(PSI) value per variant sequence. Comparison with Δlogit(PSI) values obtained by averaging across individual replicates confirmed that pooling yields highly consistent results in all five cell lines: HEK293 (r = 0.993, n = 81,270), HeLa (r = 0.980, n = 53,344), K562 (r = 0.988, n = 71,578), MCF7 (r = 0.992, n = 70,395 ), and HMC3 (r = 0.997, n = 80,500).

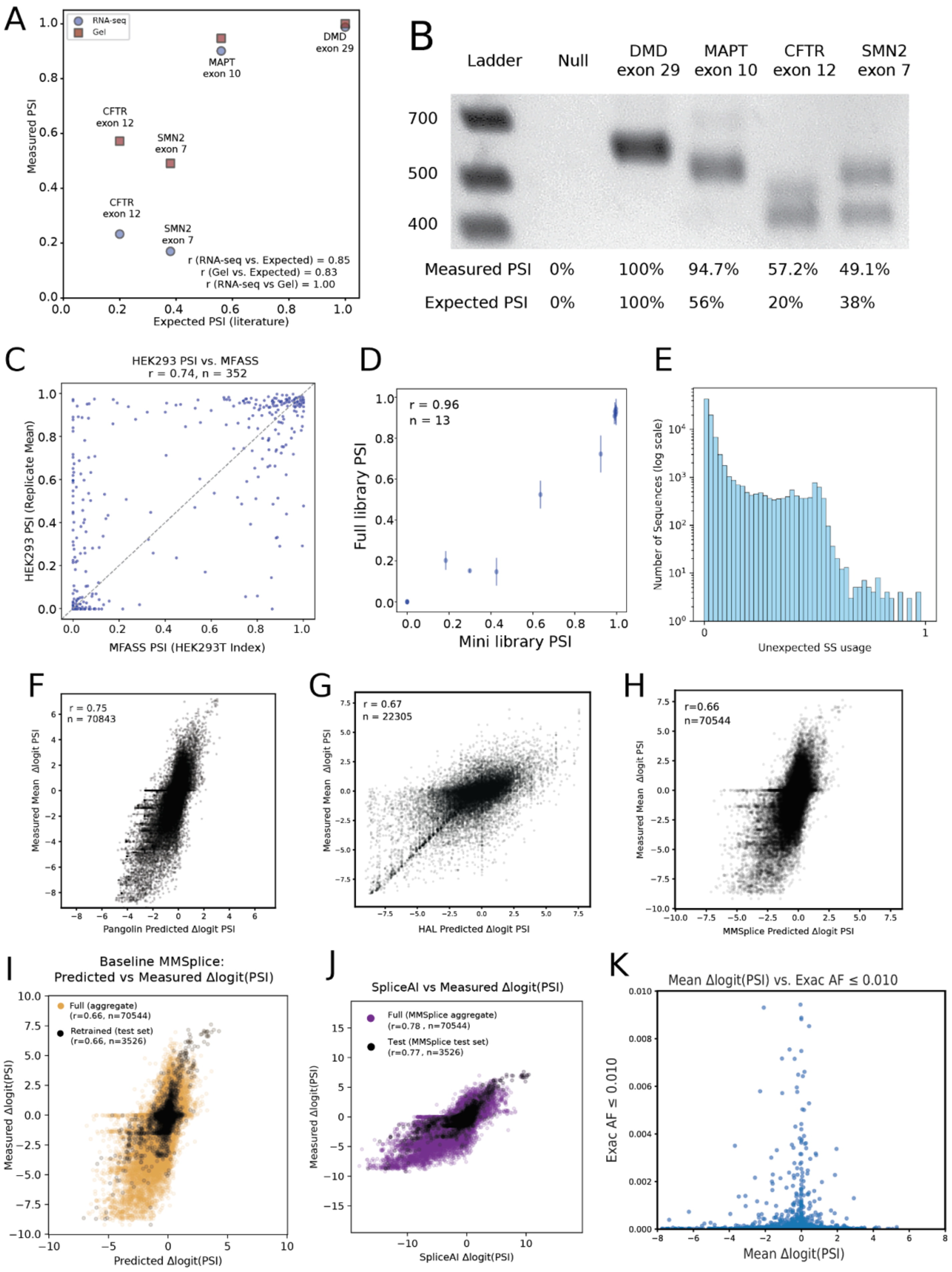
A
RNA-seq
Gel
DMD
exon 29
MAPT
exon 10
CFTR
exon 12
SMN2
exon 7
CFTR
exon 12
SMN2
exon 7
r (RNA-seq vs. Expected) = 0.85
r (Gel vs. Expected) = 0.83
r (RNA-seq vs Gel) = 1.00
Measured PSI
Expected PSI (literature)
B
Ladder
Null
DMD exon 29
MAPT exon 10
CFTR exon 12
SMN2 exon 7
700
500
400
Measured PSI 0% 100% 94.7% 57.2% 49.1%
Expected PSI 0% 100% 56% 20% 38%
C
HEK293 PSI vs. MFASS
r = 0.74, n = 352
HEK293 PSI (Replicate Mean)
MFASS PSI (HEK293T Index)
D
r = 0.96
n = 13
Full library PSI
Mini library PSI
E
Number of Sequences (log scale)
Unexpected SS usage
F
r = 0.75
n = 70843
Measured Mean Δlogit PSI
Pangolin Predicted Δlogit PSI
G
r = 0.67
n = 22305
Measured Mean Δlogit PSI
HAL Predicted Δlogit PSI
H
r=0.66
n=70544
Measured Mean Δlogit PSI
MMSplice Predicted Δlogit PSI
I
Baseline MMSplice:
Predicted vs Measured Δlogit(PSI)
Full (aggregate)
(r=0.66, n=70544)
Retrained (test set)
(r=0.66, n=3526)
Measured Δlogit(PSI)
Predicted Δlogit(PSI)
J
SpliceAI vs Measured Δlogit(PSI)
Full (MMSplice aggregate)
(r=0.78 , n=70544)
Test (MMSplice test set)
(r=0.77 , n=3526)
Measured Δlogit(PSI)
SpliceAI Δlogit(PSI)
K
Mean Δlogit(PSI) vs. Exac AF ≤ 0.010
Exac AF ≤ 0.010
Mean Δlogit(PSI)

**Figure S2 | Validation of assay design, reproducibility of splicing measurements, and predictions from alternative splicing models.**

**(A)** Disease-associated exons with well-characterized splicing outcomes (SMN2 exon 7, MAPT exon 10, CFTR exon 12, DMD exon 29) were tested in the MPRA plasmid backbone. Splicing outcomes were assayed by transfecting plasmids into cells and measuring exon inclusion by RT-PCR with primers in Citrine flanking the splice junctions. Isoform ratios were quantified from gel band intensities in Image Lab, which correlated with published values (r = 0.83) **(B)** Gel image from A with annotated PSIs as estimated from Image Lab. **(C)** PSI values for >350 shared variants were correlated between this MPRA and the independent MFASS assay (r = 0.74), despite differences in plasmid backbones, demonstrating robustness and generalizability of the approach. **(D)** A 13-plasmid mini-library in HEK293 verified by RNA-sequencing, reproduced PSI values observed in the full MPRA (r = 0.96). **(E)** Distribution of unexpected splice site usage across the reporter library. For each sequence, the fraction of reads mapping to noncanonical splice sites was calculated within each cell line. Histograms depict the number of sequences ($\log_{10}$ scale) at a given frequency of unexpected splicing. Shown is the overall average of these frequencies across cell lines. **(F-H)** Scatter plots compare measured Δlogit(PSI) values to model predictions for **(F)** Pangolin (r = 0.75, n = 70,843), **(G)** HAL (r = 0.67, n = 22,305), and **(H)** MMSplice (r = 0.66, n = 70,544).[1–3] **(I)** Scatter plots show predicted versus measured Δlogit(PSI) for the baseline MMSplice model. Each point corresponds to an individual variant, with black indicating the held-out test set from the retraining procedure in main text **Figure 2G** and orange as the full aggregate dataset. Pearson r and sample sizes are shown for each dataset. **(J)** Scatter plots show predictions from the SpliceAI model, plotted for the same sequences and in the same format as in I, using the same aggregate and held-out test sets from the MMSplice retraining in main text **Figure 2G**. This allows direct comparison of SpliceAI predictions to the same sequence sets that MMSplice can score**. (K)** ΔPSI values for variants present in ExAC are plotted against their allele frequency, restricted to rare variants (MAF < 0.01).

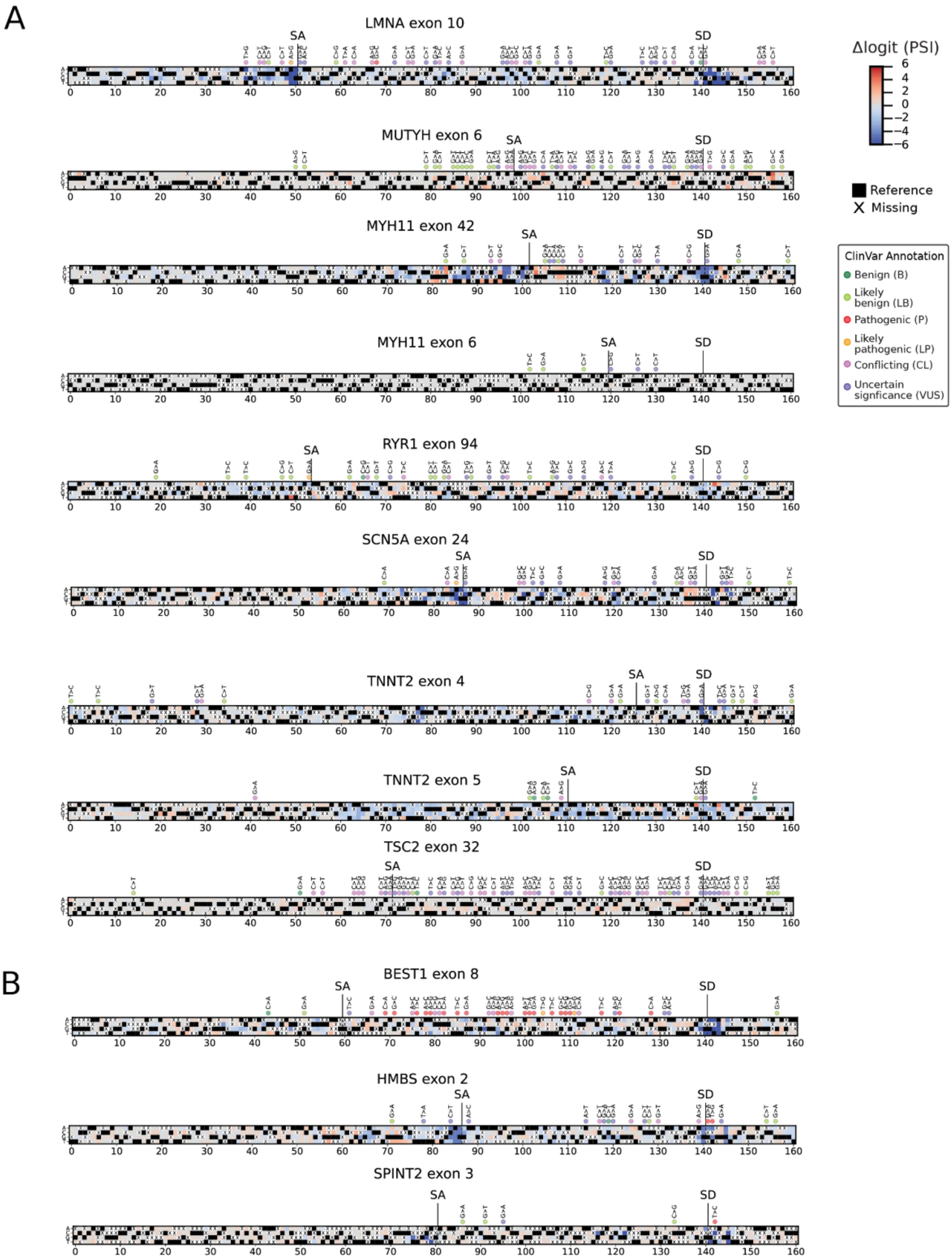
A
LMNA exon 10
MUTYH exon 6
MYH11 exon 42
MYH11 exon 6
RYR1 exon 94
SCN5A exon 24
TNNT2 exon 4
TNNT2 exon 5
TSC2 exon 32
B
BEST1 exon 8
HMBS exon 2
SPINT2 exon 3
SA
SD
Δlogit (PSI)
6
4
2
0
−2
−4
−6
Reference
Missing
ClinVar Annotation
Benign (B)
Likely benign (LB)
Pathogenic (P)
Likely pathogenic (LP)
Conflicting (CL)
Uncertain signficance (VUS)
0 10 20 30 40 50 60 70 80 90 100 110 120 130 140 150 160

**Figure S3 | Near-saturation mutagenesis (NSSM) maps of exons in disease-relevant genes.**
(A) Shown are NSSM plots spanning the full assayed sequence, with each position represented by the three possible nucleotide substitutions colored by their measured Δlogit(PSI) alongside the reference allele in black. Positions marked with an "X" indicate variants absent from the dataset. ClinVar annotations are displayed above the heatmaps for variants of known clinical significance. The full NSSMs are shown here for MYH11 exon 42, SCN5A exon 24, and LMNA exon 10, which are displayed in truncated form in Figure 3D–F. Additional exons from ACMG SF v3.2 listed genes are shown, such as MUTYH exon 6, MYH11 exon 6, RYR1 exon 94, TNNT2 exon 4 and 5, and TSC2 exon 32.
(B) Three exons not on the ACMG list but containing a large number of pathogenic or VUS variants are likewise displayed: BEST1 exon 8, HMBS exon 2, and SPINT2 exon 3.

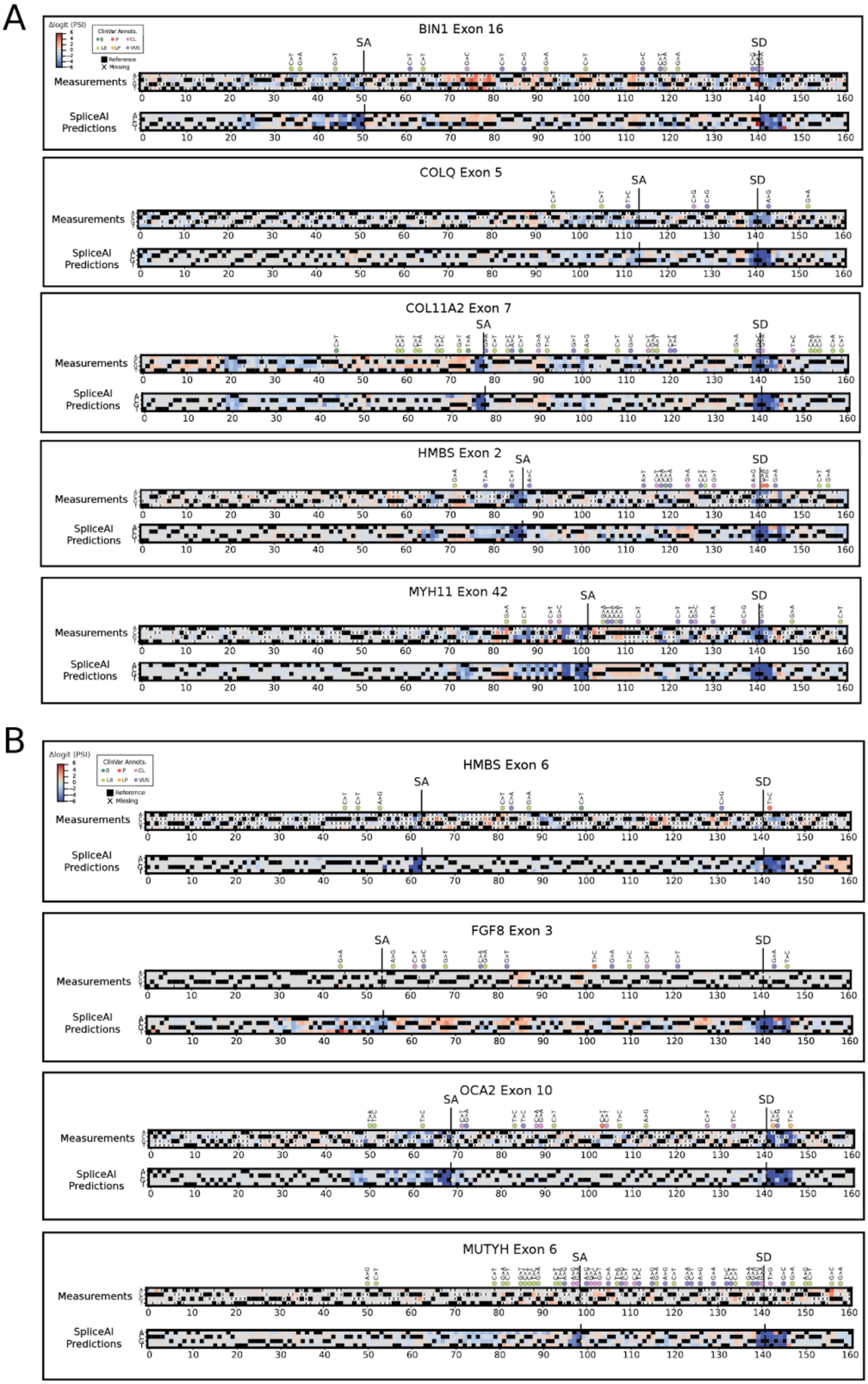
A
Δlogit (PSI)
ClinVar Annots.
Reference
Missing
BIN1 Exon 16
SA
SD
Measurements
SpliceAI Predictions
COLQ Exon 5
COL11A2 Exon 7
HMBS Exon 2
MYH11 Exon 42
B
HMBS Exon 6
FGF8 Exon 3
OCA2 Exon 10
MUTYH Exon 6

**Figure S4 | Predictions of SpliceAI compared with near-saturation mutagenesis maps.**
**(A)** Shown are some NSSM plot examples spanning the full assayed sequence, with each position represented by the three possible nucleotide substitutions colored by their measured Δlogit(PSI), alongside the reference allele in black. Positions marked with an “X” indicate variants absent from the measured MPRA dataset. ClinVar annotations are displayed above the heatmaps for variants of known clinical significance. Below each experimental map, SpliceAI predictions for the same sequence are shown. In these examples, SpliceAI closely recapitulates the position- and allele-specific effects observed experimentally while providing complete coverage across all possible substitutions. **(B)** NSSM examples are plotted the same as in A. However, in these cases, the model predicts spurious splice donor or acceptor effects that are not always supported by the data, or it fails to capture variant-driven splicing changes that are observed experimentally. These results illustrate that models can help bridge gaps in mutagenesis efforts but are not yet a substitute for direct experimental measurement.

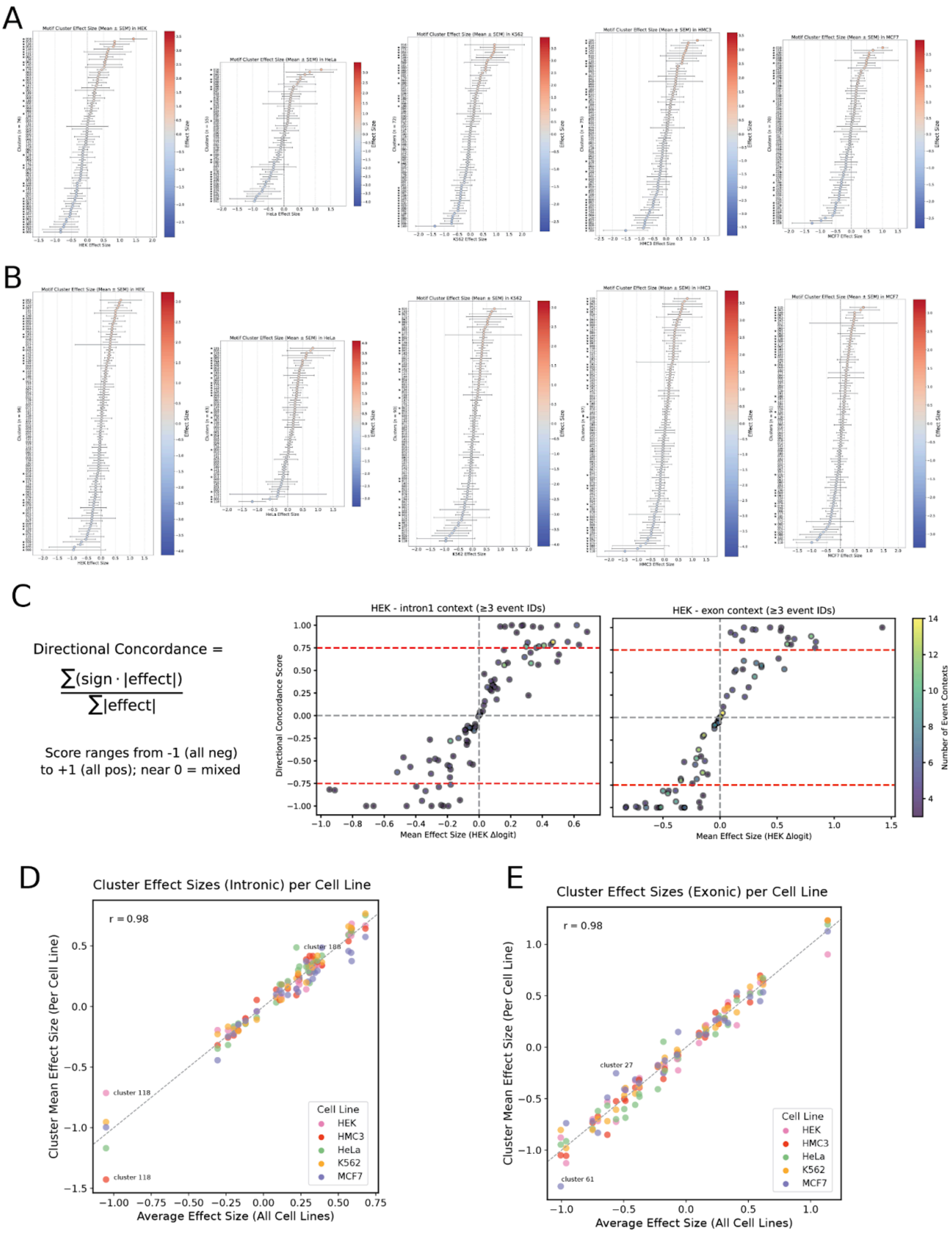
A
B
C
Directional Concordance =
∑(sign · |effect|) / ∑|effect|
Score ranges from -1 (all neg) to +1 (all pos); near 0 = mixed
HEK - intron1 context (≥3 event IDs)
HEK - exon context (≥3 event IDs)
Directional Concordance Score
Mean Effect Size (HEK Δlogit)
Number of Event Contexts
D
Cluster Effect Sizes (Intronic) per Cell Line
r = 0.98
cluster 188
cluster 118
cluster 118
Cluster Mean Effect Size (Per Cell Line)
Average Effect Size (All Cell Lines)
Cell Line
HEK
HMC3
HeLa
K562
MCF7
E
Cluster Effect Sizes (Exonic) per Cell Line
r = 0.98
cluster 27
cluster 61
Cluster Mean Effect Size (Per Cell Line)
Average Effect Size (All Cell Lines)
Cell Line
HEK
HMC3
HeLa
K562
MCF7

**Figure S5 | Concordant RBP motif effect sizes in exonic and intronic regions.**
(A) Exonic region motif effect sizes across five human cell lines (HEK293, HeLa, K562, MCF7, and HMC3). Motif clusters with effect size estimates in at least three exon families are shown. Each point represents the mean Δlogit(PSI) across exon families in a given cell line, with error bars denoting the SEM. Clusters marked with a star pass the concordance filter (≥75% agreement in effect direction across exon families).
(B) Intronic regions. The same analysis as in (A) was applied to motif clusters located in intronic regions.
(C) Concordance filtering of motif effect sizes in HEK293 as an example cell line. This analysis was applied to all cell lines. Motif clusters were included in this analysis if effect sizes were measurable in at least three exon families within each region (intron 1 on the left, exon on the right) for that cell line. Each point represents a motif cluster, with the point color indicating the number of exon families contributing to the calculation. Directional concordance was quantified as the sum of signed effect sizes divided by the sum of absolute effect sizes across exon families. Clusters with strongly correlated effects across exon families approach values of –1 or 1, whereas clusters with weak or inconsistent effects approach 0. Here, we applied a filter of |concordance| ≥ 0.75 to highlight motif clusters with strong and consistent effects across exon families.
(D) Exonic regions. To assess differences across cell types, the mean effect size of each motif cluster was calculated by averaging the effect sizes for a motif cluster across all cell lines (x-axis), and comparing this to the individual cluster means within each cell line (y-axis). Motif clusters included here were selected from the subsets that passed concordance filtering in (C) within each cell line. Points deviating from the diagonal indicate potential cell type-specific effects, such as in cluster 118.
(E) Intronic regions. The same analysis as in (D) applied to motif clusters in intronic regions. Across both contexts, effect sizes were highly correlated across cell lines, indicating that this analysis predominantly captures general *cis*-regulatory effects of motifs rather than cell type–specific activity. Points deviating from the diagonal indicate potential cell type-specific effects, such as in clusters 61 or 27.

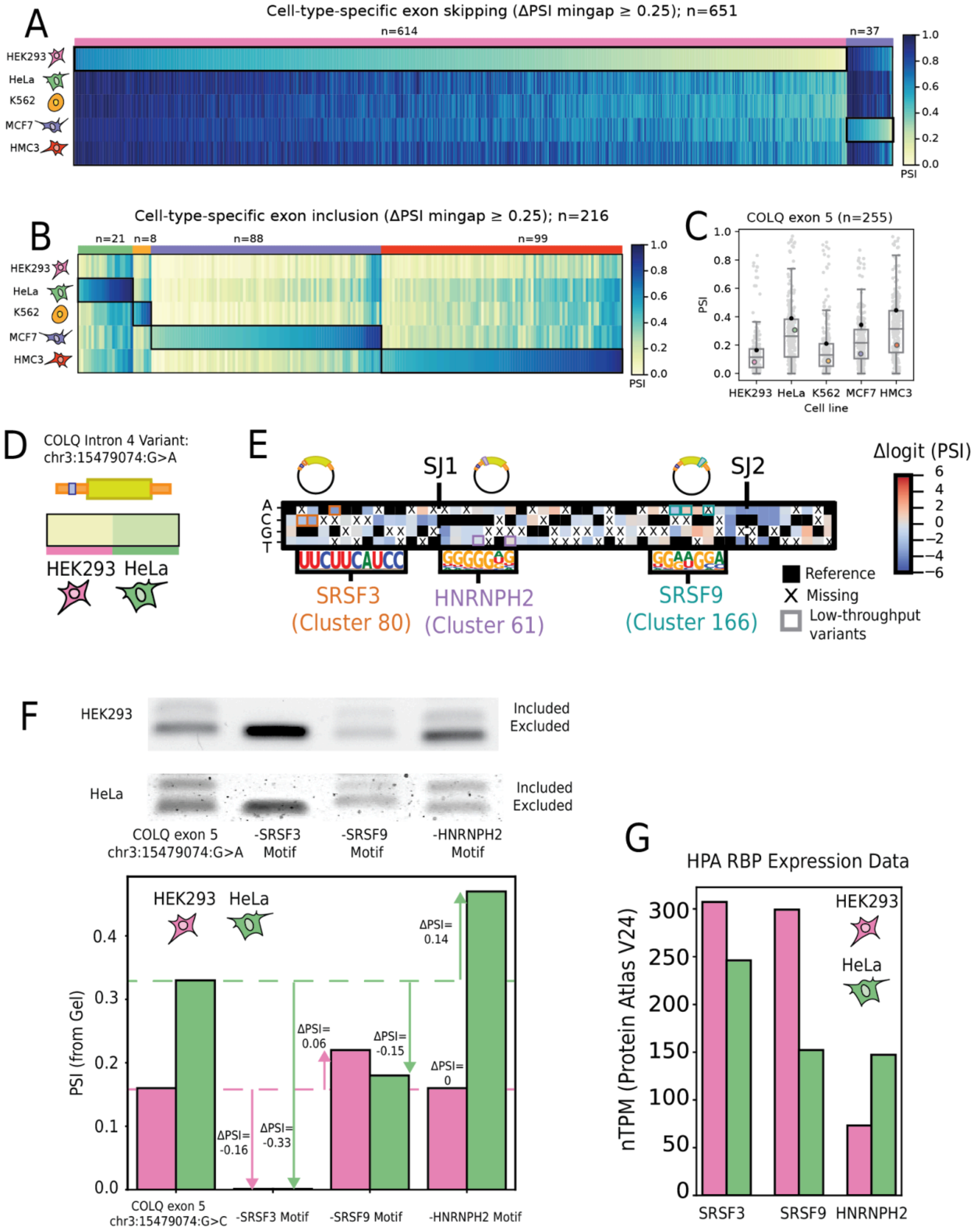
A
Cell-type-specific exon skipping (ΔPSI mingap ≥ 0.25); n=651
n=614
n=37
HEK293
HeLa
K562
MCF7
HMC3
PSI
B
Cell-type-specific exon inclusion (ΔPSI mingap ≥ 0.25); n=216
n=21
n=8
n=88
n=99
C
COLQ exon 5 (n=255)
PSI
Cell line
D
COLQ Intron 4 Variant:
chr3:15479074:G>A
E
SJ1
SJ2
Δlogit (PSI)
SRSF3
(Cluster 80)
HNRNPH2
(Cluster 61)
SRSF9
(Cluster 166)
Reference
Missing
Low-throughput variants
F
Included
Excluded
COLQ exon 5
chr3:15479074:G>A
-SRSF3 Motif
-SRSF9 Motif
-HNRNPH2 Motif
PSI (from Gel)
ΔPSI= 0.14
ΔPSI= 0.06
ΔPSI= -0.15
ΔPSI= 0
ΔPSI= -0.16
ΔPSI= -0.33
COLQ exon 5
chr3:15479074:G>C
G
HPA RBP Expression Data
nTPM (Protein Atlas V24)
SRSF3
SRSF9
HNRNPH2

**Figure S6 | Cell type-specific splicing regulation revealed through differential splicing analysis and motif perturbation.**

**(A)** Heatmap of sequences with cell-type-specific exon inclusion, identified using the mingap $\Delta PSI \geq 0.25$ threshold, defined as an average replicate absolute $\Delta PSI$ between the highest-PSI cell line and the next highest (HeLa, 21; MCF7, 88; HMC3, 99; K562, 8). **(B)** Heatmap of sequences with cell-type-specific exon skipping, identified using the mingap $\Delta PSI \geq 0.25$ threshold, defined as an average replicate absolute $\Delta PSI$ between the lowest-PSI cell line and the next lowest (HEK293, 614; MCF7, 37). **(C)** Swarm plot of COLQ exon 5 PSIs for all cell lines which shows moderate HEK293-HeLa differences. One variant passed the mingap $\Delta PSI \geq 0.25$ threshold and is shown in color. The reference PSI is marked in. black. **(D)** One representative sequence with a variant upstream of COLQ exon 5 was selected for motif disruption experiments based on its large PSI difference (0.16) between HEK293 and HeLa, and a tractable number of candidate motifs. **(E)** NSSM plot of COLQ exon 5 and portions of intron 4. The plot is shown on Δlogit(PSI) scale, with RBP motif binding sites for SRSF3, SRSF9, and HNRNPH2 identified by FIMO. Targeted mutagenesis of each motif was performed in three separate reporters to assess the regulatory and cell type-specific roles of these motifs; mutated regions are highlighted. **(F)** RT-PCR gels showing alternatively spliced isoforms for the original variant (chr3:15479074:G>C) and each of the three motif-disrupting constructs, tested in HEK293 and HeLa. Isoform proportions reflect the splicing impact of each perturbation. PSI values were derived from the quantification of gel band intensities for each construct in both cell types using ImageLab. $\Delta PSI$ values are calculated between each motif-disrupting construct and the original variant (chr3:15479074:G>C), shown separately for HEK293 and HeLa. **(G)** Expression (nTPM) of SRSF3, SRSF9, and HNRNPH2 in HEK293 and HeLa based on data from the Human Protein Atlas (HPA) (version 24).

### Supplementary Tables

| Gene & Exon | Full Sequence (hg38) | Exon Start-End (hg38) | Reference PSI | ΔLogit Min | ΔLogit Max | Number of Variants |
|---|---|---|---|---|---|---|
| ACADVL exon 3 | chr17:7220389-7220549:+ | 7220464-7220529 | 0.2380 | -3.2959 | 4.2370 | 568 |
| ADA exon 7 | chr20:44622987-44623147:- | 44623056-44623127 | 0.6809 | -5.6119 | 2.8908 | 711 |
| ADAM15 exon 20 | chr1:155061349-155061509:+ | 155061418-155061489 | 0.0036 | 0.0000 | 6.3635 | 2094 |
| AFF2 exon 5 | chrX:148837593-148837753:+ | 148837647-148837733 | 0.9515 | -7.9371 | 1.5591 | 530 |
| ALDH6A1 exon 3 | chr14:74072517-74072677:- | 74072583-74072657 | 0.9747 | -9.1901 | 0.7181 | 458 |
| ALDOA exon 7 | chr16:30069249-30069409:+ | 30069306-30069389 | 0.9808 | -8.4104 | 1.2073 | 787 |
| ANK1 exon 41 | chr8:41661856-41662016:- | 41661931-41661996 | 0.9714 | -8.5841 | 1.4020 | 1231 |
| ANK3 exon 23 | chr10:60166571-60166731:- | 60166649-60166711 | 0.2345 | -3.3940 | 5.2235 | 344 |
| AP4M1 exon 3 | chr7:100102606-100102766:+ | 100102675-100102746 | 0.0000 | 0.0000 | 0.0000 | 401 |
| AP4S1 exon 5 | chr14:31080504-31080664:+ | 31080573-31080644 | 0.9802 | -8.6137 | 1.3463 | 412 |
| ASL exon 13 | chr7:66089195-66089355:+ | 66089276-66089335 | 0.1266 | -3.4185 | 5.3007 | 646 |
| ASS1 exon 5 | chr9:130464027-130464187:+ | 130464111-130464167 | 0.9712 | -8.4237 | 1.3523 | 711 |
| AUH exon 9 | chr9:91216039-91216199:- | 91216132-91216179 | 0.6251 | -5.3617 | 4.1595 | 384 |
| BCKDHA exon 3 | chr19:41410869-41411029:+ | 41410923-41411009 | 0.8692 | -6.8148 | 3.4326 | 757 |
| BEST1 exon 8 | chr11:61959438-61959598:+ | 61959498-61959578 | 0.8397 | -6.4202 | 1.9738 | 610 |
| BIN1 exon 16 | chr2:127051134-127051294:- | 127051185-127051274 | 0.2405 | -4.4150 | 4.9946 | 726 |
| BLOC1S6 exon 3 | chr15:45603047-45603207:+ | 45603104-45603187 | 0.1382 | -2.8764 | 4.3883 | 333 |
| CACNA1C exon 10 | chr12:2547384-2547544:+ | 2547450-2547524 | 0.2930 | -4.2775 | 3.8322 | 582 |
| CACNA1C exon 30 | chr12:2634240-2634400:+ | 2634297-2634380 | 0.0000 | 0.0000 | 4.5991 | 544 |
| CACNA1C exon 31 | chr12:2633572-2633732:+ | 2633629-2633712 | 0.8969 | -7.0204 | 2.7124 | 518 |
| CACNA1C exon 31 | chr12:2648367-2648527:+ | 2648475-2648507 | 0.0017 | 0.0000 | 1.5853 | 474 |
| CACNA2D4 exon 28 | chr12:1810523-1810683:- | 1810619-1810663 | 0.4584 | -4.5835 | 4.1865 | 724 |
| CASK exon 19 | chrX:41557012-41557172:- | 41557084-41557152 | 0.9231 | -7.4050 | 1.8500 | 318 |
| CLN3 exon 5 | chr16:28488571-28488731:- | 28488640-28488711 | 0.9355 | -7.7806 | 2.4959 | 1062 |
| COL11A2 exon 6 | chr6:33185681-33185841:- | 33185744-33185821 | 0.0014 | 0.0000 | 1.9548 | 947 |
| COL11A2 exon 7 | chr6:33184972-33185132:- | 33185050-33185112 | 0.5634 | -4.8372 | 4.3156 | 1838 |
| COLQ exon 5 | chr3:15478957-15479117:- | 15479071-15479097 | 0.3083 | -3.7620 | 3.2489 | 791 |
| DEPDC5 exon 33 | chr22:31861293-31861453:+ | 31861368-31861433 | 0.9128 | -7.0807 | 2.0162 | 852 |
| DEPDC5 exon 6 | chr22:31766528-31766688:+ | 31766585-31766668 | 0.9287 | -7.4663 | 1.7239 | 411 |
| DMD exon 71 | chrX:31177912-31178072:- | 31178014-31178052 | 0.9712 | -8.2383 | 1.0030 | 375 |
| DNM1L exon 15 | chr12:32737802-32737962:+ | 32737865-32737942 | 0.0828 | -2.8745 | 5.4149 | 697 |
| DNM1L exon 4 | chr12:32708084-32708244:+ | 32708153-32708224 | 0.9452 | -7.8214 | 1.5542 | 351 |
| DNM1L exon 5 | chr12:32710875-32711035:+ | 32710929-32711015 | 0.9708 | -8.2392 | 1.8745 | 316 |
| DRD2 exon 6 | chr11:113414355-113414515:- | 113414409-113414495 | 0.9836 | -8.7065 | 1.4388 | 788 |
| DYSF exon 17 | chr2:71549251-71549411:+ | 71549350-71549391 | 0.1158 | -2.4376 | 5.9357 | 860 |
| ECEL1 exon 15 | chr2:232481071-232481231:- | 232481146-232481211 | 0.0008 | 0.0000 | 1.1710 | 722 |
| ELN exon 13 | chr7:74047576-74047736:+ | 74047675-74047716 | 0.3734 | -4.3715 | 4.7629 | 541 |
| ELN exon 24 | chr7:74060044-74060204:+ | 74060140-74060184 | 0.0489 | -1.9645 | 6.9501 | 806 |
| ELN exon 5 | chr7:74041111-74041271:+ | 74041216-74041251 | 0.4799 | -4.6034 | 4.3761 | 810 |
| EPB41L1 exon 13 | chr20:36195224-36195384:+ | 36195329-36195364 | 0.3811 | -4.0858 | 4.0838 | 688 |
| EPB41L1 exon 18 | chr20:36218822-36218982:+ | 36218876-36218962 | 0.9650 | -8.1927 | 0.9764 | 756 |
| ERCC2 exon 3 | chr19:45369050-45369210:- | 45369113-45369190 | 0.4489 | -5.2168 | 4.5096 | 679 |
| FANCA exon 38 | chr16:89740784-89740944:- | 89740862-89740924 | 0.9823 | -8.6962 | 0.4906 | 302 |
| FGF8 exon 3 | chr10:101775110-101775270:- | 101775164-101775250 | 0.0006 | 0.0000 | 2.4157 | 1548 |
| GALC exon 2 | chr14:87988435-87988595:- | 87988507-87988575 | 0.9749 | -8.4716 | 0.8123 | 359 |
| GCDH exon 5 | chr19:12892038-12892198:+ | 12892116-12892178 | 0.0717 | -2.2080 | 6.1914 | 603 |
| GDAP1 exon 2 | chr8:74351181-74351341:+ | 74351274-74351321 | 0.9380 | -8.1156 | 1.1216 | 347 |
| GH1 exon 3 | chr17:63917997-63918157:- | 63918063-63918137 | 0.0078 | -0.4091 | 2.4319 | 352 |
| GNAS exon 3 | chr20:58898845-58899005:+ | 58898941-58898985 | 0.1956 | -3.3078 | 4.8470 | 1220 |
| GNS exon 2 | chr12:64752678-64752838:- | 64752759-64752818 | 0.9831 | -8.7494 | 0.4521 | 364 |
| GPI exon 5 | chr19:34377446-34377606:+ | 34377503-34377586 | 0.8374 | -6.5666 | 2.4973 | 767 |
| GUSB exon 3 | chr7:65979814-65979974:- | 65979877-65979954 | 0.0002 | 0.0000 | 3.7914 | 760 |
| HARS2 exon 2 | chr5:140693525-140693685:+ | 140693591-140693665 | 0.4898 | -4.6065 | 3.7368 | 533 |
| HMBS exon 10 | chr11:119092023-119092183:+ | 119092125-119092163 | 0.0533 | -2.2328 | 4.6302 | 803 |
| HMBS exon 2 | chr11:119088168-119088328:+ | 119088255-119088308 | 0.4738 | -4.5249 | 2.8189 | 834 |
| HMBS exon 6 | chr11:119089620-119089780:+ | 119089683-119089760 | 0.3941 | -4.1780 | 4.7774 | 421 |
| IL2RA exon 5 | chr10:6019850-6020010:- | 6019919-6019990 | 0.9777 | -8.5023 | 1.5411 | 400 |
| IMPDH1 exon 4 | chr7:128400812-128400972:- | 128400893-128400952 | 0.0001 | 0.0000 | 2.0067 | 769 |
| IMPDH1 exon 7 | chr7:128400797-128400957:- | 128400863-128400937 | 0.0250 | -1.7823 | 4.7588 | 706 |

| | | | | | | |
|---|---|---|---|---|---|---|
| IVD exon 2 | chr15:40407585-40407745:+ | 40407636-40407725 | 0.5111 | -5.3611 | 3.8292 | 745 |
| KCNQ2 exon 9 | chr20:63431320-63431480:- | 63431431-63431460 | 0.0003 | 0.0000 | 2.0343 | 440 |
| L1CAM exon 3 | chrX:153873208-153873368:- | 153873334-153873348 | 0.0016 | 0.0000 | 1.6138 | 938 |
| LARS1 exon 4 | chr5:146171890-146172050:- | 146171950-146172030 | 0.9835 | -3.3054 | 0.6748 | 377 |
| LMNA exon 10 | chr1:156137603-156137763:+ | 156137654-156137743 | 0.9665 | -8.2386 | 1.0171 | 839 |
| MAPT exon 9 | chr17:45993837-45993997:+ | 45993924-45993977 | 0.0010 | 0.0000 | 7.8363 | 573 |
| MEF2C exon 5 | chr5:88761249-88761409:- | 88761330-88761389 | 0.0006 | 0.0000 | 7.5122 | 636 |
| MEF2C exon 8 | chr5:88730191-88730351:- | 88730308-88730331 | 0.0150 | -0.9271 | 3.8643 | 433 |
| MEFV exon 8 | chr16:3244234-3244394:- | 3244342-3244374 | 0.6251 | -5.1136 | 3.0179 | 730 |
| MFAP5 exon 3 | chr12:8660843-8661003:- | 8660948-8660983 | 0.9529 | -7.9262 | 1.5937 | 430 |
| MLC1 exon 3 | chr22:50083064-50083224:- | 50083115-50083204 | 0.9727 | -8.4516 | 1.1937 | 628 |
| MLC1 exon 4 | chr22:50080324-50080484:- | 50080411-50080464 | 0.9206 | -7.1217 | 1.9813 | 860 |
| MUTYH exon 6 | chr1:45332898-45333058:- | 45332997-45333038 | 0.0034 | 0.0000 | 5.5971 | 774 |
| MYBPC1 exon 4 | chr12:101626770-101626930:+ | 101626872-101626910 | 0.9240 | -7.2552 | 1.8918 | 748 |
| MYH11 exon 42 | chr16:15708783-15708943:- | 15708885-15708923 | 0.6108 | -5.4515 | 3.7273 | 698 |
| MYH11 exon 6 | chr16:15784678-15784838:- | 15784798-15784818 | 0.0028 | -0.0093 | 0.5977 | 430 |
| NDUFAF5 exon 9 | chr20:13816406-13816566:+ | 13816463-13816546 | 0.8672 | -6.8340 | 2.3726 | 638 |
| NEK1 exon 20 | chr4:169508749-169508909:- | 169508806-169508889 | 0.3742 | -4.2250 | 4.3460 | 311 |
| NF1 exon 31 | chr17:31252860-31253020:+ | 31252938-31253000 | 0.9057 | -6.9783 | 2.3792 | 445 |
| NLRC3 exon 14 | chr16:3548650-3548810:- | 3548707-3548790 | 0.7144 | -5.6814 | 2.7336 | 438 |
| NPC2 exon 4 | chr14:74480682-74480842:- | 74480745-74480822 | 0.9855 | -8.3815 | 1.2232 | 373 |
| NRXN1 exon 8 | chr2:50621185-50621345:- | 50621281-50621325 | 0.0000 | 0.0000 | 2.9576 | 313 |
| OCA2 exon 10 | chr15:27990556-27990716:- | 27990625-27990696 | 0.9203 | -7.6589 | 1.8333 | 706 |
| PAX5 exon 9 | chr9:36846823-36846983:- | 36846877-36846963 | 0.9819 | -8.7112 | 1.4406 | 1724 |
| PCCA exon 2 | chr13:100102820-100102980:+ | 100102883-100102960 | 0.9797 | -8.6153 | 1.3080 | 324 |
| PDE6B exon 8 | chr4:656152-656312:+ | 656245-656292 | 0.5490 | -5.3385 | 3.9616 | 660 |
| PIGH exon 3 | chr14:67592615-67592775:- | 67592675-67592755 | 0.9111 | -7.1631 | 2.0262 | 507 |
| PPT1 exon 8 | chr1:40076822-40076982:- | 40076891-40076962 | 0.9756 | -8.4246 | 1.7714 | 402 |
| RAB7A exon 4 | chr3:128806309-128806469:+ | 128806372-128806449 | 0.9500 | -8.8193 | 1.3547 | 745 |
| RYR1 exon 94 | chr19:38570553-38570713:+ | 38570607-38570693 | 0.1087 | -3.0521 | 4.2817 | 572 |
| SCN5A exon 24 | chr3:38557211-38557371:- | 38557298-38557351 | 0.8511 | -6.4295 | 2.8010 | 577 |
| SERPINB7 exon 3 | chr18:63792303-63792463:+ | 63792393-63792443 | 0.9807 | -8.6203 | 1.3284 | 424 |
| SHOX2 exon 2 | chr3:158105038-158105198:- | 158105107-158105178 | 0.0477 | -2.1311 | 7.6068 | 1010 |
| SLC13A5 exon 2 | chr17:6707008-6707168:- | 6707131-6707148 | 0.0680 | -1.6083 | 5.4041 | 304 |
| SLC26A4 exon 11 | chr7:107694340-107694500:+ | 107694403-107694480 | 0.6373 | -5.1567 | 3.3187 | 557 |
| SLC26A5 exon 12 | chr7:103390409-103390569:- | 103390472-103390549 | 0.7272 | -5.6380 | 3.8558 | 555 |
| SNCA exon 3 | chr4:89828123-89828283:- | 89828222-89828263 | 0.9843 | -9.1902 | 0.6244 | 417 |
| SOS1 exon 21 | chr2:38989250-38989410:- | 38989346-38989390 | 0.1462 | -2.7697 | 5.6752 | 365 |
| SPINT2 exon 3 | chr19:38287795-38287955:+ | 38287876-38287935 | 0.9704 | -8.4735 | 1.1078 | 548 |
| STXBP2 exon 13 | chr19:7643105-7643265:+ | 7643165-7643245 | 0.0017 | 0.0000 | 4.6999 | 946 |
| SUMF1 exon 3 | chr3:4449246-4449406:- | 4449312-4449386 | 0.9637 | -8.1277 | 1.6723 | 326 |
| SUMF1 exon 8 | chr3:4376310-4376470:- | 4376391-4376450 | 0.9880 | -7.4389 | 0.9993 | 354 |
| SYNGAP1 exon 14 | chr6:33442354-33442514:+ | 33442453-33442494 | 0.0016 | 0.0000 | 3.0971 | 762 |
| TAC3 exon 5 | chr12:57012802-57012962:- | 57012889-57012942 | 0.9791 | -8.9206 | 0.9203 | 641 |
| TIMM44 exon 11 | chr19:7928057-7928217:- | 7928108-7928197 | 0.0547 | -2.6541 | 4.7784 | 784 |
| TNFRSF1A exon 3 | chr12:6333825-6333985:- | 6333945-6333965 | 0.0000 | 0.0000 | 0.0000 | 636 |
| TNNT2 exon 4 | chr1:201372007-201372167:- | 201372133-201372147 | 0.8881 | -6.6186 | 2.8294 | 512 |
| TNNT2 exon 5 | chr1:201369796-201369956:- | 201369907-201369936 | 0.1645 | -2.9302 | 4.5015 | 842 |
| TSC2 exon 32 | chr16:2082364-2082524:+ | 2082436-2082504 | 0.0059 | -0.4247 | 2.9869 | 874 |
| TTC8 exon 7 | chr14:88843710-88843870:+ | 88843806-88843850 | 0.7465 | -5.6842 | 3.3308 | 388 |
| TTN-AS1 exon 4 | chr2:178537307-178537467:+ | 178537361-178537447 | 0.0000 | 0.0000 | 0.0000 | 441 |
| UROS exon 3 | chr10:125816157-125816317:- | 125816214-125816297 | 0.9598 | -8.7013 | 1.1258 | 421 |
| USB1 exon 4 | chr16:58014186-58014346:+ | 58014273-58014326 | 0.9295 | -7.8930 | 1.8662 | 351 |
| USP28 exon 6 | chr11:113834229-113834389:- | 113834283-113834369 | 0.3645 | -4.1615 | 4.2147 | 449 |
| VPS33B exon 2 | chr15:91017785-91017945:- | 91017845-91017925 | 0.9798 | -8.5880 | 0.8321 | 532 |
| WDR35 exon 11 | chr2:19962275-19962435:- | 19962383-19962415 | 0.0025 | 0.0000 | 4.1151 | 332 |

**Supplementary Table 1** | Summary of reference PSI and Δlogit(PSI) ranges for near-saturation exons.
This table reports all exons with at least 300 assayed variants. For each exon we provide the gene and exon number, event identifier with the genomic coordinates (hg38) of the variable sequence, coordinates of the exon start and stop, the mean reference PSI calculated across all cell lines, and the minimum and maximum average Δlogit(PSI) across all cell lines. The minimum and maximum is determined across all variants for that exon. We also report the total number of variants observed for that exon. These data summarize the splicing landscape of highly covered exons and serve as a reference for interpreting mutational effects in this study.
