## Supplementary material for "Massively parallel assay of human splice variants reveals cis-regulatory drivers of disease-associated and cell type-specific splicing regulation": Data S1

A

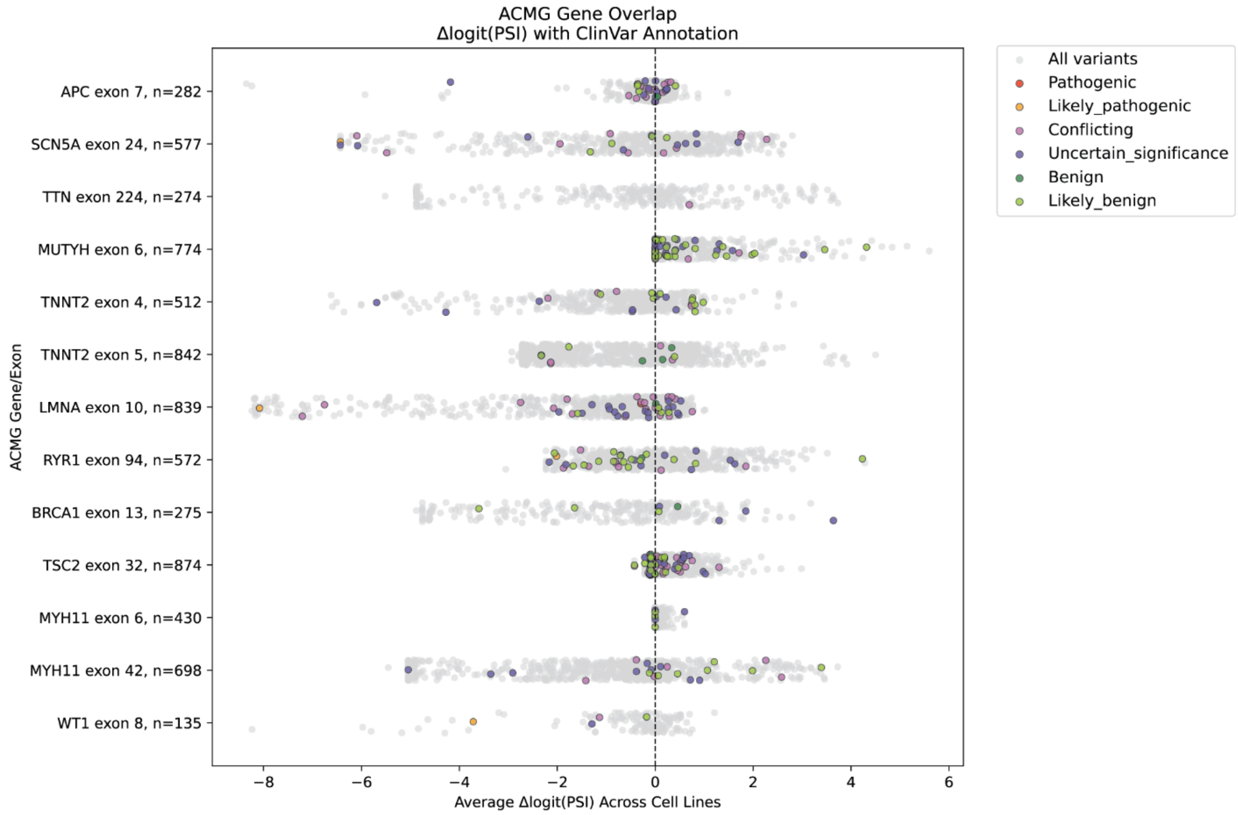
ACMG Gene Overlap
Δlogit(PSI) with ClinVar Annotation
APC exon 7, n=282
SCN5A exon 24, n=577
TTN exon 224, n=274
MUTYH exon 6, n=774
TNNT2 exon 4, n=512
TNNT2 exon 5, n=842
LMNA exon 10, n=839
RYR1 exon 94, n=572
BRCA1 exon 13, n=275
TSC2 exon 32, n=874
MYH11 exon 6, n=430
MYH11 exon 42, n=698
WT1 exon 8, n=135
ACMG Gene/Exon
Average Δlogit(PSI) Across Cell Lines
−8
−6
−4
−2
0
2
4
6
All variants
Pathogenic
Likely_pathogenic
Conflicting
Uncertain_significance
Benign
Likely_benign

B

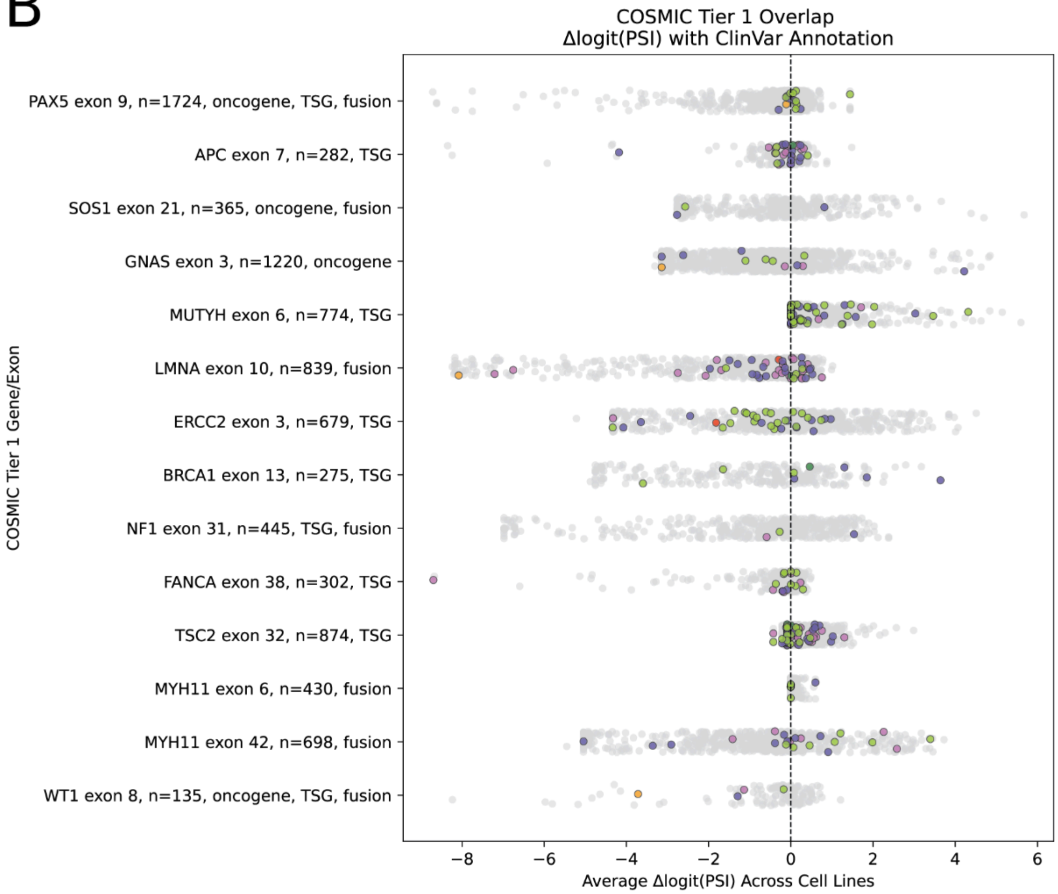
COSMIC Tier 1 Overlap
Δlogit(PSI) with ClinVar Annotation
PAX5 exon 9, n=1724, oncogene, TSG, fusion
APC exon 7, n=282, TSG
SOS1 exon 21, n=365, oncogene, fusion
GNAS exon 3, n=1220, oncogene
MUTYH exon 6, n=774, TSG
LMNA exon 10, n=839, fusion
ERCC2 exon 3, n=679, TSG
BRCA1 exon 13, n=275, TSG
NF1 exon 31, n=445, TSG, fusion
FANCA exon 38, n=302, TSG
TSC2 exon 32, n=874, TSG
MYH11 exon 6, n=430, fusion
MYH11 exon 42, n=698, fusion
WT1 exon 8, n=135, oncogene, TSG, fusion
COSMIC Tier 1 Gene/Exon
Average Δlogit(PSI) Across Cell Lines
−8
−6
−4
−2
0
2
4
6

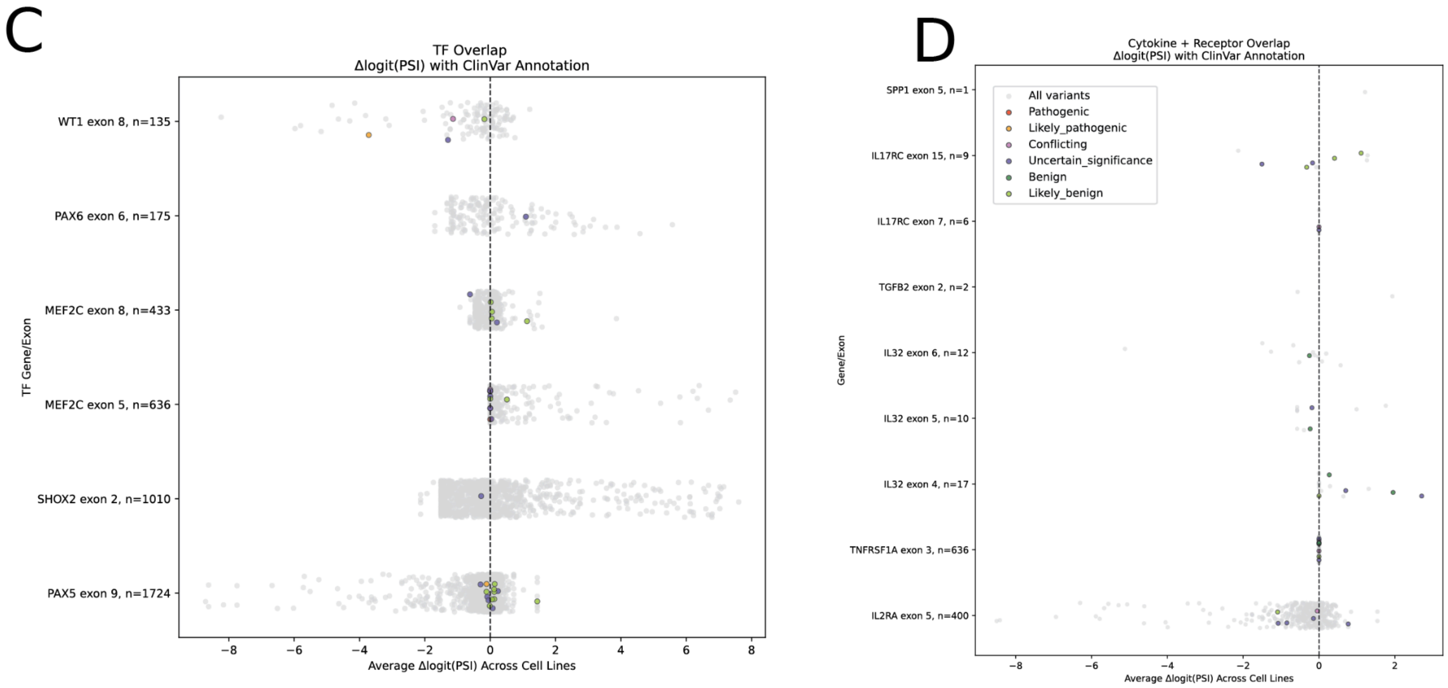


**Δlogit(PSI) distributions for variants in clinically and biologically important gene sets.**
**(A)** Variants from genes in the ACMG SF v3.2 list.[1] Of the 81 genes in this set, 13 exons from 11 genes were represented in our dataset with at least 100 variants measured in our MPRA. Distributions of variant Δlogit(PSI) are shown, with ClinVar-annotated SNVs highlighted.
**(B)** Exons from COSMIC Cancer Gene Census (CGC) Tier 1 genes that were represented in our MPRA with at least 100 measured variants are shown.[2] Δlogit(PSI) distributions are shown with ClinVar annotations indicated, and genes further classified by functional category (tumor suppressor genes, oncogenes, or gene fusions). Tier 1 genes have strong evidence of cancer relevance from both mutational and functional studies.
**(C)** Exons from six transcription factor (TF) genes, overlapping a catalog of 1,639 known and likely human TFs and their motifs, were represented in our MPRA with at least 100 measured variants. Distributions of variant Δlogit(PSI) values are shown.[3]
**(D)** Variants from 9 exons belonging to cytokines and cytokine receptors, as defined in the ImmPort Cytokine registry. Δlogit(PSI) distributions are shown. [4]

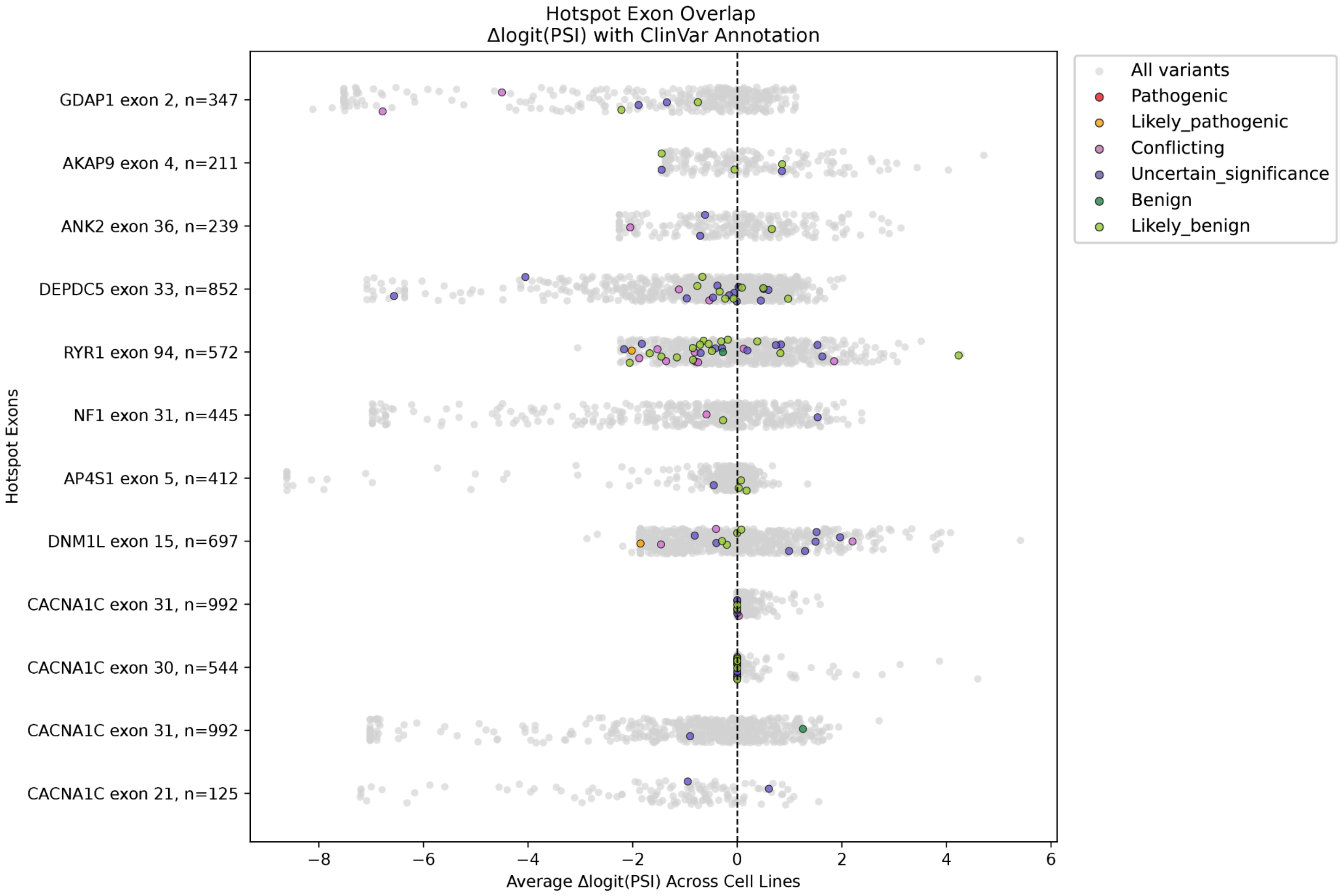


**Δlogit(PSI) distributions for variants in mutational hotspot exons.**

Of the exons classified as hotspot exons in the HEK293T, 45 of these exons from 42 genes were represented in our dataset.[5] Distributions of variant Δlogit(PSI) are shown for 12 of these hotspot exons that have at least 100 variants measured in our MPRA. ClinVar-annotated SNVs are highlighted. Hotspot exons are common targets of splicing perturbations, consistent with their enrichment for SDVs in our MPRA.

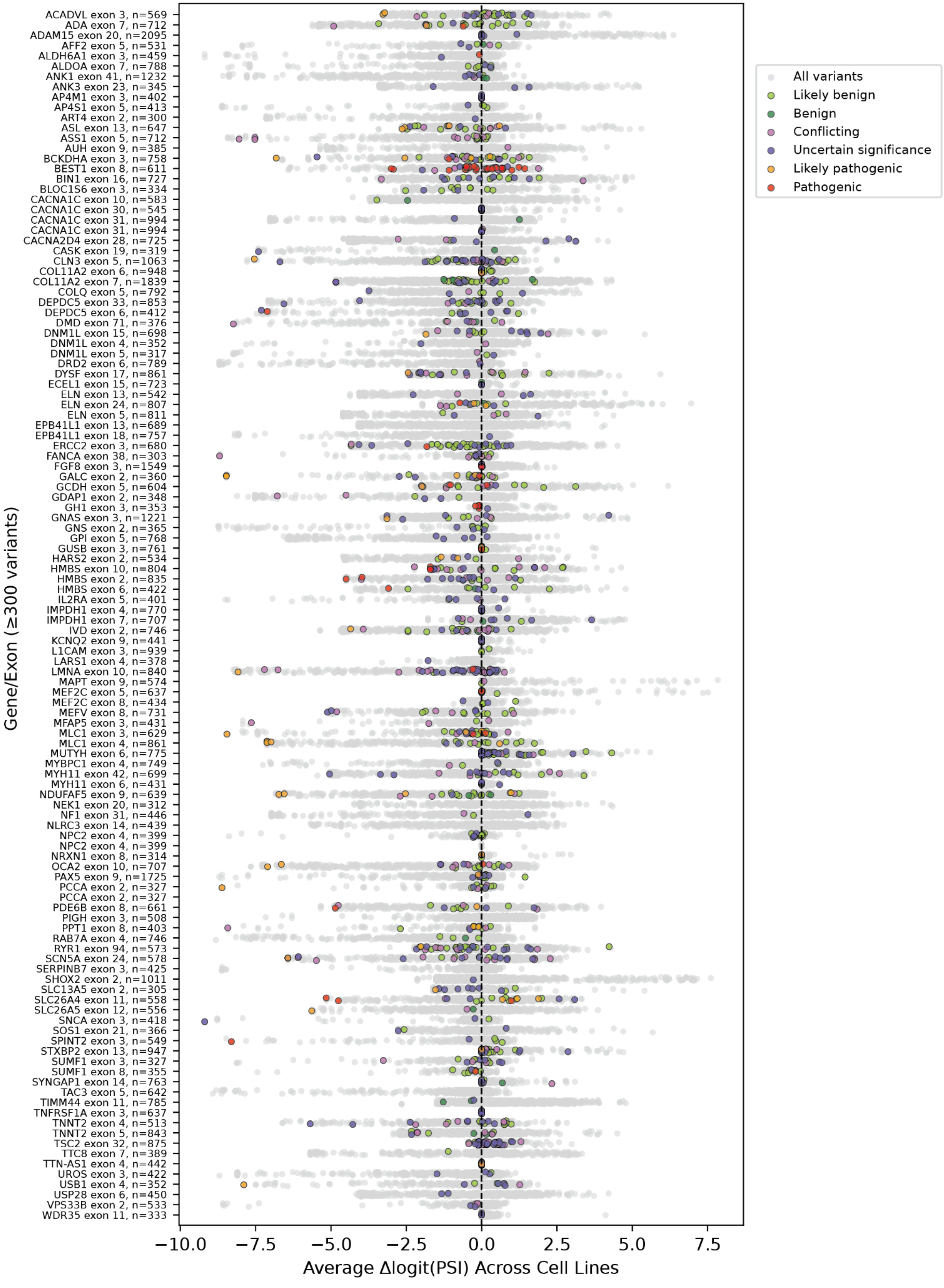
ACADVL exon 3, n=569
ADA exon 7, n=712
ADAM15 exon 20, n=2095
AFF2 exon 5, n=531
ALDH6A1 exon 3, n=459
ALDOA exon 7, n=788
ANK1 exon 41, n=1232
ANK3 exon 23, n=345
AP4M1 exon 3, n=402
AP4S1 exon 5, n=413
ART4 exon 2, n=300
ASL exon 13, n=647
ASS1 exon 5, n=712
AUH exon 9, n=385
BCKDHA exon 3, n=758
BEST1 exon 8, n=611
BIN1 exon 16, n=727
BLOC1S6 exon 3, n=334
CACNA1C exon 10, n=583
CACNA1C exon 30, n=545
CACNA1C exon 31, n=994
CACNA1C exon 31, n=994
CACNA2D4 exon 28, n=725
CASK exon 19, n=319
CLN3 exon 5, n=1063
COL11A2 exon 6, n=948
COL11A2 exon 7, n=1839
COLQ exon 5, n=792
DEPDC5 exon 33, n=853
DEPDC5 exon 6, n=412
DMD exon 71, n=376
DNM1L exon 15, n=698
DNM1L exon 4, n=352
DNM1L exon 5, n=317
DRD2 exon 6, n=789
DYSF exon 17, n=861
ECEL1 exon 15, n=723
ELN exon 13, n=542
ELN exon 24, n=807
ELN exon 5, n=811
EPB41L1 exon 13, n=689
EPB41L1 exon 18, n=757
ERCC2 exon 3, n=680
FANCA exon 38, n=303
FGF8 exon 3, n=1549
GALC exon 2, n=360
GCDH exon 5, n=604
GDAP1 exon 2, n=348
GH1 exon 3, n=353
GNAS exon 3, n=1221
GNS exon 2, n=365
GPI exon 5, n=768
GUSB exon 3, n=761
HARS2 exon 2, n=534
HMBS exon 10, n=804
HMBS exon 2, n=835
HMBS exon 6, n=422
IL2RA exon 5, n=401
IMPDH1 exon 4, n=770
IMPDH1 exon 7, n=707
IVD exon 2, n=746
KCNQ2 exon 9, n=441
L1CAM exon 3, n=939
LARS1 exon 4, n=378
LMNA exon 10, n=840
MAPT exon 9, n=574
MEF2C exon 5, n=637
MEF2C exon 8, n=434
MEFV exon 8, n=731
MFAP5 exon 3, n=431
MLC1 exon 3, n=629
MLC1 exon 4, n=861
MUTYH exon 6, n=775
MYBPC1 exon 4, n=749
MYH11 exon 42, n=699
MYH11 exon 6, n=431
NDUFAF5 exon 9, n=639
NEK1 exon 20, n=312
NF1 exon 31, n=446
NLRC3 exon 14, n=439
NPC2 exon 4, n=399
NPC2 exon 4, n=399
NRXN1 exon 8, n=314
OCA2 exon 10, n=707
PAX5 exon 9, n=1725
PCCA exon 2, n=327
PCCA exon 2, n=327
PDE6B exon 8, n=661
PIGH exon 3, n=508
PPT1 exon 8, n=403
RAB7A exon 4, n=746
RYR1 exon 94, n=573
SCN5A exon 24, n=578
SERPINB7 exon 3, n=425
SHOX2 exon 2, n=1011
SLC13A5 exon 2, n=305
SLC26A4 exon 11, n=558
SLC26A5 exon 12, n=556
SNCA exon 3, n=418
SOS1 exon 21, n=366
SPINT2 exon 3, n=549
STXBP2 exon 13, n=947
SUMF1 exon 3, n=327
SUMF1 exon 8, n=355
SYNGAP1 exon 14, n=763
TAC3 exon 5, n=642
TIMM44 exon 11, n=785
TNFRSF1A exon 3, n=637
TNNT2 exon 4, n=513
TNNT2 exon 5, n=843
TSC2 exon 32, n=875
TTC8 exon 7, n=389
TTN-AS1 exon 4, n=442
UROS exon 3, n=422
USB1 exon 4, n=352
USP28 exon 6, n=450
VPS33B exon 2, n=533
WDR35 exon 11, n=333
Gene/Exon (≥300 variants)
−10.0
−7.5
−5.0
−2.5
0.0
2.5
5.0
7.5
Average Δlogit(PSI) Across Cell Lines
All variants
Likely benign
Benign
Conflicting
Uncertain significance
Likely pathogenic
Pathogenic

**Δlogit(PSI) distributions for highly saturated exons.**

A subset of 115 exons from the MPRA with high mutational saturation, each represented by at least 300 variants (single and double nucleotide substitutions). This subset illustrates the dense mutational coverage achieved, approaching saturation and providing context into splicing effects when perturbing nearly all of the cis-regulatory elements in each exon.
