## Supplementary material for "Massively parallel assay of human splice variants reveals cis-regulatory drivers of disease-associated and cell type-specific splicing regulation": Data S2

### Motif Effects Across Exon Contexts (Mean ± SEM, All Cell Lines)

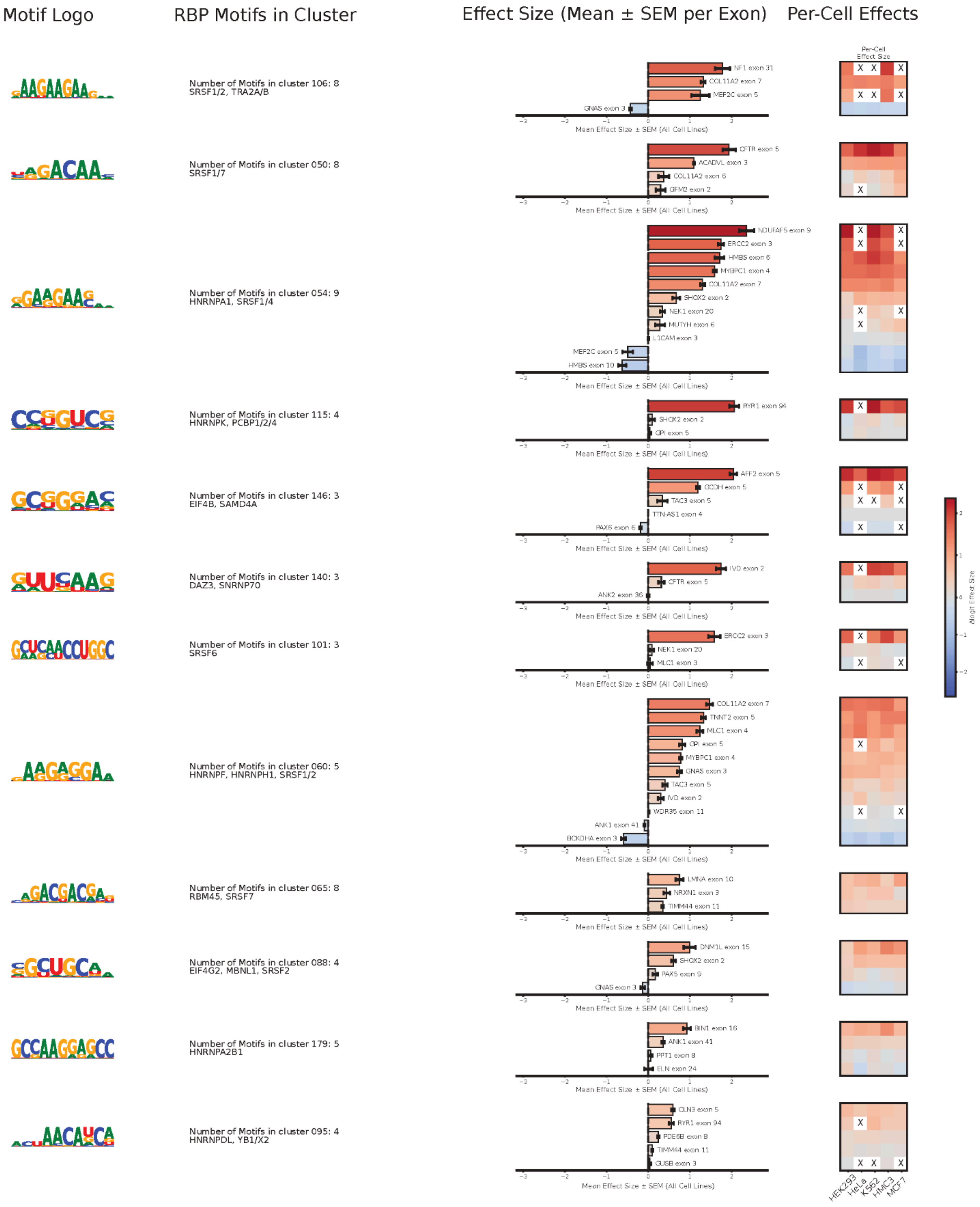

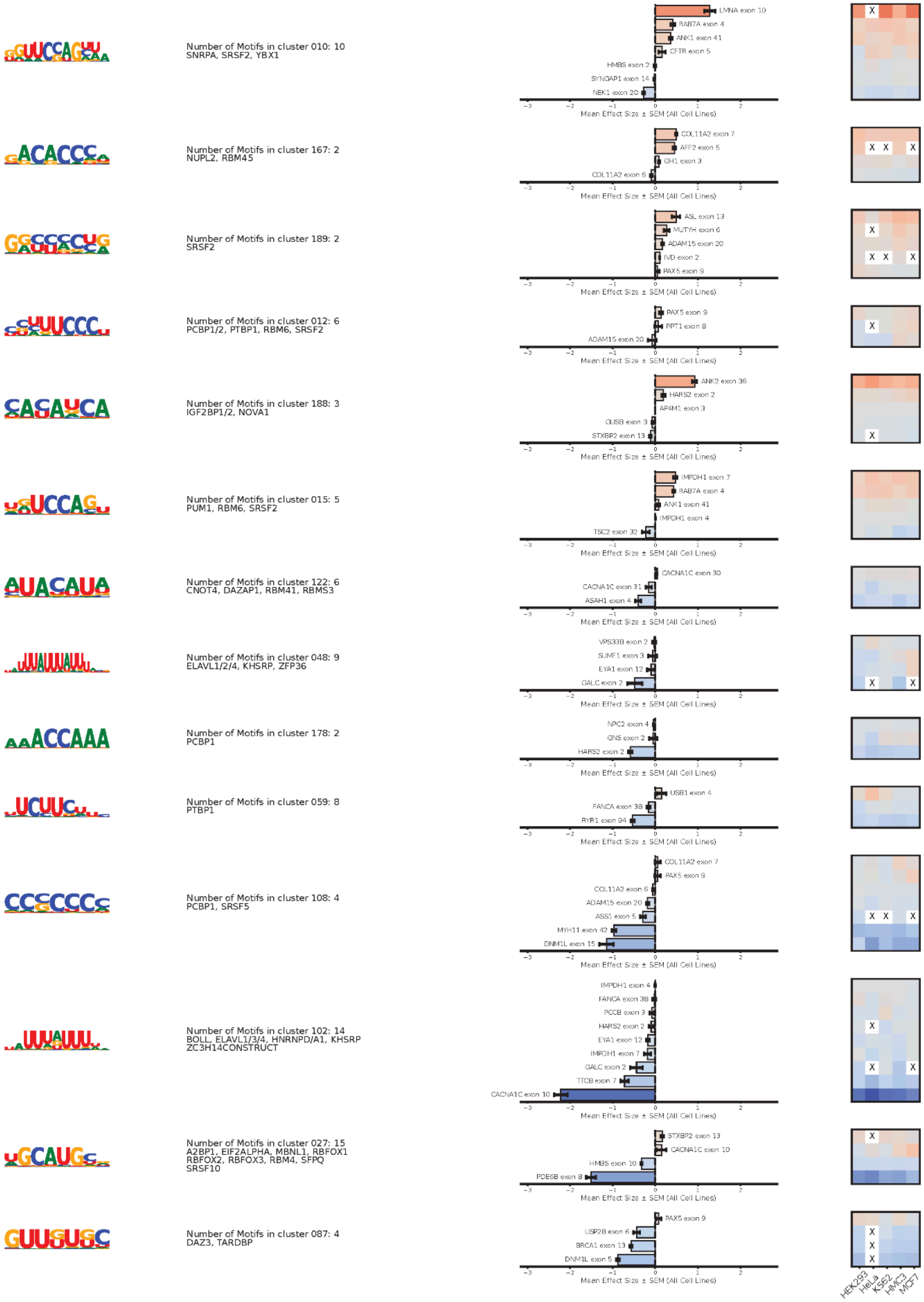

Number of Motifs in cluster 010: 10
SNRPA, SRSF2, YBX1
LMNA exon 10
RAB7A exon 4
ANK1 exon 41
CFTR exon 5
HMBS exon 2
SYNGAP1 exon 14
NEK1 exon 20
Mean Effect Size ± SEM (All Cell Lines)
Number of Motifs in cluster 167: 2
NUPL2, RBM45
COL11A2 exon 7
AFF2 exon 5
GH1 exon 3
COL11A2 exon 6
Mean Effect Size ± SEM (All Cell Lines)
Number of Motifs in cluster 189: 2
SRSF2
ASL exon 13
MUTYH exon 6
ADAM15 exon 20
IVD exon 2
PAX5 exon 9
Mean Effect Size ± SEM (All Cell Lines)
Number of Motifs in cluster 012: 6
PCBP1/2, PTBP1, RBM6, SRSF2
PAX5 exon 9
PPT1 exon 8
ADAM15 exon 20
Mean Effect Size ± SEM (All Cell Lines)
Number of Motifs in cluster 188: 3
IGF2BP1/2, NOVA1
ANK2 exon 36
HARS2 exon 2
AP4M1 exon 3
GUSB exon 3
STXBP2 exon 13
Mean Effect Size ± SEM (All Cell Lines)
Number of Motifs in cluster 015: 5
PUM1, RBM6, SRSF2
IMPDH1 exon 7
RAB7A exon 4
ANK1 exon 41
IMPDH1 exon 4
TSC2 exon 32
Mean Effect Size ± SEM (All Cell Lines)
Number of Motifs in cluster 122: 6
CNOT4, DAZAP1, RBM41, RBMS3
CACNA1C exon 30
CACNA1C exon 31
ASAH1 exon 4
Mean Effect Size ± SEM (All Cell Lines)
Number of Motifs in cluster 048: 9
ELAVL1/2/4, KHSRP, ZFP36
VPS33B exon 2
SUMF1 exon 3
EYA1 exon 12
GALC exon 2
Mean Effect Size ± SEM (All Cell Lines)
Number of Motifs in cluster 178: 2
PCBP1
NPC2 exon 4
GNS exon 2
HARS2 exon 2
Mean Effect Size ± SEM (All Cell Lines)
Number of Motifs in cluster 059: 8
PTBP1
USB1 exon 4
FANCA exon 38
RYR1 exon 94
Mean Effect Size ± SEM (All Cell Lines)
Number of Motifs in cluster 108: 4
PCBP1, SRSF5
COL11A2 exon 7
PAX5 exon 9
COL11A2 exon 6
ADAM15 exon 20
ASS1 exon 5
MYH11 exon 42
DNM1L exon 15
Mean Effect Size ± SEM (All Cell Lines)
Number of Motifs in cluster 102: 14
BOLL, ELAVL1/3/4, HNRNPD/A1, KHSRP
ZC3H14CONSTRUCT
IMPDH1 exon 4
FANCA exon 38
PCCB exon 3
HARS2 exon 2
EYA1 exon 12
IMPDH1 exon 7
GALC exon 2
TTC8 exon 7
CACNA1C exon 10
Mean Effect Size ± SEM (All Cell Lines)
Number of Motifs in cluster 027: 15
A2BP1, EIF2ALPHA, MBNL1, RBFOX1
RBFOX2, RBFOX3, RBM4, SFPQ
SRSF10
STXBP2 exon 13
CACNA1C exon 10
HMBS exon 10
PDE6B exon 8
Mean Effect Size ± SEM (All Cell Lines)
Number of Motifs in cluster 087: 4
DAZ3, TARDBP
PAX5 exon 9
USP28 exon 6
BRCA1 exon 13
DNM1L exon 5
Mean Effect Size ± SEM (All Cell Lines)
HEK293
HeLa
K562
HMC3
MCF7

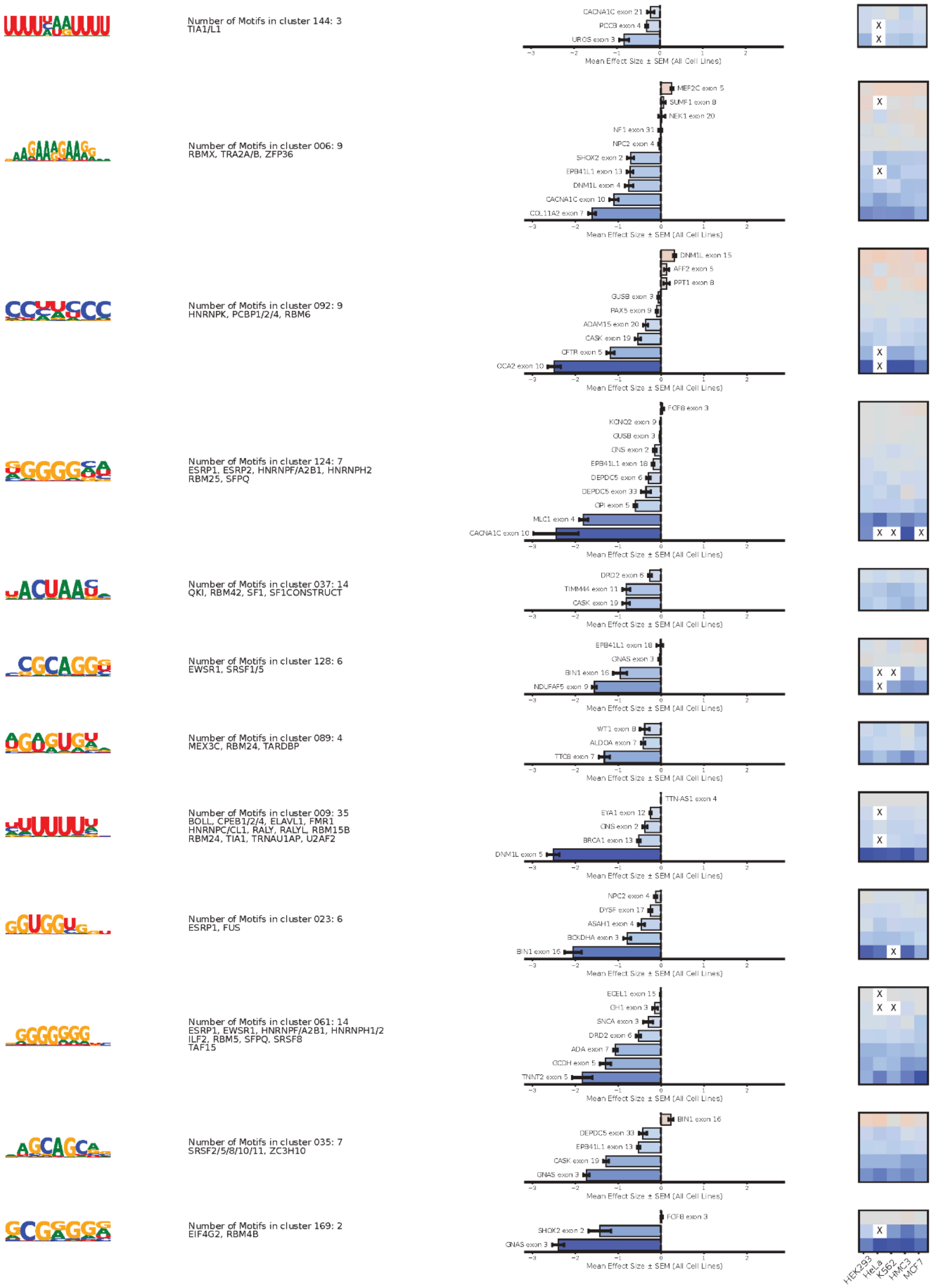

Number of Motifs in cluster 144: 3
TIA1/L1
CACNA1C exon 21
PCCB exon 4
UROS exon 3
Number of Motifs in cluster 006: 9
RBMX, TRA2A/B, ZFP36
MEF2C exon 5
SUMF1 exon 8
NEK1 exon 20
NF1 exon 31
NPC2 exon 4
SHOX2 exon 2
EPB41L1 exon 13
DNM1L exon 4
CACNA1C exon 10
COL11A2 exon 7
Number of Motifs in cluster 092: 9
HNRNPK, PCBP1/2/4, RBM6
DNM1L exon 15
AFF2 exon 5
PPT1 exon 8
GUSB exon 3
PAX5 exon 9
ADAM15 exon 20
CASK exon 19
CFTR exon 5
OCA2 exon 10
Number of Motifs in cluster 124: 7
ESRP1, ESRP2, HNRNPF/A2B1, HNRNPH2
RBM25, SFPQ
FGFB exon 3
KCNQ2 exon 9
GUSB exon 3
GNS exon 2
EPB41L1 exon 18
DEPDC5 exon 6
DEPDC5 exon 33
GPI exon 5
MLC1 exon 4
CACNA1C exon 10
Number of Motifs in cluster 037: 14
QKI, RBM42, SF1, SF1CONSTRUCT
DRD2 exon 6
TIMM44 exon 11
CASK exon 19
Number of Motifs in cluster 128: 6
EWSR1, SRSF1/5
EPB41L1 exon 18
GNAS exon 3
BIN1 exon 16
NDUFAF5 exon 9
Number of Motifs in cluster 089: 4
MEX3C, RBM24, TARDBP
WT1 exon 8
ALDOA exon 7
TTC8 exon 7
Number of Motifs in cluster 009: 35
BOLL, CPEB1/2/4, ELAVL1, FMR1
HNRNPC/CL1, RALY, RALYL, RBM15B
RBM24, TIA1, TRNAU1AP, U2AF2
TTN-AS1 exon 4
EYA1 exon 12
GNS exon 2
BRCA1 exon 13
DNM1L exon 5
Number of Motifs in cluster 023: 6
ESRP1, FUS
NPC2 exon 4
DYSF exon 17
ASAH1 exon 4
BCKDHA exon 3
BIN1 exon 16
Number of Motifs in cluster 061: 14
ESRP1, EWSR1, HNRNPF/A2B1, HNRNPH1/2
ILF2, RBM5, SFPQ, SRSF8
TAF15
ECEL1 exon 15
GH1 exon 3
SNCA exon 3
DRD2 exon 6
ADA exon 7
GCDH exon 5
TNNT2 exon 5
Number of Motifs in cluster 035: 7
SRSF2/5/8/10/11, ZC3H10
BIN1 exon 16
DEPDC5 exon 33
EPB41L1 exon 13
CASK exon 19
GNAS exon 3
Number of Motifs in cluster 169: 2
EIF4G2, RBM4B
FGFB exon 3
SHOX2 exon 2
GNAS exon 3
Mean Effect Size ± SEM (All Cell Lines)
HEK293
HeLa
K562
HMC3
MCF7

**Data S2 | Concordant RBP motifs in exonic regions.**

Motif clusters with consistent effects across exon families are shown. Exonic Splicing Enhancers (ESEs) promote exon inclusion (red), whereas Exonic Splicing Silencers (ESSs) promote exon skipping (blue). The heatmap displays effect sizes for each exon family across individual cell lines, and the accompanying bar plots show mean effect sizes across all cell lines with error bars representing the SEM.
