## Supplementary material for "Massively parallel assay of human splice variants reveals cis-regulatory drivers of disease-associated and cell type-specific splicing regulation": Data S3

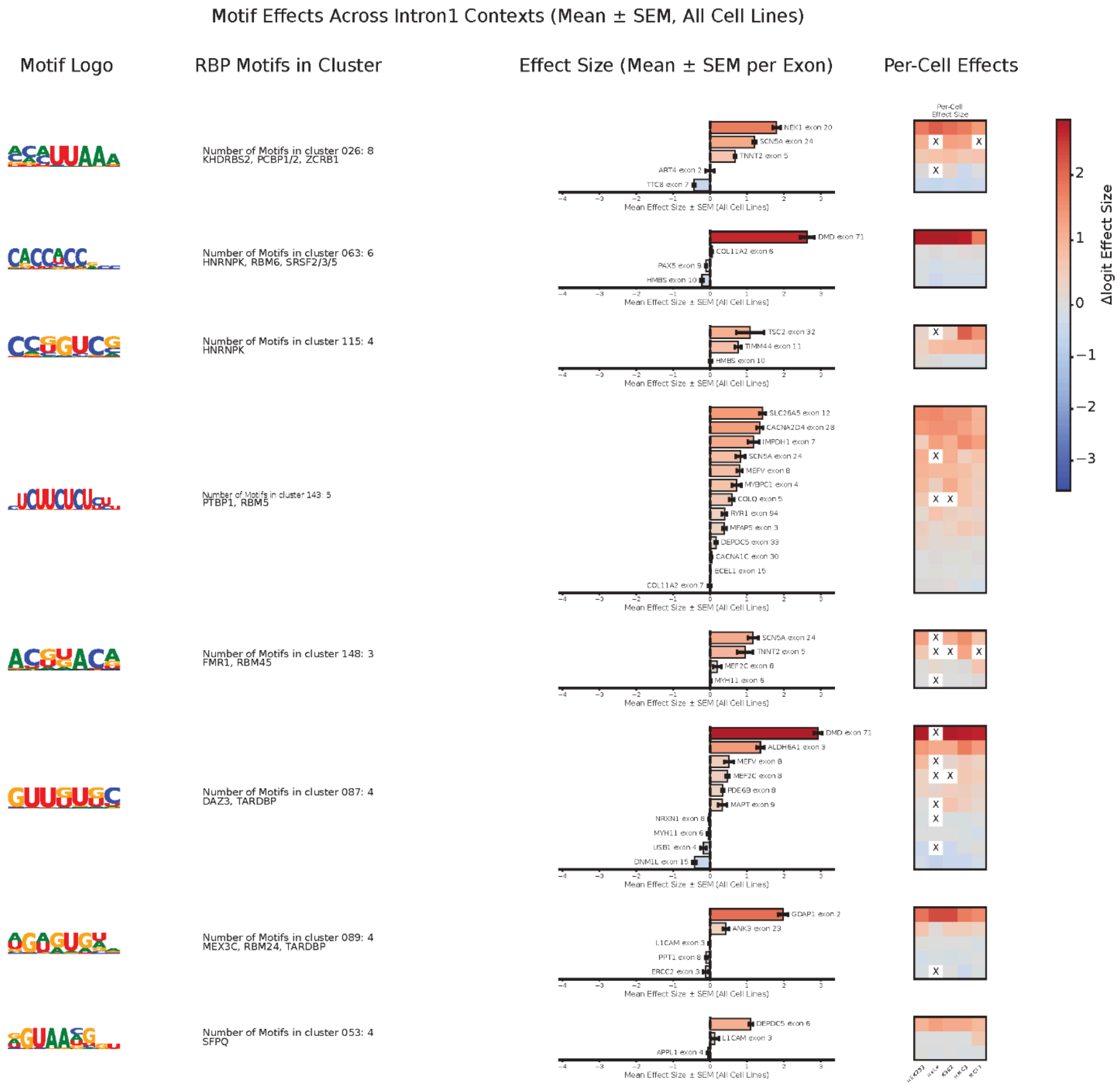

Motif Effects Across Intron1 Contexts (Mean ± SEM, All Cell Lines)
Motif Logo
RBP Motifs in Cluster
Effect Size (Mean ± SEM per Exon)
Per-Cell Effects
Number of Motifs in cluster 026: 8
KHDRBS2, PCBP1/2, ZCRB1
NEK1 exon 20
SCN5A exon 24
TNNT2 exon 5
ART4 exon 2
TTC8 exon 7
Number of Motifs in cluster 063: 6
HNRNPK, RBM6, SRSF2/3/5
DMD exon 71
COL11A2 exon 6
PAX5 exon 9
HMBS exon 10
Number of Motifs in cluster 115: 4
HNRNPK
TSC2 exon 32
TIMM44 exon 11
HMBS exon 10
Number of Motifs in cluster 143: 5
PTBP1, RBM5
SLC26A5 exon 12
CACNA2D4 exon 28
IMPDH1 exon 7
SCN5A exon 24
MEFV exon 8
MYBPC1 exon 4
COLQ exon 5
RYR1 exon 94
MFAP5 exon 3
DEPDC5 exon 33
CACNA1C exon 30
ECEL1 exon 15
COL11A2 exon 7
Number of Motifs in cluster 148: 3
FMR1, RBM45
SCN5A exon 24
TNNT2 exon 5
MEF2C exon 8
MYH11 exon 6
Number of Motifs in cluster 087: 4
DAZ3, TARDBP
DMD exon 71
ALDH6A1 exon 3
MEFV exon 8
MEF2C exon 8
PDE6B exon 8
MAPT exon 9
NRXN1 exon 8
MYH11 exon 6
USB1 exon 4
DNM1L exon 15
Number of Motifs in cluster 089: 4
MEX3C, RBM24, TARDBP
GDAP1 exon 2
ANK3 exon 23
L1CAM exon 3
PPT1 exon 8
ERCC2 exon 3
Number of Motifs in cluster 053: 4
SFPQ
DEPDC5 exon 6
L1CAM exon 3
APPL1 exon 4
Mean Effect Size ± SEM (All Cell Lines)
Per-Cell Effect Size
Δlogit Effect Size

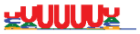

Number of Motifs in cluster 009: 35
BOLL, CPEB1/2/4, ELAVL1, FMR1
HNRNPC/CL1, RALY, RALYL, RBM15B
RBM24, TIA1/L1, TRNAU1AP, U2AF2

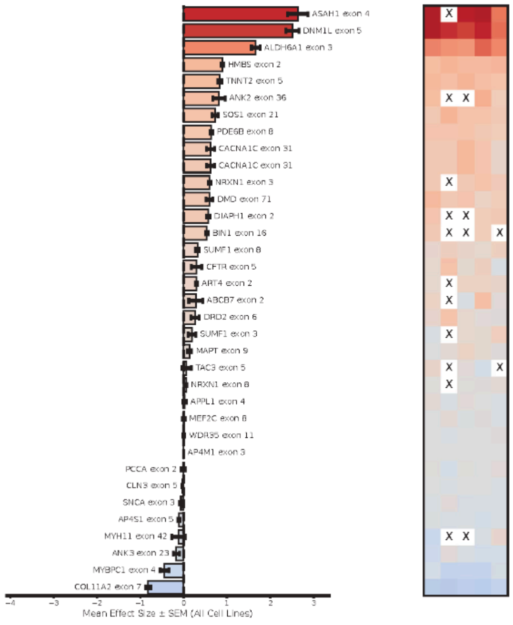


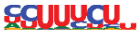

Number of Motifs in cluster 094: 4
PTBP1/3

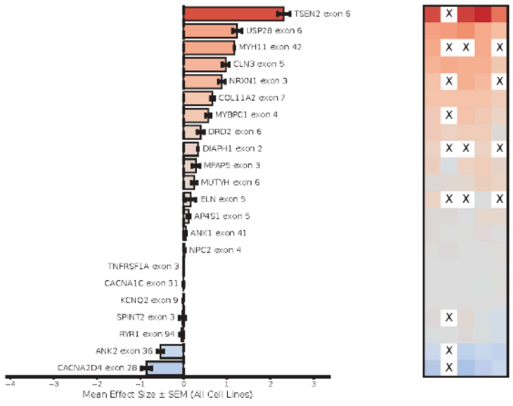


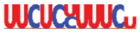

Number of Motifs in cluster 183: 2
PTBP1, TIA1

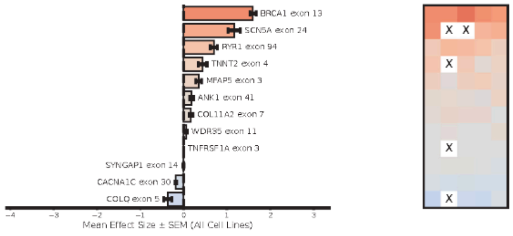


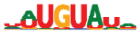

Number of Motifs in cluster 091: 4
FUBP1, KHSRP

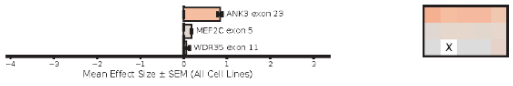


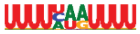

Number of Motifs in cluster 144: 3
TIA1/L1

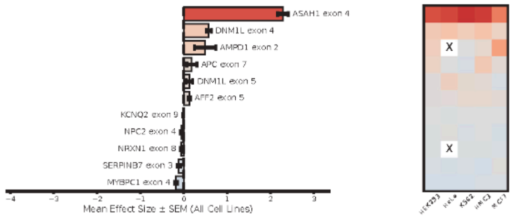

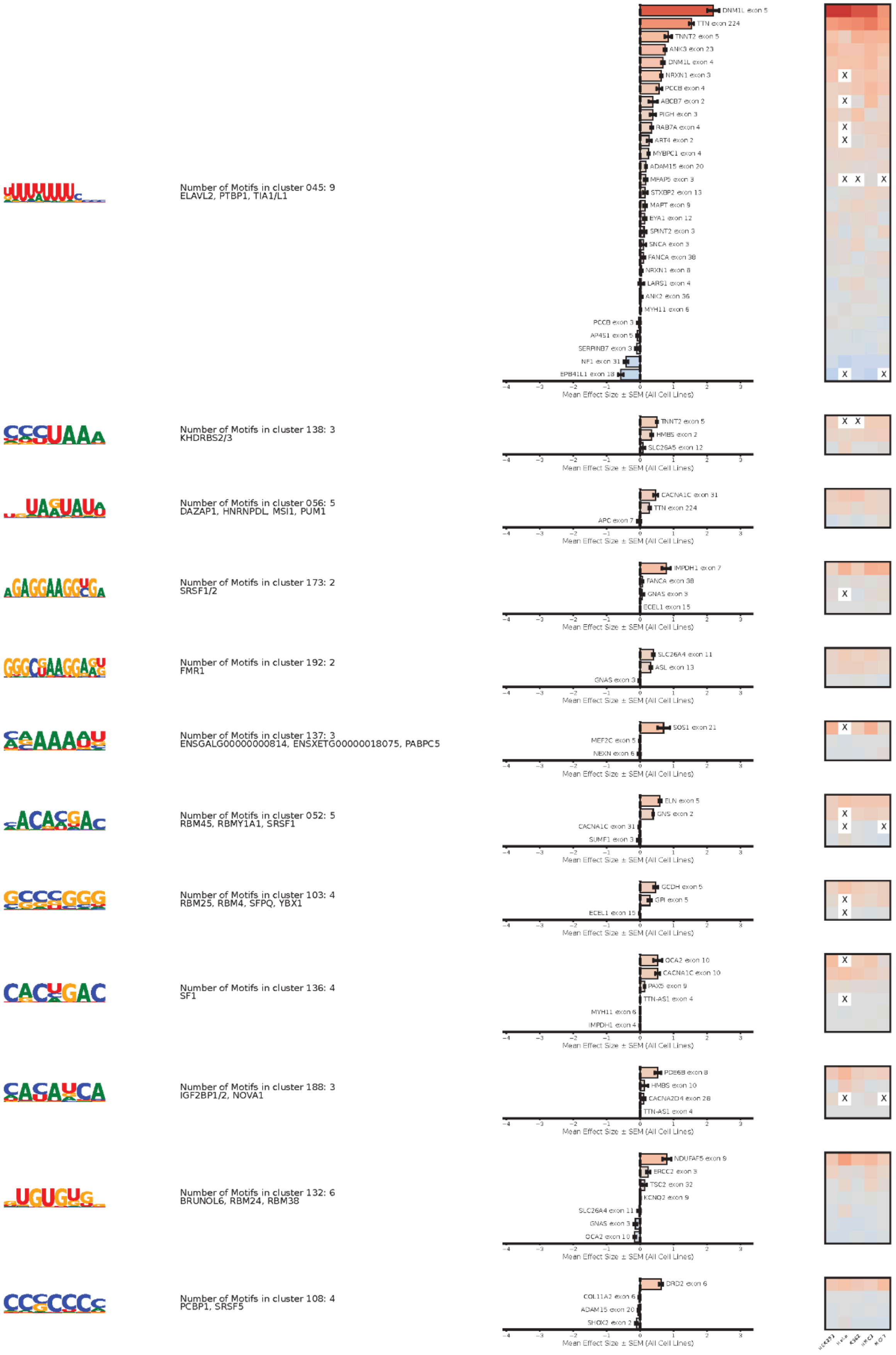

Number of Motifs in cluster 045: 9
ELAVL2, PTBP1, TIA1/L1
DNM1L exon 5
TTN exon 224
TNNT2 exon 5
ANK3 exon 23
DNM1L exon 4
NRXN1 exon 3
PCCB exon 4
ABCB7 exon 2
PIGH exon 3
RAB7A exon 4
ART4 exon 2
MYBPC1 exon 4
ADAM15 exon 20
MFAP5 exon 3
STXBP2 exon 13
MAPT exon 9
EYA1 exon 12
SPINT2 exon 3
SNCA exon 3
FANCA exon 38
NRXN1 exon 8
LARS1 exon 4
ANK2 exon 36
MYH11 exon 6
PCCB exon 3
AP4S1 exon 5
SERPINB7 exon 3
NF1 exon 31
EPB41L1 exon 18
Mean Effect Size ± SEM (All Cell Lines)
Number of Motifs in cluster 138: 3
KHDRBS2/3
TNNT2 exon 5
HMBS exon 2
SLC26A5 exon 12
Number of Motifs in cluster 056: 5
DAZAP1, HNRNPDL, MSI1, PUM1
CACNA1C exon 31
TTN exon 224
APC exon 7
Number of Motifs in cluster 173: 2
SRSF1/2
IMPDH1 exon 7
FANCA exon 38
GNAS exon 3
ECEL1 exon 15
Number of Motifs in cluster 192: 2
FMR1
SLC26A4 exon 11
ASL exon 13
GNAS exon 3
Number of Motifs in cluster 137: 3
ENSGALG00000000814, ENSXETG00000018075, PABPC5
SOS1 exon 21
MEF2C exon 5
NEXN exon 6
Number of Motifs in cluster 052: 5
RBM45, RBMY1A1, SRSF1
ELN exon 5
GNS exon 2
CACNA1C exon 31
SUMF1 exon 3
Number of Motifs in cluster 103: 4
RBM25, RBM4, SFPQ, YBX1
GCDH exon 5
GPI exon 5
ECEL1 exon 15
Number of Motifs in cluster 136: 4
SF1
OCA2 exon 10
CACNA1C exon 10
PAX5 exon 9
TTN-AS1 exon 4
MYH11 exon 6
IMPDH1 exon 4
Number of Motifs in cluster 188: 3
IGF2BP1/2, NOVA1
PDE6B exon 8
HMBS exon 10
CACNA2D4 exon 28
TTN-AS1 exon 4
Number of Motifs in cluster 132: 6
BRUNOL6, RBM24, RBM38
NDUFAF5 exon 9
ERCC2 exon 3
TSC2 exon 32
KCNQ2 exon 9
SLC26A4 exon 11
GNAS exon 3
OCA2 exon 10
Number of Motifs in cluster 108: 4
PCBP1, SRSF5
DRD2 exon 6
COL11A2 exon 6
ADAM15 exon 20
SHOX2 exon 2
Mean Effect Size ± SEM (All Cell Lines)

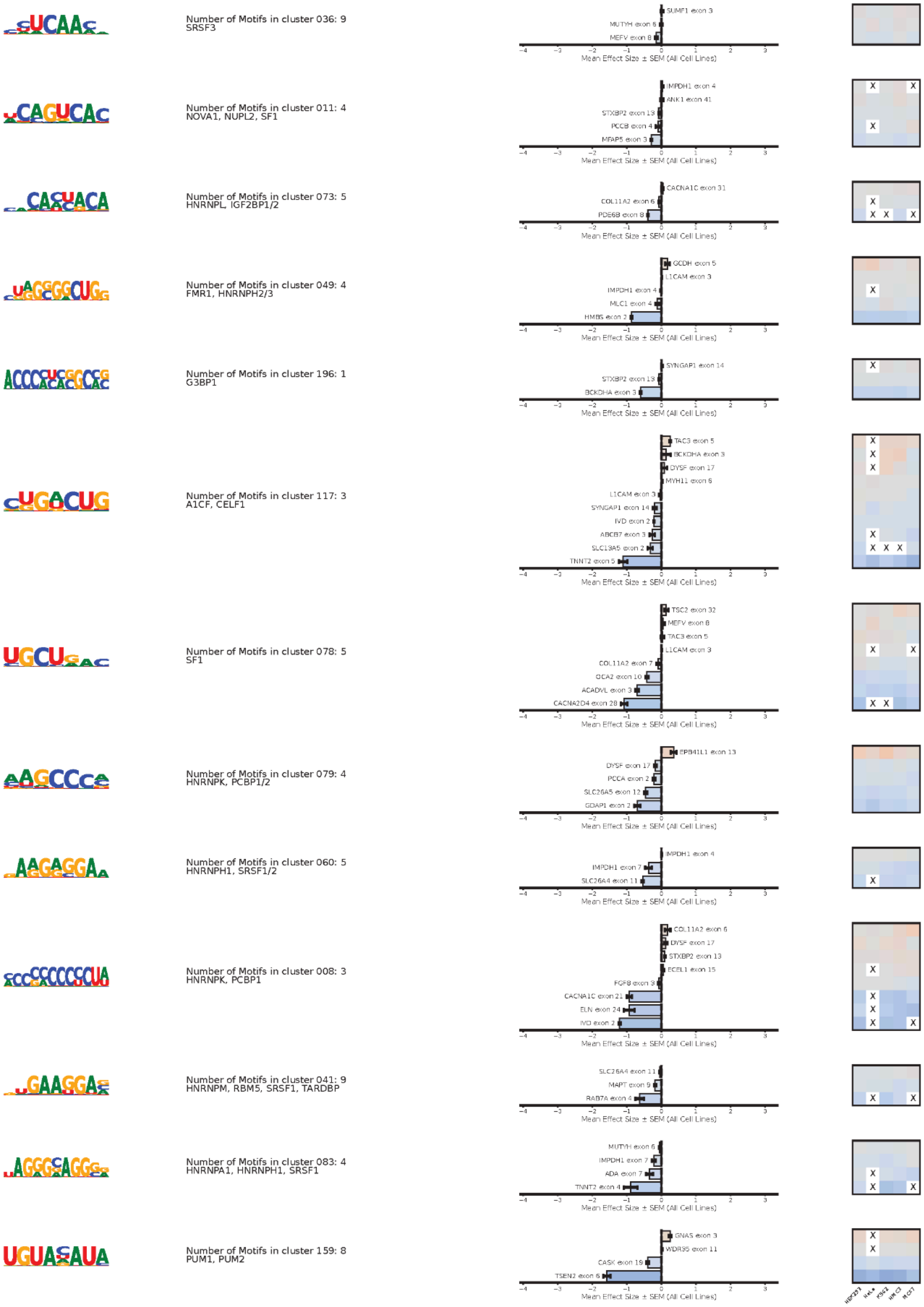

Number of Motifs in cluster 036: 9
SRSF3
SUMF1 exon 3
MUTYH exon 6
MEFV exon 8
Mean Effect Size ± SEM (All Cell Lines)
Number of Motifs in cluster 011: 4
NOVA1, NUPL2, SF1
IMPDH1 exon 4
ANK1 exon 41
STXBP2 exon 13
PCCB exon 4
MFAP5 exon 3
Number of Motifs in cluster 073: 5
HNRNPL, IGF2BP1/2
CACNA1C exon 31
COL11A2 exon 6
PDE6B exon 8
Number of Motifs in cluster 049: 4
FMR1, HNRNPH2/3
GCDH exon 5
L1CAM exon 3
IMPDH1 exon 4
MLC1 exon 4
HMBS exon 2
Number of Motifs in cluster 196: 1
G3BP1
SYNGAP1 exon 14
STXBP2 exon 13
BCKDHA exon 3
Number of Motifs in cluster 117: 3
A1CF, CELF1
TAC3 exon 5
BCKDHA exon 3
DYSF exon 17
MYH11 exon 6
L1CAM exon 3
SYNGAP1 exon 14
IVD exon 2
ABCB7 exon 3
SLC13A5 exon 2
TNNT2 exon 5
Number of Motifs in cluster 078: 5
SF1
TSC2 exon 32
MEFV exon 8
TAC3 exon 5
L1CAM exon 3
COL11A2 exon 7
OCA2 exon 10
ACADVL exon 3
CACNA2D4 exon 28
Number of Motifs in cluster 079: 4
HNRNPK, PCBP1/2
EPB41L1 exon 13
DYSF exon 17
PCCA exon 2
SLC26A5 exon 12
GDAP1 exon 2
Number of Motifs in cluster 060: 5
HNRNPH1, SRSF1/2
IMPDH1 exon 4
IMPDH1 exon 7
SLC26A4 exon 11
Number of Motifs in cluster 008: 3
HNRNPK, PCBP1
COL11A2 exon 6
DYSF exon 17
STXBP2 exon 13
ECEL1 exon 15
FGF8 exon 3
CACNA1C exon 21
ELN exon 24
IVD exon 2
Number of Motifs in cluster 041: 9
HNRNPM, RBM5, SRSF1, TARDBP
SLC26A4 exon 11
MAPT exon 9
RAB7A exon 4
Number of Motifs in cluster 083: 4
HNRNPA1, HNRNPH1, SRSF1
MUTYH exon 6
IMPDH1 exon 7
ADA exon 7
TNNT2 exon 4
Number of Motifs in cluster 159: 8
PUM1, PUM2
GNAS exon 3
WDR35 exon 11
CASK exon 19
TSEN2 exon 6
Mean Effect Size ± SEM (All Cell Lines)
HEK293
HeLa
K562
HMC3
MCF7

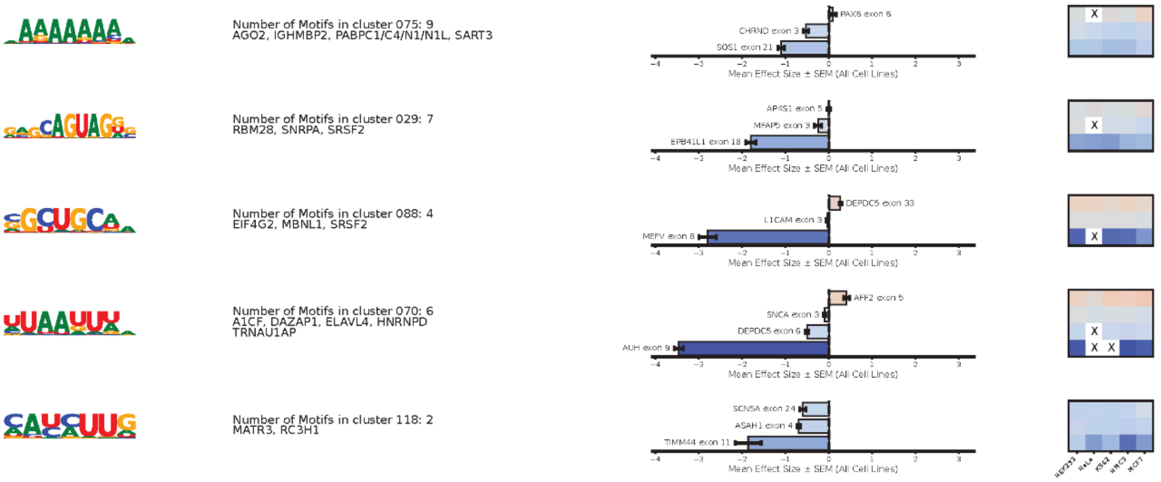
Number of Motifs in cluster 075: 9
AGO2, IGHMBP2, PABPC1/C4/N1/N1L, SART3
PAX6 exon 6
CHRND exon 3
SOS1 exon 21
Mean Effect Size ± SEM (All Cell Lines)
X
Number of Motifs in cluster 029: 7
RBM28, SNRPA, SRSF2
AP4S1 exon 5
MFAP5 exon 3
EPB41L1 exon 18
Mean Effect Size ± SEM (All Cell Lines)
X
Number of Motifs in cluster 088: 4
EIF4G2, MBNL1, SRSF2
DEPDC5 exon 33
L1CAM exon 3
MEFV exon 8
Mean Effect Size ± SEM (All Cell Lines)
X
Number of Motifs in cluster 070: 6
A1CF, DAZAP1, ELAVL4, HNRNPD
TRNAU1AP
AFF2 exon 5
SNCA exon 3
DEPDC5 exon 6
AUH exon 9
Mean Effect Size ± SEM (All Cell Lines)
X
X X
Number of Motifs in cluster 118: 2
MATR3, RC3H1
SCN5A exon 24
ASAH1 exon 4
TIMM44 exon 11
Mean Effect Size ± SEM (All Cell Lines)
HEK293
HeLa
K562
MCF7

**Data S3 | Concordant RBP motifs in intronic regions.**

Motif clusters found in the 5′ intron with consistent effects across exon families are shown. Intronic Splicing Enhancers (ISEs) promote exon inclusion (red), whereas Intronic Splicing Silencers (ISSs) promote exon skipping (blue). The heatmap displays effect sizes for each exon family across individual cell lines, and the accompanying bar plots show mean effect sizes across all cell lines with error bars representing the SEM.
